# A metric for comparing complex systems by their dynamics

**DOI:** 10.64898/2026.07.16.738953

**Authors:** Mitchell Ostrow, Adam J. Eisen, Leo Kozachkov, William T. Redman, Ila Fiete

## Abstract

Comparisons are fundamental to science: experiment against model, one organism against another, a system against itself across time. Because many systems, from brains to climate, are characterized by how they evolve in time, it is a natural goal to compare their dynamics. Dynamical systems comparison is well defined, but has been intractable for nonlinear, high-dimensional, noisy, and partially observed data. As a result, standard comparison methods have focused on the geometry or topology of data. Here we present Dynamical Similarity Analysis (DSA), a class of metrics to compare systems by their temporal evolution. Its foundation is Koopman Operator theory, which recasts nonlinear systems as linear operators. We estimate these operators from data, then compare the operators across systems. The computation is fast, scalable, and robust to noise and partial observation. It is also differentiable. DSA identifies dynamical structure that geometric and topological methods miss. It matches recordings from the head direction circuit to ring attractor models. It shows that macaque motor cortex dynamics for two reaching tasks drift apart across years despite preserved behavior, and that primary motor cortex breaks from premotor cortex as movement begins. As an optimization objective, it induces neural networks to learn never-before hypothesized solutions that run counter to their inductive biases. Thus, DSA transforms the dynamics of a system into an object that can be measured, compared, and optimized.

## Introduction

Empirical science rests on comparison: between experiments, between model and data, across conditions, across time. For systems well-characterized by a static set of states, such as sets of images, gene expression vectors, and activations in feedforward or equilibrium neural networks, a mature toolkit exists to characterize the geometric and topological structure of the states (*Pearson, 1901; Tenenbaum et al., 2000; van der Maaten and Hinton, 2008; Roweis and Saul, 2000; Belkin and Niyogi, 2003; Ghrist, 2008*) and measure similarity (*Williams et al., 2021; Kornblith et al., 2019; Kriegeskorte et al., 2008; Raghu et al., 2017*, although some of these methods are not proper metrics; *Kriegeskorte et al., 2008; Schrimpf et al., 2020*). However, the climate, ecosystems, financial markets, learning algorithms, most brain circuits, and many natural systems are defined by how they change over time. For these systems a key notion of similarity is not the geometry or topology of states but rather the dynamics that generate them.

Comparing dynamics is crucial in neuroscience, where brain circuits generate ring attractor dynamics to construct an updating compass estimate during navigation (*Taube, 1995; Zhang, 1996; Chaudhuri et al., 2019; Kim et al., 2017*), line attractor dynamics for evidence integration (*Seung, 1996; Khona and Fiete, 2022*), and trajectories for speech generation and comprehension (*Anumanchipalli et al., 2019; Jamali et al., 2024*) as well as movement execution (*Churchland et al., 2012*). Validating circuit models with neural recordings, or comparing neural dynamics across animals, days, brain regions, and behavioral contexts, requires asking whether the systems realize equivalent dynamics. This need arises in any discipline where time-series data is collected at scale (Fig. 1a).

**Figure 1.**
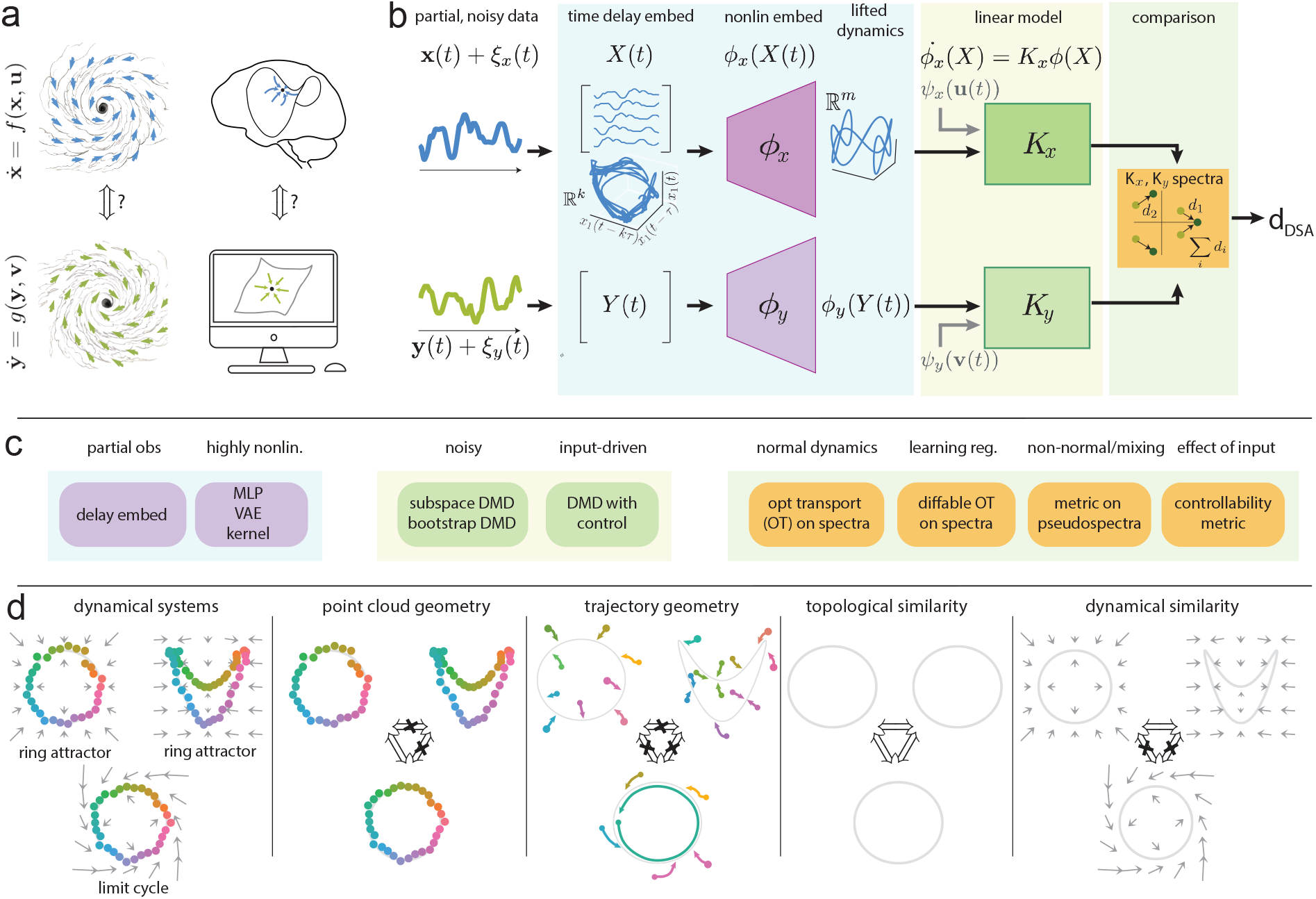
DSA for quantitatively comparing dynamics between systems. **(a)** We frequently seek to evaluate the similarity of two systems, e.g. two hurricanes (first column), or the dynamics of a brain circuit and that of a model (second column). Formally, two systems are conjugate or dynamically equivalent if their vector fields can be exactly aligned by a smooth transformation. **(b)** DSA optimally aligns dynamics and quantitatively measures the degree of alignment to generate a dynamical distance measure in a generic, modular pipeline that can be applied to any time-varying processes: Observed data (states **x, y** or their noisy counterparts) are delay embedded into a higher-dimensional space, then optionally additionally nonlinearly embedded (with a nonlinear kernel, neural network, or other methods). Linear dynamics models *K*_*x*_, *K*_*y*_ are then fit (with cross-validation) to the embedded data together with potential system inputs or condition data *u, v* to predict a future timestep. Some summary statistic (e.g. the eigenvalue spectrum) is aligned between models, using optimal transport or a related process. The DSA distance is then generated based on the quantitative match of the models’ summary statistic. **(c)** Examples of modular DSA components for different settings: (left) nonlinear embeddings, (middle) various linear modeling frameworks based on Dynamic Mode Decomposition (DMD), (right) metrics for comparison of the linear models’ summary statistic. **(d)** Complementarity of similarity measures: (first panel) three example dynamical systems; (second to fifth panels): similarity estimates of point-cloud geometric comparison methods (second panel); methods focused on the geometry of observed trajectories (third panel); methods focused on topological structure (fourth panel); DSA (fifth panel).

Existing similarity methods address complementary needs. Geometric methods (e.g. Procrustes analysis; *Schönemann, 1966; Williams et al., 2021*; Representational Similarity Analysis; *Shepard and Chipman, 1970; Kriegeskorte et al., 2008*; Canonical Correlation Analysis; *Hotelling, 1936; Raghu et al., 2017; Gallego et al., 2020*) compare the shape of point clouds in state space and are agnostic to the temporal order in which points are visited. Topological methods (*Ghrist, 2008; Curto and Sanderson, 2025; Chaudhuri et al., 2019; Gardner et al., 2022*) abstract beyond geometry to compare the connectivity of the data manifold, but likewise ignore temporal information. Recent attempts at direct dynamical comparison (*Gosztolai et al., 2025; Chen et al., 2024*) train artificial neural networks to compare systems, an approach that scales poorly when the relevant analysis involves hundreds or thousands of pairwise distances on massive datasets (SI F.1). Notably, as far as we are aware, no prior method can handle the effect of time-varying input to the relevant systems. A scalable, theoretically grounded metric on dynamics, computable from raw time-series data, has been missing.

Here we present Dynamical Similarity Analysis (DSA), a theoretically driven, scalable method to directly compare dynamics in time-series data (Fig. 1b). DSA is built on Koopman Operator theory (*Koopman, 1931; Mezić, 2005; Brunton et al., 2022; Budišić et al., 2012*), which states that any well-behaved nonlinear dynamical system can be represented exactly as a linear operator acting on an infinite-dimensional space of functions of the state. DSA uses approaches such as the Extended Dynamic Mode Decomposition (eDMD, *Schmid, 2022; Williams et al., 2015*) to construct finite-dimensional approximations of these operators, using delay embeddings (*Takens, 1981; Brunton et al., 2017; ArbabiandMezić, 2017*) and nonlinear feature embeddings (*Williams et al., 2016; Lusch et al., 2018*) to deal with noisy and partially observed data. These model-agnostic methods have been applied to diverse data domains, such as neuroscience (*Brunton et al., 2016, 2017*), fluid dynamics (*Schmid, 2010; Krake et al., 2021*), behavior (*Fujii et al., 2020; Costa et al., 2019, 2024*), finance (*Mann and Kutz, 2016*), infectious disease (*Proctor and Eckhoff, 2015*), and more, making them a strong foundation on which to build robust similarity metrics. DSA then computes matrix distances on these data-driven operators or their summary statistics to obtain a dynamical distance measure. DSA is modular, and each step in the pipeline is fast, parallelizable, customizable, and can be made differentiable end-to-end (Fig. 1c).

Components of DSA were recently developed across a series of focused studies in conference proceedings, each establishing one piece in a specialized setting: the original formulation with a Procrustes metric on vector fields (*Ostrow et al., 2023*), the optimal-transport distance on Koopman spectra (*Redman et al., 2022, 2024*), and the extension to input-driven systems (*Huang et al., 2025*). Here we unify these advances into a single coherent and general framework that exposes the shared theoretical basis underlying methods that were introduced separately, with prior methods reducing to specific cases, and present improvements that make DSA modular, easily extensible to new variants, and significantly more efficient and accurate. We then demonstrate DSA across problems that no single prior study could address—spanning synthetic benchmarks, direct brain-to-model comparison, multi-region and multi-timescale neural dynamics, and the use of dynamical distance as a training objective. Overall, we will show that DSA is a general method that can be applied whenever dynamical time-series data are collected, providing a principled, scalable approach to the problem of measuring dynamical similarity.

### Theoretical framework: Comparing dynamical systems via Koopman Operators

Two dynamical systems, described by 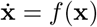 and 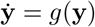 with states **x**(*t*) ∈ ℝ^*n*^ and **y**(*t*) ∈ ℝ^*m*^ (Fig. 1a), are mathematically equivalent in the strongest sense when one can be smoothly warped into the other—formally, when there is a smooth invertible map Φ (with Jacobian *D*Φ) between their state spaces such that *g* о Φ = *D*Φ · *f*. Conjugate flow maps over some time *t g*^*t*^, *f* ^*t*^, are related via *g* о Φ = Φ о *f*. This condition, known as *smooth topological conjugacy*, guarantees that the systems share all of their qualitative dynamical features, including fixed points, limit cycles, invariant manifolds, and basins of attraction, as well as quantitative invariants like Lyapunov exponents. To compare systems by degree, one can ask how close they come to being conjugate:

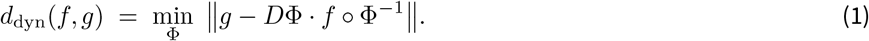

However, evaluating Eq. (1) directly is typically intractable, requiring optimization over an infinite-dimensional space of smooth maps Φ between two possibly high-dimensional state spaces, and the definition of norms over high dimensional function spaces is likewise typically intractable for a given Φ (*Skufca and Bollt, 2008*). To make matters worse, experimental systems are usually partially observed and underlying dynamics *f* and *g* are not explicitly known. DSA is motivated by these problems. We approach the conjugacy problem by relaxing it to a tractable computation (Fig. 1b). First, we leverage Takens’ theorem (*Takens, 1981*) to estimate the full latent states from partial observations by stacking time-delayed copies of the measurements into a delay-embedding matrix,

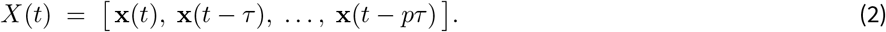

Second, we solve the challenge of finding a conjugacy map (Fig. 1b) by leveraging the fact that conjugacy maps between linear dynamical systems are themselves linear and applying Koopman Operator theory to justify a globally linear description of the nonlinear system. Koopman operator theory (*Koopman, 1931; Mezić, 2005; Brunton et al., 2022; Budišić et al., 2012*) guarantees that any (sufficiently regular) nonlinear system 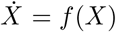 becomes exactly and globally linear with appropriate embeddings into a Hilbert space by scalar functions *ψ*(*x*):

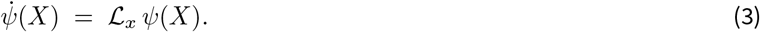

Here ℒ_*x*_ is the Koopman generator that linearly propagates the dynamics, at the cost of acting on the infinite-dimensional space of functions *ψ* rather than on the finite-dimensional state *X*. In discrete time, we convert ℒ_*x*_ into the corresponding discretetime operator *K*_*x*,Δ*t*_ = exp(Δ*t* ℒ_*x*_), so that Eq. (3) becomes *ψ*(*X*_*t*+Δ*t*_) = *K*_*x*,Δ*t*_ *ψ*(*X*_*t*_). A central result of Koopman theory (when the associated Koopman Operators do not have continuous spectra) is that two conjugate systems (related by a smooth invertible Φ in Eq. (1)) have identical Koopman eigenvalues (*Budišić et al., 2012*). As a corollary, mismatched eigenvalues imply that no such Φ exists. Thus, the Koopman framework approximates the conjugacy assessment via the efficient computation and comparison of the eigenvalues of the Koopman Operator. Other metrics on the linear operators are also easy to compute and can be used to compare different aspects of dynamics. This is the conceptual core of DSA.

We approximate the infinite-dimensional Koopman Operator *K*_*x*,Δ*t*_ by a finite-dimensional matrix *K*_*x*,Δ*t*_ that acts on a nonlinear feature embedding of the data *X*. We find a nonlinear *d*-dimensional feature embedding *ϕ*_*x*_ that approximately linearizes dynamics in the feature space:

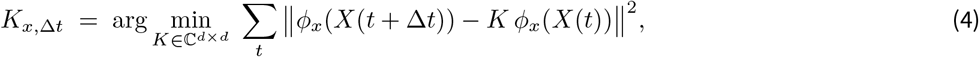

where formally, the goal is to make span(*ϕ*_*x*_) approximately closed under *K*_*x*_. *K*_*y*_ is defined analogously from *Y* (*t*) via embedding *ϕ*_*y*_. The embeddings *ϕ*_*x*_, *ϕ*_*y*_ may be kernel feature maps, polynomial bases, or neural networks (*Gilpin, 2020; Pandarinath et al., 2018; Tenenbaum et al., 2000; Williams et al., 2016; Lusch et al., 2018; Schneider et al., 2023*) and may be chosen ahead of time or learned (*Li et al., 2017; Lusch et al., 2018*). Eq. (4) is the extended dynamic mode decomposition (eDMD) (*Schmid, 2022; Williams et al., 2015*). Delay embeddings, used above to mitigate partial observability, themselves also serve as a simple and efficient nonlinear dimensionality expanding embedding for nonlinear systems (*Arbabi and Mezić, 2017; Brunton et al., 2017*). For further details on how the operator is learned, see SI F.2, F.3.

Likewise, regression variants exist that best suit different settings, such as estimating the operator efficiently under noise or obtaining confidence intervals on its entries (*Pan et al., 2023; Ichinaga et al., 2024*). When the dynamics are driven by time-varying inputs **u**(*t*), **v**(*t*) (which could be known, measured, or designed), input effects can be estimated by augmenting Eq. (4) with input operators *B*_*x*_, *B*_*y*_ that project the (possibly separately nonlinearly-embedded) inputs into the embedding space, and jointly fitting the intrinsic and input-driven dynamics. This class of eDMD models is called DMD with control (*Proctor et al., 2016; Huang et al., 2025*):

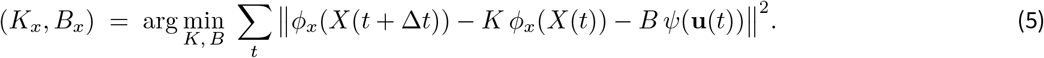

Third, given learned operators *K*_*x*_ and *K*_*y*_, we measure their dissimilarity by

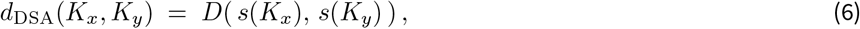

where *s*(·) is some statistic of the matrix that is invariant to coordinate choice—its spectrum, pseudospectrum, or a related object—and *D* is a distance metric or pseudometric between such statistics (SI F.4). By virtue of its linearity, the Koopman Operator admits various well-defined similarity metrics that are informative of different features of the dynamics. For the case of input-driven systems, the similarity function has the form *d*_DSA_(*K*_*x*_, *K*_*y*_, *B*_*x*_, *B*_*y*_); the downstream comparison metric is again modular; using a metric based on controllability would be referred to as InputDSA (SI F.4.4 *Huang et al., 2025*).

In the examples that follow, we typically use the eigenspectra ***λ***_*x*_, ***λ***_*y*_ ∈ *ℂ*^d^ of *K*_*x*_ and *K*_*y*_ and set *D* to be the 2-Wasserstein distance. This eigenvalue distance is a metric and most informative for systems dominated by discrete eigenvalues, such as fixed point dynamics, limit cycles, quasiperiodic motion, and attractor dynamics. For general non-normal operators or systems with a continuous spectral component (strongly mixing or chaotic regimes), distinct operators can be isospectral, so the distance is a pseudometric. There, we may further compare *pseudospectra* (*Colbrook et al., 2023; Colbrook and Townsend, 2024; Trefethen and Embree, 2005*) of *K*_*x*_, *K*_*y*_ or directly compare *K*_*x*_, *K*_*y*_ with the Procrustes alignment on vector fields (PAVF) measure (*Ostrow et al., 2023*) to capture the transient, non-normal structure that eigenspectra might miss (SI F.4). If the underlying state space is equivalent across systems, then the eigenvectors (Koopman modes) are also comparable across systems, motivating Grassmannian distances on eigenspaces (*Cohen et al., 2023; Germain et al., 2026*, SI F.4).

Each component of DSA can be computed quickly, enabling tens to hundreds of thousands of pairwise comparisons on the order of seconds. Further, DSA can be made differentiable in *ϕ*_*x*_, *ϕ*_*y*_ using an entropic regularization of the Wasserstein metric (*Cuturi, 2013*), so that it can serve as an optimization objective, as we will show below.

Given a DSA distance, what aspects of dynamical systems would it differentiate? Conceptually, DSA captures similarities in the spatiotemporal evolution of two systems based on their full dynamical operators (thus flow-fields). Consider three different dynamical systems (schematized in Fig. 1d), one consisting of a continuum of stable fixed points in a circular ring of uniform curvature (circular ring attractor), one that is a ring of fixed points with nonuniform curvature (crumpled ring attractor), and one that is a circular limit cycle. Measures that treat the data as an unordered set and compare the geometry of states (e.g. Maximum-Mean Discrepancy (MMD (*Gretton et al., 2012*)) or Wasserstein Procrustes (WP (*Grave et al., 2019*))) would identify the circular ring attractor with the circular limit cycle and distinguish both from the crumpled ring attractor. Measures comparing the geometry of the temporally-ordered trajectories would consider all three to be different (e.g. Canonical Correlation Analysis (CCA (*Gallego et al., 2018; Safaie et al., 2023*)), Centered Kernel Alignment (CKA (*Kornblith et al., 2019*)) or Procrustes analysis (PA (*Williams et al., 2021*))), while measures based on the connectivity of the point clouds via Topological Data Analysis (TDA (*Zomorodian and Carlsson, 2004; Chaudhuri et al., 2019; Wasserman, 2018*)) would identify all three as equivalent. DSA, by contrast, would compare the underlying vector fields, identifying the two ring attractors as similar and both as different from the limit cycle. Note that all of these metrics produce useful insight; DSA’s role is to complement prior methods with insight into temporal similarity.

## Results

### DSA captures dynamical similarity across diverse systems with known ground truth

We begin by assessing DSA on complex dynamical systems consisting of various kinds of fixed point, limit cycle, and chaotic attractors, for which we know the ground truth (Methods B).

We considered four types of models that generate continuous attractors with the topology of a ring and two types of limit cycle models (Fig. 2a,b; Methods B.1). The first is a recurrent neural network (*Xie et al., 2002; Goodridge and Touretzky, 2000*) model of the brain’s compass circuit for continuously tracking heading direction as animals navigate in space (*Taube et al., 1990; Taube, 1995; Peyrache et al., 2015; Turner-Evans et al., 2017*) (NN ring, Fig. 2a, top). Though the states exist in a high-dimensional space, the system generates a continuous set of fixed points that form a one-dimensional ring, such that perturbations away from the ring rapidly decay to it. When driven by angular velocity inputs, the state shifts along the ring, and in the absence of velocity inputs, the state along the ring remains stable. The second model is a simple 2-variable dynamical system (simple ring, Fig. 2b, top) that also implements a ring attractor. Each ring was driven with angular velocities sampled from an Ornstein-Uhlenbeck process (Methods B.1). The third and fourth models are versions of the first and second, respectively, but we drive them with a constant angular velocity input (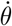 in Fig. 2a,b, top), converting their dynamics into limit cycles. Additionally, we construct cut ring models, where we introduce a cut in the ring structure of each ring model, generating a curved one-dimensional line attractor manifold (cut rings, Fig. 2a,b, bottom). Like the ring models, the cut ring models were driven with an angular velocity sampled from an Ornstein-Uhlenbeck process (Methods B.1); however, states cannot traverse the gap in the ring, thus the onring dynamics of the cut rings are globally distinct from that of the rings.

**Figure 2.**
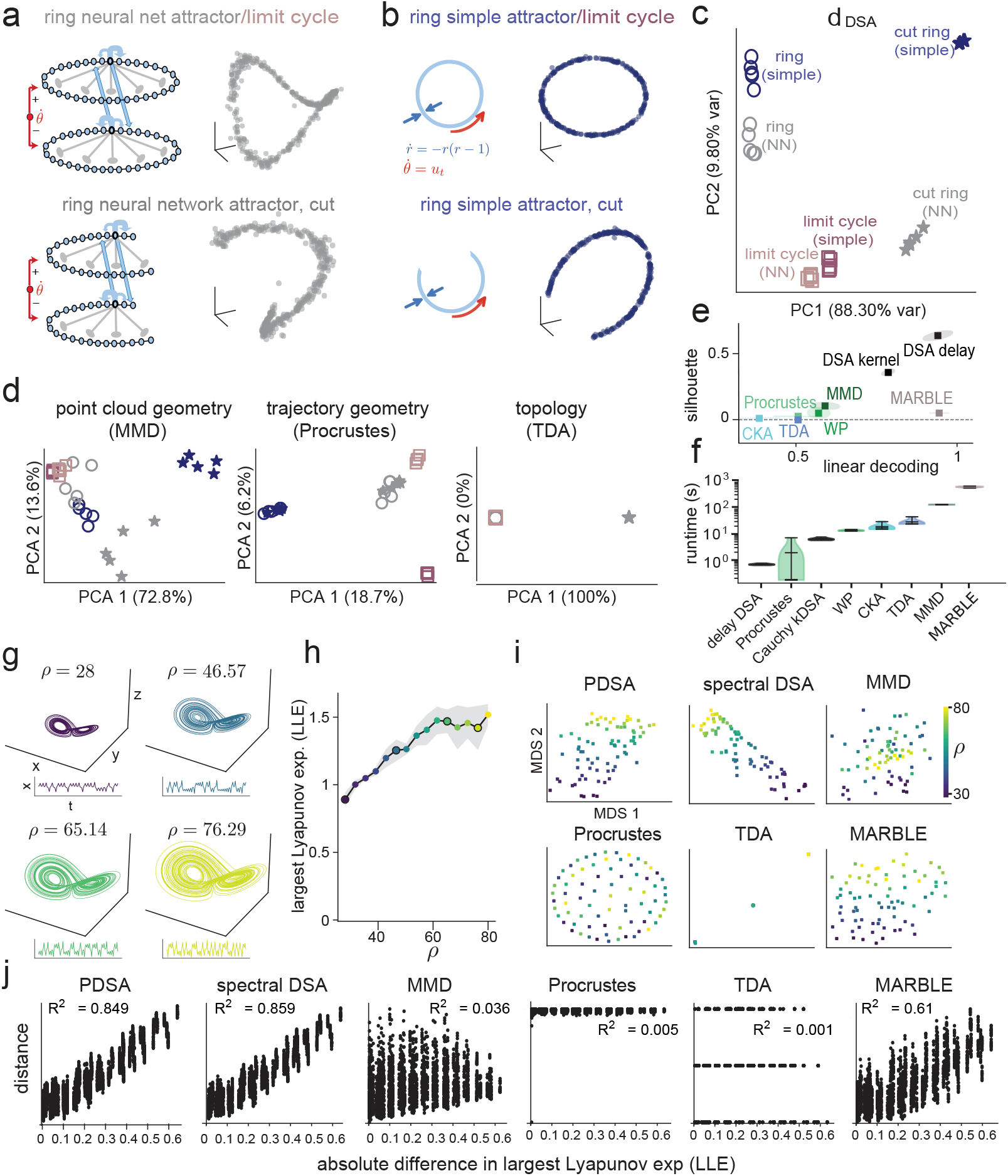
DSA accurately captures similarity between complex dynamical models with known ground truth. **(a)** (top) Schematic of the neural ring attractor, which is parameterized as a ring of neurons that are locally exciting and globally inhibiting. On the bottom, the periodic boundary conditions are broken, giving rise to a line (cut ring) attractor. (Right) scatterplot of observations, projected into the top 3 principal components. The ring and limit cycle models have matched manifold structures, but on-manifold dynamics on the ring models follow an OU process while in the limit cycle models they correspond to unidirectional fixed-speed rotations. **(b)** Similar to (a) but for the simple ring attractor, which is directly parameterized as a two dimensional polar system of differential equations. The cut ring attractor is a broken form of the ring attractor, which induces different dynamics (no pass-through of trajectories through the cut). (Right) scatterplot of observations. **(c)** DSA distances between simulated models, distance matrix projected to the top two PCs. DSA clusters ring models together, cut rings together, and limit cycles together (across simple and NN versions of each), showing an emphasis on dynamics over geometry. **(d)** PCA visualization of baseline comparison methods for point cloud geometry (MMD, left), trajectory geometry (Procrustes, middle), and topology (TDA, right). **(e)** Linear decoding validation accuracy and silhouette score for each comparison method using the dynamics class labels. Scores computed across 30 synthetic systems (3 dynamics *×* 2 model classes *×* 5 random initializations); ellipses indicate *±* 1 s.d. over 10 bootstrap resamples (Methods Sec. B.2). **(f)** Average runtime per iteration of the comparison methods from (e). **(g)** Trajectories of Lorenz attractor models for 4 settings of the parameter *ρ*, with a sample of *x*(*t*) below. All display a canonical butterfly attractor whose geometries and trajectories differ. **(h)** Largest Lyapunov Exponent computed along 5 randomly initialized trajectories for each *ρ* parameter. Shaded region denotes standard deviation, circled points indicate parameters visualized in (g). **(i)** Different comparison measures applied to these models (5 random initializations for each of 15 *ρ* values), with each 75 *×* 75 matrix reduced with MDS and visualized as a scatterplot, colored by *ρ* value. PDSA utilizes the Procrustes Analysis over Vector Fields metric (PAVF, SI F.2), while spectral DSA utilizes the default Wasserstein distance. **(j)** Correlation between the distances estimated by the various comparison methods across models and the absolute difference in LLE estimates between models.

We sampled data from the models without trial structure, as there are no events onto which trajectories can be time-aligned. Here, geometric comparison methods that require such alignment cannot practically be applied. Point-cloud comparison methods do not require temporal alignment, however they neglect the temporal ordering necessary to compare dynamics.

We first examined the ability of DSA and other methods to distinguish different variations of ring attractor models (simple rings, neural network rings, and cut and limit cycle versions), where the ground truth labels are known. DSA clearly distinguished the model ring and cut ring attractors from each other (Fig. 2c). This was true even when the cut rings were simulated with arbitrarily small cuts such that their point cloud geometry was indistinguishable from the uncut versions (Fig. S1). DSA’s discrimination of cut and uncut rings arises because the cut induced different dynamics and therefore distinct Koopman eigenspectra along the ring, with trajectories unable to traverse the cut (analytically derived eigenspectra for the models in SI G). For similar reasons, DSA is able to separate limit cycle dynamics on the manifold from non-limit cycle dynamics on the same manifold (Fig. 2c). DSA also separates simple and neural network ring models, indicating that it also has some sensitivity to geometry; however, the primary distinctions it makes are by dynamics rather than shape, as the separation between rings and cut rings and between rings and limit cycles is much greater than that between simple and neural network rings.

Existing methods, as expected, have much varying and generally lower degrees of success in comparing these models (Fig. 2d,e, Fig. S1, Fig. S2, Methods B.3). These methods include point cloud geometry comparisons with Maximum Mean Discrepancy (MMD) (*Gretton et al., 2012*) and Wasserstein Procrustes (WP) (*Grave et al., 2019*); trajectory geometry compared with Procrustes and Nonlinear CKA (*Kornblith et al., 2019*) (done by placing the datasets in correspondence by their shared timepoints, so each metric compares ordered trajectories rather than unordered point sets), and topology comparison with TDA (*Zomorodian and Carlsson, 2004; Wasserman, 2018*). In addition, we implemented dynamical comparison using a leading neural network technique for embedding dynamics, MARBLE (*Gosztolai et al., 2025*), which likewise was unable to distinguish the rings and cut rings when the cuts became small (Fig. S1b).

To quantify performance across methods, we applied linear decoders and computed the silhouette score (a clustering metric) on ring, cut-ring, and limit cycle labels. Linear decoding accuracy quantifies separability, while the silhouette score quantifies variance within classes relative to across classes. DSA computed with delay embeddings was the most effective method (perfect linear decoding accuracy and average silhouette score of 0.67 across 10 bootstraps), while DSA with a Cauchy kernel DMD was close behind (Fig. 2e). Most distance metrics had a linear decoding accuracy only slightly greater than chance and silhouette scores effectively at zero (Fig. 2e). MARBLE linearly separated the data by dynamics type but did not cluster them well, with a silhouette score near zero (Fig. 2e). Only DSA and TDA reported significantly greater distances between models of different topologies (rings versus cut rings) than between models of the same topology (Fig. S2a).

Further, when we timed the methods on the same data in a standardized computing environment, DSA had a faster wall-clock time than every method except for trajectory geometry comparison with Procrustes (Fig. 2f). MARBLE, being a neural network method, was roughly 1000× slower than DSA.

We next evaluated DSA on the chaotic Lorenz attractor, sweeping the *ρ* parameter at 15 equally-spaced values in the chaotic regime from 28 to 80 (Methods B.4), and comparing the resulting systems with each other. This parameter dials the degree of chaos in the system (as measured by the largest Lyapunov exponent), and reshapes the well-known chaotic butterfly attractor structure (Fig. 2g,h).

Because the Lorenz attractor is chaotic and has a continuous Koopman spectrum (*Arbabi and Mezić, 2017*), we hypothesized that pseudospectral information or comparisons on the full learned linear operators would be critical, beyond comparisons of the operator spectra. We therefore computed two different DSA distance measures: Wasserstein on the spectra, and Procrustes alignment on vector fields (or PAVF), which compares the full learned operators and not only their spectra. We will refer to these as spectral DSA and PDSA, respectively, for this analysis. We visualized the dissimilarity matrices from all methods in low dimensions, finding that the DSA variants recover the graded relationships induced by varying *ρ* between Lorenz models. Surprisingly, PDSA and spectral DSA were effectively equivalent, indicating that the spectrum was sufficient to distinguish the different parameter settings (Fig. 2i).

Moreover, the DSA distances showed the highest correlation with the largest Lyapunov exponent difference between models. It is interesting that even in a highly non-normal chaotic system, with Koopman spectrum known to be continuous (*Arbabi and Mezić, 2017*), spectral DSA performs better than expected, PDSA performs similarly well, and other methods are much less effective at capturing the Lyapunov differences between models (Fig. 2j). We additionally evaluated DSA on other non-normal dynamical systems and in the transition to chaos in the classical logistic map model (*Alligood et al., 1996*), finding that both DSA metrics were informative there as well (Figs. S3, S4).

In sum, across complex models with fixed-point, limit-cycle, and chaotic dynamics where ground truth is known, DSA and PDSA capture the dynamical relationships between systems, outperforming other methods not only in quality of the comparisons related to known dynamical structure but also in efficiency and scalability.

### DSA reveals dynamical structure of a brain compass circuit

As animals freely explore open spaces (Fig. 3a), the head direction circuit in the brain integrates angular velocity signals to update an estimated compass bearing relative to the external world (Fig. 3b, *Taube et al., 1990; Taube, 1995*). Theorists have proposed neural network models of this head direction circuit (*Touretzky and Redish, 1996; Zhang, 1996; Ben-Yishai et al., 1995; Xie et al., 2002*), which we implemented above (Fig. 2a) (*Touretzky and Redish, 1996; Xie et al., 2002*).

**Figure 3.**
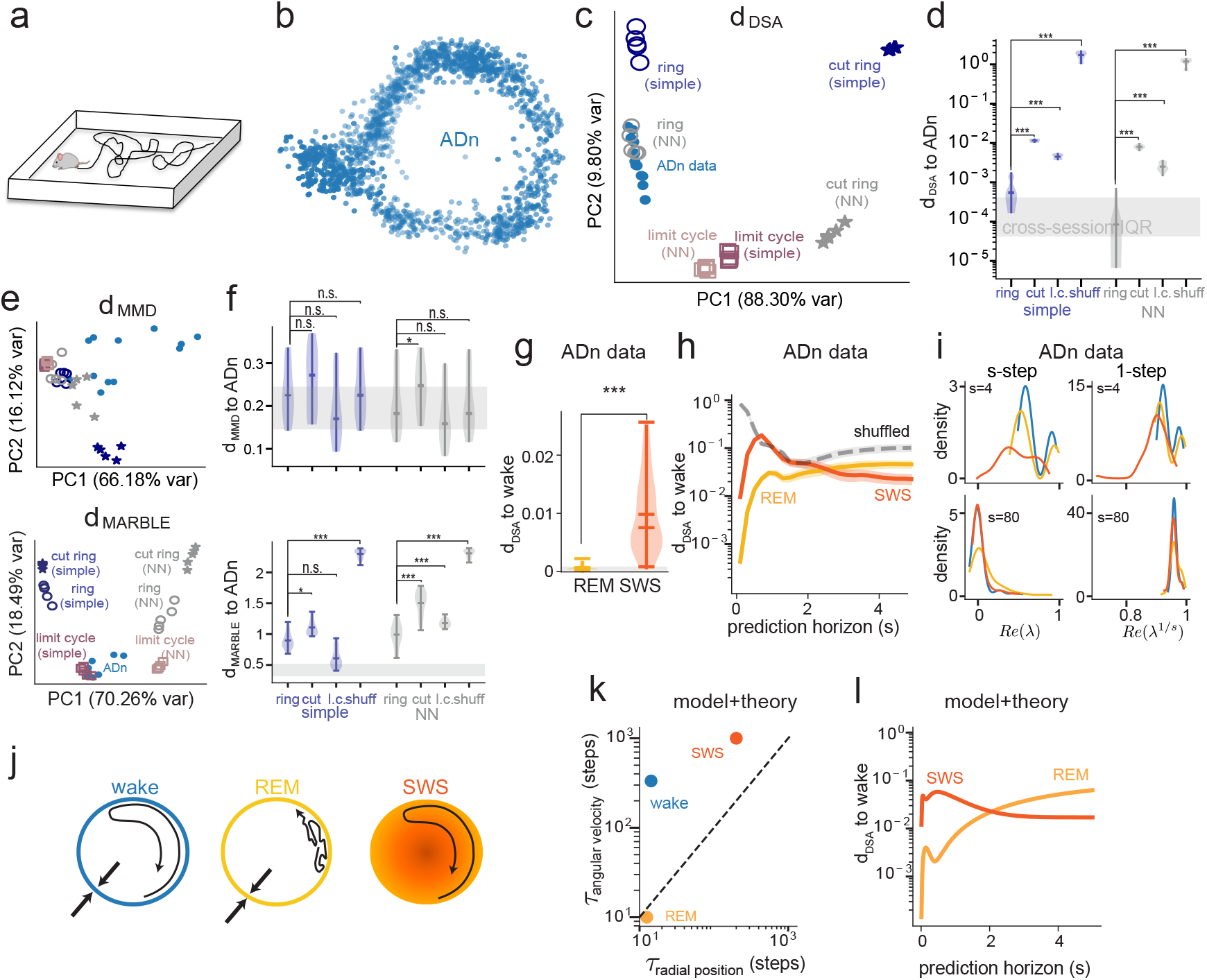
Comparing neural population data directly to dynamical models and across multiple timescales. **(a)** Mice freely exploring in an open environment. **(b)** Two dimensional embedding of the ADn population data from ~ 20 minutes of unconstrained waking exploration in an open environment like in (a). **(c)** DSA comparison of ADn recordings with all ring, cut ring, and limit cycle models (as in Fig. 2c). DSA puts ADn data closest to the neural network ring model. **(d)** DSA comparison of ADn data with all models as well as the ADn data shuffled in time (shuff). Gray shaded area indicates the interquartile range of cross-session ADn-ADn DSA distances. **(e)** PCA visualization with MMD (top) and MARBLE (bottom) between all models and all datasets. **(f)** Same analysis as (d) using the MMD and MARBLE distances. DSA distance of REM and SWS data to wake. **(h)** DSA comparison of REM and SWS to wake across multiple prediction horizons (timescales). Shuffle denotes the distance of a Wake dataset to itself shuffled in time. **(i)** Eigenvalue distributions across all ADn datasets for DMD models with a prediction horizon of four time steps (400 ms, top) and 80 time steps (8 sec, bottom). (left) true s-step eigenvalues. (right) effective 1-step eigenvalues, by raising s-step eigenvalues to the 1/s power. **(j)** Schematics of the hypothesized differences between sleep stages. Wake: coherent angular sweeps along the manifold, strong radial dynamics. REM: diffusive angular sweeps, strong radial dynamics. SWS: coherent angular sweeps, weak radial dynamics. **(k)** Autocorrelation times from theoretical analysis of the models in (j) on the radial position and angular velocity variables. These times are directly related to the Koopman eigenvalues in the theoretical derivation of DSA distance. **(l)** Theoretically derived DSA distances for the models in (j) with the autocorrelation parameters from (k), between the wake and SWS models (orange) and wake and REM models (yellow).

Predictions of these models for neural data, for instance the stability of cell-cell relationships (*Taube, 1998; Peyrache et al., 2015*), the existence of a continuum of attractor states in a ring topology and the dynamics as attractive flows onto this ring (*Chaudhuri et al., 2019*), have been rigorously validated. However, these predictions involved conceptual features or abstractions (low-dimensionality, ring topology, etc.) that had to be identified in a bespoke fashion, and then model-data comparisons were carried out indirectly and with many separate and cumbersome analyses on these bespoke features (*Taube, 1998; Peyrache et al., 2015; Turner-Evans et al., 2017; Chaudhuri et al., 2019*). Here DSA presents an interesting opportunity: to directly compare neural dynamics data from the compass circuit with data sampled from the computational models, without bespoke abstraction (Methods C).

The ADn data were collected during waking behavior (Fig. 3b), as well as during rapid eye-movement (REM) and slow-wave sleep (SWS) (Methods C.1). As with the ring models above, ADn data lacked trial structure, however DSA is able to handle temporal comparisons without timepoint alignment.

We focused first on waking dynamics. DSA preferentially clustered the neural dynamics near the neural network ring attractor models, and away from the simple rings, cut rings, and limit cycles (Fig. 3c), with significantly higher similarity to the ring attractors than any other model (Fig. 3d, one-sided Wilcoxon signed-rank test, *N* = 12 paired sessions averaged over 5 independent model comparisons, all *p* < 0.00025 with test statistic 0, indicating 0 overlap between distributions, Cliff’s *δ* = 1.00). Furthermore, the neural network ring attractor model was indistinguishable from ADn data under DSA, as the distribution of neural network ring attractor comparisons to ADn data was contained within the interquartile range of the ADn-ADn distances.

Further, we compared the ADn data to the neural network ring attractor data shuffled in time, which preserves the manifold of states while destroying the attractor structure, finding that DSA put the shuffled model farthest from ADn data, and indeed from all the other models (Fig. 3d), indicating that attractive dynamics toward the ring are necessary for alignment with the data.

By contrast, the best alternative methods (MMD by silhouette score and MARBLE by linear decoding, according to Fig. 2e) did not provide a meaningful clustering of ADn data with the models (Fig. 3e-f). MMD finds no distinction between the shuffled and unshuffled ring models (Fig. 3f, top), as expected because their point clouds are equivalent while their dynamics are not. MARBLE clusters ADn data with the simple ring limit cycles (Fig. 3e, bottom), which are two-dimensional oscillators, and away from both neural network limit cycles and neural network ring attractors. However, this result highlights an inconsistency in MARBLE’s similarity comparisons that DSA does not share: if its placement of ADn closer to the simple limit cycle than the simple ring is because ADn dynamics are more limit cycle-like, then it should also place ADn closer to the NN limit cycle than to the NN ring, which it does not (Fig. 3e-f, bottom). Moreover, the known behavior of ADn wake dynamics involves briefly coherent but overall quasi-random sweeps in angle, suggesting the MARBLE result is a mischaracterization of ADn’s dynamics.

To ensure that our conclusions were not artifacts of particular DSA hyperparameter choices, we performed systematic sweeps, finding that DSA preferentially clustered ADn data with the ring over the cut ring for both simple and NN models (96.9% of DSA hyperparameters for the simple rings and 82.9% for the NN rings, Fig. S5a), across a wide range of hyperparameters. We further identified that results were robust whether ADn data were preprocessed with Isomap or PCA (Fig. S5b). We additionally performed two noise and subsampling robustness analyses (Fig. S5c,d), adding progressively larger amounts of observational Gaussian noise and recomputing the DSA distance from each model class to the ADn data. For each model class, the orderings produced by DSA were highly robust and almost invariant to noise, even when the noise variance was equal to the data variance (Fig. S5c). In the NN model, we progressively subsampled the observed subset of neurons until only 1% of the neurons were observed and recomputed the DSA distances from each model class to the ADn data with 5% Gaussian noise added (Fig. S5d). Here too, DSA appeared almost unaffected. Thus, DSA robustly demonstrates for the first time through direct data-model comparison (without needing to compare bespoke abstractions) that neural network ring attractor models are indistinguishable in their dynamics from the head direction circuit of the mouse.

#### DSA permits interpretable comparisons of dynamics at different timescales

We next asked whether DSA could reveal interpretable differences in ADn dynamics across behavioral states. We began by computing DSA distances between each REM and SWS dataset and all waking datasets.

Waking and REM ADn states are known to exhibit fast (radial) relaxations to a 1-dimensional ring (*Chaudhuri et al., 2019*). During waking, states undergo temporally coherent, long timescale angular sweeps that are quasi-random, matching exploration dynamics as animals forage for randomly scattered food pellets. In REM sleep, the angular dynamics follow diffusive random walks in angle (*Chaudhuri et al., 2019*). In SWS, the manifold of states becomes a cone-like 2-dimensional surface with both diffusive and coherent sweep dynamics in the angular variable. The SWS radial dynamics are not confined to a 1-dimensional ring, and therefore do not exhibit fast radial relaxations; instead, the states exhibit slow and strong modulations of the amplitude of population activity, induced by SWS oscillations (*Chaudhuri et al., 2019*).

DSA reports REM-wake distances to be significantly smaller than the SWS-wake distances (Fig. 3g) (*p* < 0.00025, one-sided Wilcoxon signed-rank test with test statistic 0 and Cliff’s *δ* = 1.00; the REM-wake distribution lies entirely to the left of and does not overlap the SWS-wake distribution, *N* = 12 wake sessions, paired across 108 wake–REM and 108 wake–SWS pairwise comparisons; Methods C.3). Moreover, REM-wake distances fell within the inter-animal wake-wake distance distribution (Fig. 3g). To more finely parse this assignment and relate it to prior behavioral characterizations of the ADn data (*Chaudhuri et al., 2019*), we recall that DSA rests on the Koopman Operator (eDMD model) to predict data in the future given past states. The prediction horizon *s* is the number of timesteps into the future that the Koopman Operator (*K*_*x,s*_) evolves the state:

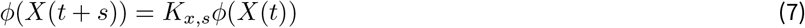

When this operator is trained to predict further into the future by increasing *s*, slow coherent modes dominate over faster modes (SI Sec. H). Thus long-(short-) horizon DSA should enable us to separately examine slow (fast) dynamics, and thus compare dynamical similarity at different timescales.

By varying the time-horizon, DSA identifies waking dynamics to be more similar to REM than to SWS for short horizon prediction, and waking to be more similar to SWS at longer horizons (Fig. 3h). Consistent with this, the Koopman spectra (eigenvalues of *K*_*x,s*_) of waking and REM are closer at short horizons, while waking and SWS spectra are closer at long horizons (Fig. 3i, left column).

We examined the interpretation that short and long horizon prediction captures the fast and slow dynamics of the system, respectively, by looking at the real parts of the principal *s*-th roots of the eigenvalues of the learned *K*_*x,s*_ on ADn data; taking the *s*-th root of an *s*-step horizon model shows the timescales that model emphasizes for a single timestep of evolution. Confirming our expectation, the distribution of long horizon eigenvalues are much more peaked near unity than the short horizon models, reflecting slow-decaying activity modes (Fig. 3i, right column). Thus, the long horizon models predominantly capture slow ADn dynamics while the short horizon models emphasize fast ADn dynamics.

The crossover (Fig. 3h) from smaller REM-wake DSA distance at fast timescales to smaller SWS-wake distances at slower timescales therefore maps well onto the previously characterized dynamics of the ADn circuit (*Chaudhuri et al., 2019*): the fastest timescale ADn dynamics consist of rapid radial relaxations to the 1D ring attractor, which are similar only between REM and waking dynamics. The longer timescale dynamics in ADn involve angular trajectories, which are better matched in waking and SWS, because they both involve some coherent angular sweeps, while REM angular dynamics are random and diffusive.

Finally, we confirmed our interpretation of the ADn crossover (Fig. 3h) by an analytical derivation of DSA distances on a simple set of differential equations modeling waking, REM, and SWS dynamics (Fig. 3j). In these simple models, the angular and radial (amplitude) dynamics were qualitatively matched to ADn waking, REM and SWS data (Fig. 3j) based on relative autocorrelations of dynamics along the angular and radial directions (SI I). Our analytical theory shows that if the model equation parameters take into account the relative ordering of radial and angular autocorrelation timescales of ADn sleep stage data (Fig. 3k), then DSA will emphasize radial relaxation similarities at short horizons (one timestep ahead) and angular dynamics at long horizons, thereby clustering model REM with model wake at short horizons and model SWS with model wake at longer horizons under DSA (Fig. 3l, Fig. S6a,b). This theoretical model exhibits a qualitatively equivalent crossover profile as seen for the neural data (Fig. 3h). We also replicated the empirical ADn result when smoothing the ADn data with successively wider Gaussian kernels, which suppress higher frequencies in the data (Fig. S6c). Thus, we have confirmed in many ways that DSA parses similarity for dynamics at fast or slow timescales based on the horizon at which the Koopman model is fit, and on the ADn data this translates to a preferential focus on radial or angular dynamics, respectively.

These results demonstrate how DSA can be used to compare not only overall dynamical similarity, but also similarity within specific dynamical timescales.

### DSA for cross-context comparisons in neural control of movements

DSA enables comparison of dynamics across contexts, which could refer to a change in the system’s inputs (e.g. during different computations performed by the same brain circuit), different instances of the dynamical system (e.g. different individuals performing the same task), or some change that occurs over time (e.g. due to learning). Prior work examined macaque motor cortex dynamics during reaching across animals, task contexts and time (*Safaie et al., 2023; Gallego et al., 2018, 2020*), but was limited to making comparisons when the trajectories could be aligned. Because the data distribution and inputs can differ in important ways across conditions (e.g. because the reach targets and durations change between tasks), it is often not possible to perform the alignment necessary for geometric methods, and the comparison method must take into account input differences. It has therefore been challenging to compare the underlying neural dynamics, making this an ideal use-case for DSA.

#### DSA shows different neural dynamics underlie similar cross-task behavioral dynamics

We applied DSA to compare motor cortical dynamics in different monkeys performing the same behavior, the same monkey performing a reaching task across many months, different monkeys performing different tasks, and different subregions of motor cortex during the same task (*Perich et al., 2025*) (Methods D). In each task, animals had to reach to targets on a screen, either starting from a center point and moving out to eight unique points on a circle (Center Out – CO) or moving between four different random points in succession (Random Target – RT, Fig. 4a). The kinematic trajectories and hand speeds exhibited high diversity across contexts (Fig. 4b, Fig. S9a,b).

**Figure 4.**
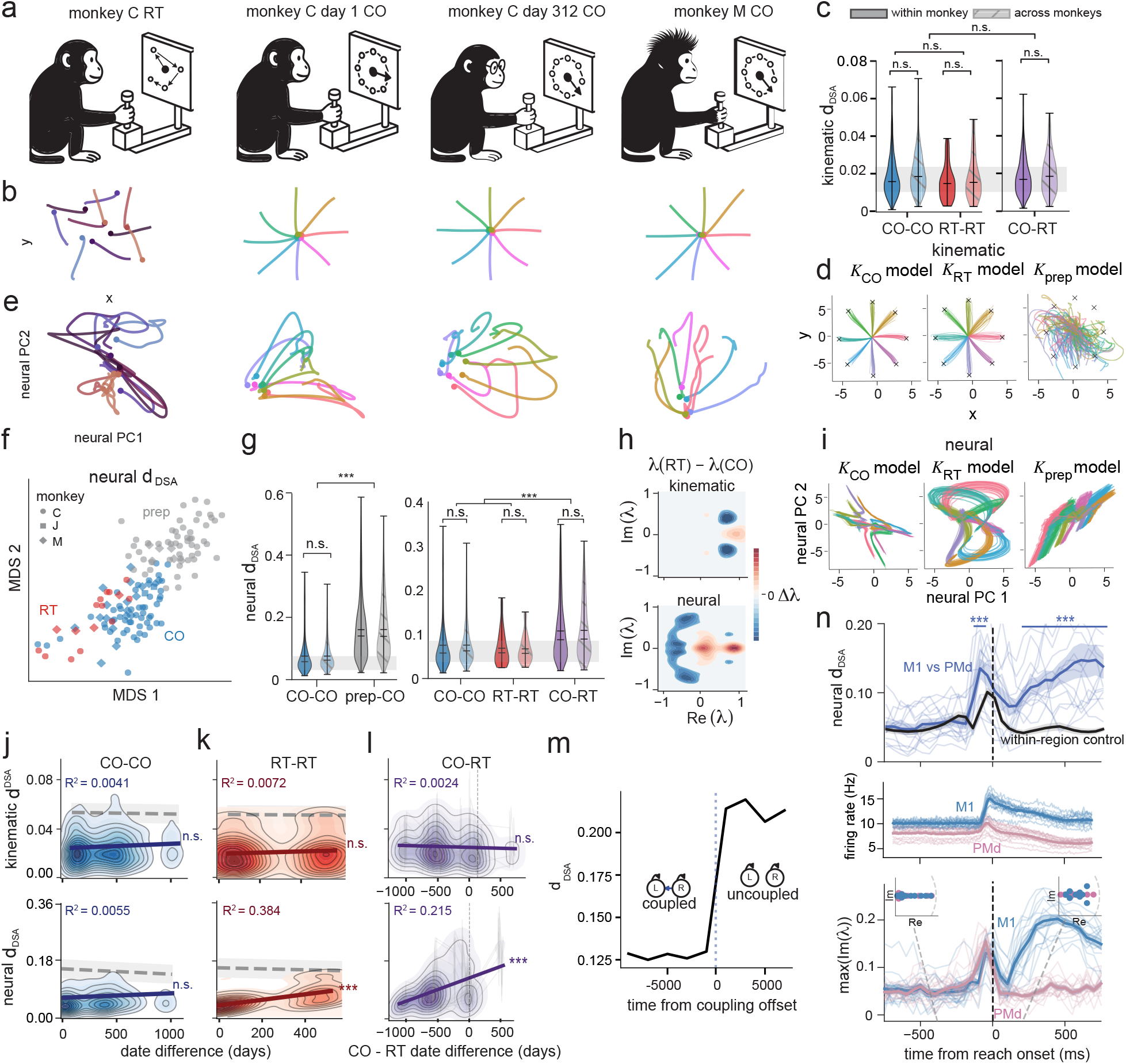
DSA identifies similarities in neural dynamics across monkeys, tasks, time, and brain regions. **(a)** Multiple macaques perform a center out (CO) and random target (RT) reaching task across multiple months. **(b)** Trial-averaged trajectories of the reaching kinematic data. Colors indicate trial conditions. RT trials are clustered by start and end point, 8 random clusters are shown. **(c)** DSA on the hand reaching (kinematic) trajectories. Gray shaded area indicates the inter-quartile range of within-animal within-task DSA distance. **(d)** Autonomous SubspaceDMDc kinematic data model rollouts for 8 kinematic conditions defined by the CO task. x and y indicate spatial position, X’s indicate the target hand position for each color. (left) CO model (middle) RT model (right) preparatory control epoch model. **(e)** Trial averaged trajectories of the neural data, top 2 Principal Components, colored and aligned with (b). **(f)** MDS plot of the neural DSA matrix between all valid neural datasets for a SubspaceDMDc model of rank 26 and 14 delays. **(g)** Cross-epoch DSA neural distances, in the same fashion as (c). ∗ ∗ ∗ denotes p < 0.001. **(h)** Difference in KDE-estimated distributions of DMD eigenvalues fit to kinematic data (top) and neural data (bottom). Differences are taken between RT and CO densities, thresholded by significance and z-scored with respect to a null label shuffle. **(i)** Autonomous SubspaceDMDc neural data model rollouts for 8 kinematic conditions as in (d), on the top 2 PCs of the neural data. **(j)** DSA distance between CO sessions within each monkey, as a function of days in between session (top, kinematic distance, bottom neural distance). Solid line indicates regression line of best fit with R^2^ coefficient labeled in top left, gray line indicates cross epoch comparison baseline. *n*.*s*. denotes insignificant under a session-pair date one-sided permutation test testing slope > 0, ∗ ∗ ∗ denotes p < 0.001. **(k)** Same analysis as (j), on the RT datasets. **(l)** Same analysis as (j,k), on the cross-task distances. **(m)** Time-windowed DSA distance between a Rössler and Lorenz attractor system before and after the systems are decoupled from one another. DSA computed on a sliding window over time. **(n)** (top) Cross-brain region neural DSA distance (PMd and M1) before and during reaching, centered at movement onset. Blue line indicates the average cross region distance, black line denotes the cross-session within-region score at the same timepoint. Shaded regions indicate individual sessions. Top bars denote significance at p < 0.001 according to a one-sided Mann-Whitney test. P-values for each window and effect sizes are given in Table S8. (middle) Average firing rate of all neurons over time in M1 and PMd, centered at movement onset. Firing rate computed on the same windows as DSA in (n). (bottom) Maximal imaginary component of neural DMD eigenvalues within each window, describing the maximal oscillation frequency within the system. Shaded lines denote individual sessions. Insets display sample eigenvalue distributions for particular time windows (denoted with gray dashed lines).

To account for the task differences, we apply a DSA variant (which we call conditional DSA) that fits Koopman Operators with an additional input or task condition variable (the eDMD formulation with this input is known as subspaceDMDc (*Takeishi et al., 2017; Huang et al., 2025*)). Conditional DSA interpolates between DSA and InputDSA (*Huang et al., 2025*, which introduced a control-based similarity metric), in the sense that like InputDSA it uses condition-aware inputs but like DSA retains a focus on the intrinsic dynamics by using the Wasserstein metric on Koopman eigenvalues. For both kinematic and neural data, we used the same input structure: a constant encoding of the target position in Euclidean space for each trial.

Unsurprisingly, given the observed behavioral consistency, DSA found reach kinematics to be highly similar across trials within each monkey and across monkeys on each task (Fig. 4c, striped violins; Table S1), such that cross-animal kinematic differences on a taskwere not significantly greater than intra-animal variability (one-sided label permutation test, *p* = 0.0188, 0.4025, 0.1361, all insignificant under Bonferroni correction for within-vs-cross animal comparisons on CO, RT and cross-task distributions respectively, Cliff’s *δ* = 0.14, 0.11, 0.09). These results were robust across hyperparameters (Tables S3 and S4) and agree with previous reports of consistent behavioral dynamics across trained animals (*Safaie et al., 2023; Gallego et al., 2020*). A one-sided test for equivalence with a margin *σ*_*w*_ at the standard deviation of the within-animal distribution yielded a similar result, finding that cross-animal distances were equivalent to within-animal distances for each comparison (*p* < 10^−5^, *p* = 1.80 × 10^−4^, *p* < 10^−5^, standardized effect sizes Δ_*med*_*/σ*_*w*_ = 0.26, 0.07, 0.17 for CO, RT and cross-task distances respectively, Methods Sec. D.4.2).

More surprisingly, when we compared kinematic dynamics between the CO and RT tasks (cross-task) within the same animal, the cross-task DSA distances were not statistically different from within-task distances (Fig. 4c, Table S1, label-shuffle permutation test, *p* = 0.1589 insignificant under Bonferroni corrected threshold *α* = 0.05*/*8, Cliff’s *δ* = 0.05). The lack of a significant difference in DSA distance between the CO and RT tasks leaves unclear whether DSA lacked sensitivity to important kinematic differences, or whether the dynamics on RT and CO tasks are genuinely equivalent. To answer this question, we used the learned Koopman dynamics operators for the kinematics as generative models. We reasoned that if the RT and CO kinematics were similar then using the RT operators with the CO task targets should reproduce the CO reach trajectories. Thus, we initialized both CO and RT Koopman models near the origin, providing CO targets as the contextual input, and let the operators autonomously evolve for the length of a trial (Fig. 4d). Both CO and RT models qualitatively replicated the CO reach trajectories, while a Koopman model trained on the preparatory epoch dynamics as a control could not. Thus, we conclude that DSA correctly identifies that kinematic dynamics are preserved across both animals and tasks.

Next, we examined the motor cortical neural dynamics that underlie arm kinematics across tasks, task phases, and animals. The average neural population responses for eight task conditions projected onto the top two principal components were visually dissimilar across tasks (Fig. 4e). We computed the neural DSA distance matrix within the RT and CO tasks, as well as between these reach tasks and the preparatory period (Fig. 4f). As with the kinematics comparisons, cross-animal neural distances for each task (intermingled symbol shapes in Fig. 4f) were not statistically different from within-animal distances (one-sided permutation test for CO, RT and cross-task within-versus cross-animal comparison, *p* = 0.0179, 0.1751, 0.3401, all insignificant under Bonferroni correction for 8 comparisons, Cliff’s *δ* = 0.14, 0.07, 0.04, Table S1, Fig. S10a-c), falling within the inter-quartile range of the within-animal within-task distances (Fig. 4g gray shaded region), consistent with prior reports of preserved cross-subject within-task neural trajectory geometry (*Safaie et al., 2023; Gallego et al., 2020*).

At the same time, the neural DSA distances were large between tasks (in the reach periods; separate clusters in Fig. 4f, with distances quantified in Fig. 4g, *p* = 9.00 × 10^−5^, one-sided permutation test, Cliff’s *δ* = 0.36, Fig. S10d, Tables S1, S3) and between the pre-reach preparatory and reach periods (Fig. 4g (left), *p* < 10^−5^, one-sided permutation test, Cliff’s *δ* = 0.660, Table S1).

Further, the cross-task neural distances far exceeded the cross-task kinematic distances, even when normalized by a baseline (within-task distance) and ceiling (cross-period, reach vs. preparatory) (Fig. S10d-e). Correspondingly, the underlying neural Koopman eigenvalue distributions for CO and RT tasks were also more distinct from each other relative to the kinematic distributions (Fig. S9d, Fig. 4h, bottom versus top, respectively: the point-wise sum of the neural CO-RT eigenvalue difference distribution, for significant values relative to a null label shuffle control, made up 48% of the total neural eigenvalue distributions; for the kinematic distribution, this difference was only 25%).

We again assessed the validity of the learned Koopman Operators for neural DSA by using them as generative models, giving each model the CO target states and allowing them to autonomously generate neural trajectories. The resulting modeled dynamics are clearly distinct (Fig. 4i) with strong rotational modes in the RT model, as expected from the Koopman eigenspectral differences (Fig. 4h, bottom). Thus, DSA dissociates motor cortical activity in different contexts by interpretable dynamical features, and reveals distinct neural dynamics in the RT and CO tasks even when the behavioral kinematics are similar, both corroborating and sharpening the conclusion of (*Safaie et al., 2023*).

#### DSA reveals phenomenon of differential neural representational drift for different tasks

Next, we used DSA to assess the stability of neural dynamics as two monkeys performed the CO and RT tasks over years (Fig. 4a). The question of neural stability has been addressed in imaging experiments where the same neurons can be identified across time (*Ziv et al., 2013*) and via trajectory geometric comparisons (*Gallego et al., 2020*), but no methods prior to DSA existed to measure drift in population dynamics when neurons are not maintained longitudinally.

We first applied DSA to behavior, probing for kinematic drift over years of performing the tasks. In line with prior work, we found no behavioral drift on either task (Fig. 4j,k, top, *p* = 0.2629, 0.4543, all tests in this section were one-sided date permutation tests evaluating Δ*DSA/*day > 0 with Bonferroni correction for simultaneous comparisons, *R*^2^ = 0.0041, 0.0072); cross task differences likewise did not change over time (Fig. 4l, top, *p* = 0.6162, *R*^2^ = 0.0024).

We then examined neural dynamics and found representational stability over years within the CO task (Fig. 4j, bottom, *p* = 0.2994, *R*^2^ = 0.006). Basic statistics such as number of neurons recorded, average neural firing rate, and PCA dimensionality (via participation ratio) remained consistent across the CO and RT tasks (Fig. S11a). Trajectory geometry comparison methods (with CCA as in *Gallego et al., 2020*) detected no notable trends in CO or RT representations across recording dates (Fig. S11b).

By contrast, DSA found that the RT neural representations drifted significantly despite behavioral stability (Fig. 4k, bottom, *p* = 9.995 × 10^−4^, *R*^2^ = 0.384). Likewise, the CO-RT neural DSA distance increased significantly over time (Fig. 4l, bottom row; Table S5, *p* = 9.995 × 10^−4^, *R*^2^ = 0.215). These drift findings were robust across DSA hyperparameter sets, with the slopes of neural DSA distance over time for RT and CO-RT roughly an order of magnitude larger than corresponding slopes for any of the kinematic distances (Tables S5, S6, Fig. S11c), and roughly 5 times larger than the within-CO neural slope. CO sessions were far more numerous than RT ones (70 vs 16, respectively). To control for this difference, we subsampled CO sessions to match the number of RT sessions and recomputed the slope, finding here too that the CO slope was not significant, with a 95% confidence interval of [−3.313 × 10^−5^, 6.131 × 10^−5^] *d*_*DSA*_/day for the neural data and [−7.084 × 10^−6^, 6.376 × 10^−6^] *d*_*DSA*_/day for the kinematic data. Parsing the CO-RT DSA distance by session dates suggests that the early RT neural dynamics were highly similar to the CO sessions, but later RT sessions diverged (Fig. S11d).

We separately evaluated the predictive capacity of the learned Koopman models across sessions–if the dynamics drift over time, a model fit to early sessions should more poorly predict a later session, with error increasing as a function of date difference. DSA learns an optimal alignment between eDMD models, making this predictive comparison well-posed even when cells are not recorded longitudinally. We found that the DSA-aligned Koopman models successfully generalized across CO sessions over the entire years-long recording window, but did not for RT (Fig. S11e, *p* = 0.6830, *p* < 10^−4^ for CO and RT respectively, one-sided slope permutation test evaluating slope > 0).

In sum, DSA identified a novel phenomenon of a differential drift on one task relative to another within the same circuit, motivating potential future study of the mechanistic causes, e.g. whether RT dynamics are intrinsically more prone to reorganization than CO dynamics; or whether the CO dynamics were more stable due to more frequent practice on the CO than the RT task (70 versus 16 sessions, respectively, Fig. S11a), although the sessions were interleaved over time and behavior was equally stable. More generally, while (*Gallego et al., 2020*) demonstrated uniform long-term stability in neural population geometry, our results open the door to questions of *when, why, and how* neural dynamics drift even if geometry remains stable.

#### M1 becomes distinct from PMd at movement onset

We additionally leveraged the motor control data to consider a different use case for DSA: whether it can reveal dynamical modulations in the interactions between two systems. Before movement onset, premotor area PMd is believed to drive M1 into an active, movement-generating state (*Kaufman et al., 2014; Elsayed et al., 2016*). When two different systems are coupled, they share Koopman eigenvalues (SI J), thereby lowering their DSA distance; when the systems evolve independently, all else equal, their DSA distance will be higher. Because DSA is symmetric, it can quantify the degree of dynamical coupling but not its direction. We first tested this when the coupling is known, by simulating a Lorenz and a Rössler attractor with the second Rössler dimension driving the second Lorenz dimension; then subsequently removing the coupling. DSA detected the decoupling as a sudden increase in *d*_*DSA*_ distance between the two systems (evaluated by comparing the systems in sliding local time windows, Fig. 4m). This suggests that DSA is capable of detecting changes in coupling between dynamical systems.

We applied this to estimate the coupling between PMd and M1 in 300 ms sliding windows (50 ms increments), straddling the time of movement onset in the CO task. Interestingly, the dynamical dissimilarity between the two significantly increased just before movement onset, dipped transiently, and remained elevated throughout the reach (Fig. 4n, top). This temporal variation in PMd-M1 DSA distance is not explained by changes like simple variations in neural firing rates near movement onset (Fig. 4n, middle), which have distinct temporal profiles with the rate changes trailing the first DSA peak. Analysis of the Koopman eigenvalues (Fig.4n, bottom) reveals that at least one oscillatory mode emerges in M1 shortly before movement onset and persists throughout movement, consistent with prior reports of rotational M1 dynamics (*Churchland et al., 2012*). By contrast, a transient oscillatory mode emerges in PMd immediately before movement onset, consistent with prior analysis (*Elsayed et al., 2016*). The DSA distance and its Koopman eigenvalues therefore show that PMd rapidly becomes dynamically distinct from M1 via a transient rotation before movement onset, after which M1 generates rotational dynamics that drive movement and further dynamical separation.

Overall, the results in this section illustrate how DSA provides insight about relative dynamics in diverse and rich real-world systems from empirical data.

### DSA for the generation of novel neural network task solutions and models

So far we have primarily considered DSA as a direct comparator of sampled data from multiple dynamical systems. Here we consider a distinct use of DSA: to encourage the discovery of dynamically diverse solutions to a problem (Fig. 5a). Motivated by recent work (*Qian and Pehlevan, 2026*), we apply DSA as a regularizer during optimization (Methods E). Regularizing away from existing solutions rather than encouraging similarity to a desired solution is a powerful tool for discovery, because it does not require a priori hypotheses about alternative solutions.

**Figure 5.**
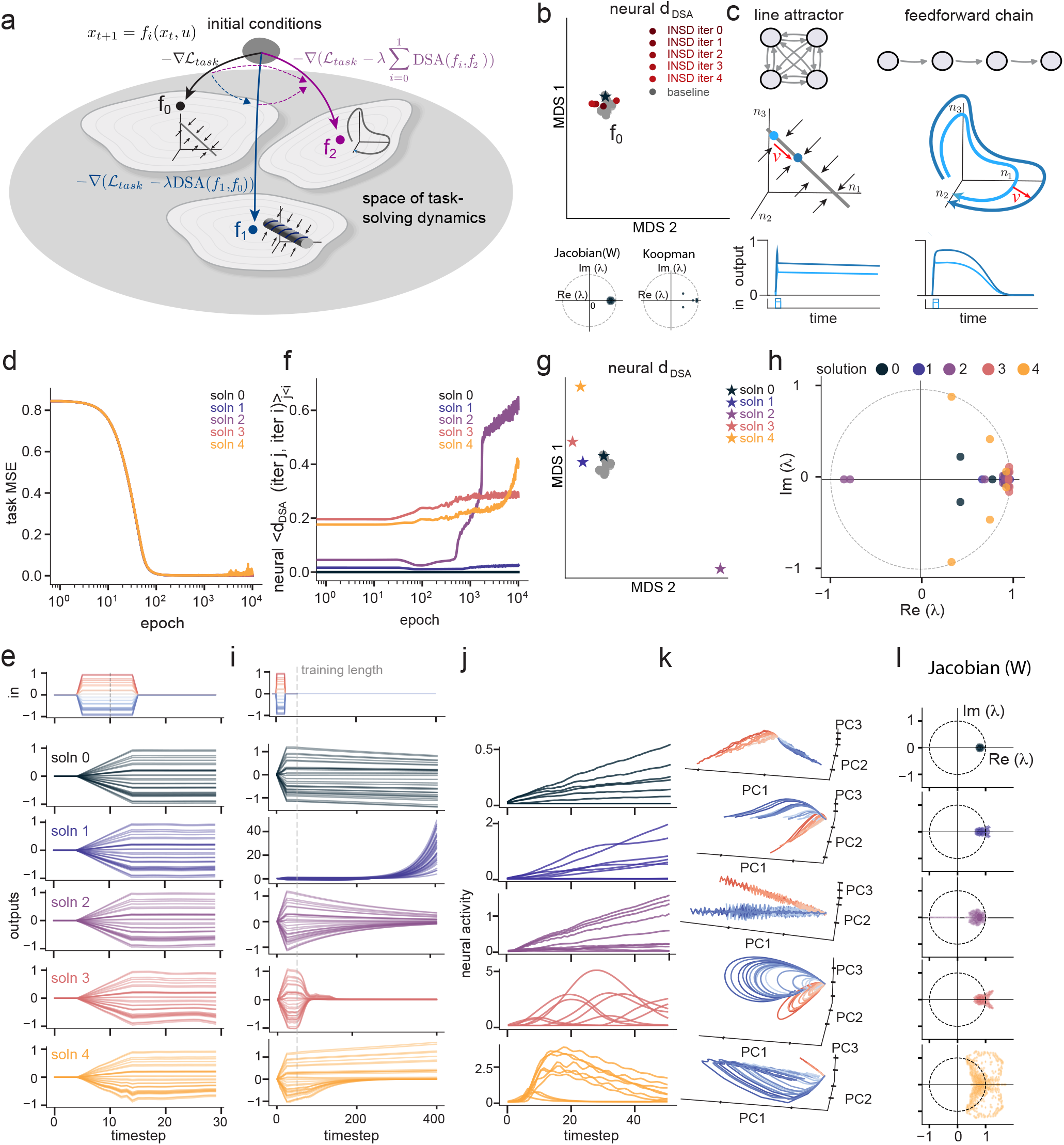
DSA regularization pushes RNNs to discover diverse and novel dynamical solutions to an integration task. **(a)** Schematic of the iterative DSA regularization procedure and how it drives dynamics to different qualitative solutions by repelling from previous ones. RNN 0 is trained on the task alone, then each RNN is trained on the task as well as the auxiliary goal of maximizing the sum of DSA distances between all of the previous models’ hidden state dynamics. Insets denote sample dynamical structures for each solution type. **(b)** Distribution of RNNs trained without DSA regularization. DSA distance matrix visualized in low dimensions with MDS, Iterative Neural Similarity Deflation (INSD, *Qian and Pehlevan, 2026*) iterations colored in red and baseline trained RNNs in gray. Bottom insets: Jacobian and Koopman eigenspectra for the starred solution (f_0_). **(c)** Schematic of line attractor and feedforward chain dynamics. (top) Connectivity graph, (middle) dynamic landscape of the models, with 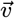 denoting how input changes the activity, (bottom) output behavior after input is turned off. **(d)** Learning curves on the task for each iteration (overlapping). **(e)** (top) Sample time series of inputs with varying magnitudes (color) provided to the models. (bottom) sample output time series for each solution. **(f)** Average DSA distance of each iteration to all previous iterations across training epochs. **(g)** DSA distribution as in (b) with iteratively regularized solutions. Baselines are shaded gray as in (b). **(h)** Eigenvalue distribution in the complex plane of the Koopman models for each iteration. **(i)** Relaxation dynamics of the output of each RNN after input has been provided for 25 steps, and then turned off. **(j)** Visualization of the dynamics of 10 random neurons from each model for a single trial of input. **(k)** Visualization of the dynamics in the PCA space when input (colored by magnitude) is provided. **(l)** Jacobian eigenvalues of each model aggregated across multiple trials.

High-performance optimizers such as stochastic gradient descent can be effectively applied to dynamical systems models and recurrent neural networks (RNNs), but strong inductive biases from architecture and training routines drive convergence to restricted classes of solutions. This reduces diversity, potentially excluding solutions that may be a closer match to the natural system being modeled, that generalize better, that are more robust to adversarial attack (*Dapello et al., 2020*), or that are desired for other reasons. This limitation is easily demonstrated on the task of accurately integrating a time-varying scalar input, for which RNNs are biased towards line attractor solutions (*Maheswaranathan et al., 2019; Mante et al., 2013*). When we trained 430 RNNs with various seeds and hyperparameters (including learning rate, hidden state dimension, recurrent weight initialization standard deviation, batch size, and weight decay) the solutions reached are similar to each other, with the recurrent dynamics of different models clustered in the DSA space (Fig. 5b, top; grey).

We analyzed these solutions by studying the Jacobian matrix of their dynamics, which for ReLU RNNs is simply the submatrix of the recurrent weights restricted to the active neurons. The Jacobian spectrum of a baseline RNN (Fig. 5b, bottom-left) matches expectations for a low-dimensional continuous attractor network (*Seung, 1996*), with all eigenvalues essentially real-valued with one eigenvalue maintaining norm close to 1. The rest have norm less than one, corresponding to leaky or decaying modes (Fig. 5b, bottom-left for the solution *f*_0_, marked by a star, Fig. S12a,b).

In contrast to the monotony of these baseline RNN solutions, the integration task is theoretically known to be solvable in different ways with recurrent neural networks (Fig. 5c). One theoretical construction (*Seung, 1996*) is a line attractor: symmetric excitatory recurrent connectivity generates positive feedback, canceling activity decay and creating a persistent mode with a near-unity eigenvalue, along which inputs drive state-changes that reflect the integrated value of the inputs. Our baseline RNNs resemble this construction (Fig. 5b). A dynamically distinct solution to the task involves a feedforward chain (*Goldman, 2009*), which maintains memory by propagating the state in a feedforward sequence through the network’s nodes (Fig. 5c, top right). This solution manifests as rotational dynamics in the activity space (Fig. 5c, middle right). RNNs appear strongly biased against this solution under standard training, and an existing strategy to drive the learning trajectory away from other solutions (*Qian and Pehlevan, 2026*) did not succeed at learning feedforward chains under a comparable training budget and across a sweep of hyperparameters (Fig. 5b, Fig. S13).

We thus asked whether DSA could help RNNs discover the feedforward chain solution, among others. We implemented solution repulsion by defining ∂DSA, a differentiable version of DSA that utilizes the differentiable Sinkhorn distance from optimal transport (*Cuturi, 2013*) in place of the Wasserstein distance (Methods E.2). Across two distinct tasks (integration; 3-bit flip-flop, Fig. S14 (*Sussillo and Barak, 2013*)), ∂DSA regularization reliably increases dynamical diversity over unregularized baselines (Fig. 5). We demonstrate this here on the integration task. We iteratively trained five RNNs on an integration task, beginning with the starred solution (“solution 0”, Fig. 5b). At each gradient step, the model being trained seeks to minimize task loss ℒ_*task*_ while jointly maximizing ∂DSA distance between its hidden state dynamics and those of all previously trained RNN solutions (Fig. 5a):

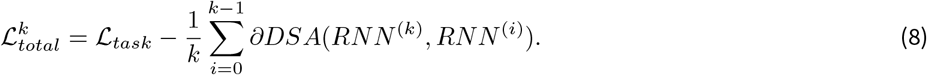

All model iterations learned to effectively solve the integration task within 1000 epochs (Fig. 5d,e), although the unregularized learning process (solution 0) had a slightly lower test error (Fig. S15a). Consistent with this, the DSA distances between the outputs of the iteratively trained solutions were all essentially at zero (Fig. S15b). However, average DSA distances of each ∂DSA-regularized RNN to all prior solutions (evaluated over the trial window) grew over training (Fig. 5f). After training, each RNN solution was identified to be distinct by DSA, with the distances between solution-repelling networks outside the baseline distribution of unregularized network solutions (Fig. 5g and Fig. S15c).

To interpret the dynamics of the learned solutions, we considered the learned Koopman Operator eigenvalues for the hidden state dynamics (Fig. 5h), their dynamics over a window beyond the trained integration window (Fig. 5i), their hidden unit activation profiles (Fig. 5j), their population dynamics (Fig. 5k), and the Jacobian eigenspectra of the trained networks (Fig. 5l).

Together these illustrate the nature of the distinct dynamical solutions found with DSA regularization, showing that the networks learn the two qualitatively distinct theoretical proposals and additionally generate entirely new, previously un-hypothesized solutions to the integration task. Specifically, solutions 0 and 3 map to the two theoretical proposals. This confirms that ∂DSA regularization can be used to drive RNNs to form feedforward chain solutions to the integration task, which was not learnable via standard training or via the method introduced by *Qian and Pehlevan, 2026* (Fig. S13). Notably, solution 3 utilizes a different mechanism to produce the feedforward chain than the theoretical linear feedforward chain (*Goldman, 2009*). Whereas the linear feedforward chain utilizes non-normality to propagate memory across nodes, solution 3 appears to use transient growth via unstable local dynamics as well. Solution 1 constructs a higher-dimensional (planar or hyperplanar) attractor, with at least two near-unity Jacobian eigenvalues within each local region (Fig. S12), and it computes the integrated input along more than one of these dimensions. It is also a slightly unstable or mistuned system, as running the model for long periods of time leads to divergence of the state. Solution 2 is a qualitatively different solution in the sense that it integrates along one dimension with a near-unity real Jacobian eigenvalue as for solution 0, but also introduces oscillatory dynamics along orthogonal directions, a combined oscillating-integrating system. Finally, solution 4 finds a different hybrid solution, combining persistent state dynamics like solution 0 for positive direction excursions and transient feedforward chain-like behavior like solution 3 for the negative direction.

We additionally applied ∂DSA regularization to identify novel solutions on a 3-bit flip-flop task (*Sussillo and Barak, 2013; Qian andPehlevan, 2026*) finding that not only did it generate a distinct oscillating solution (*QianandPehlevan, 2026*), but additionally identified a more complex double-lobed oscillator solution as well as other non-standard, non-oscillating solutions (Fig. S14), and did so in one-tenth the training iterations of *Qian and Pehlevan*.

Together, these results show that DSA can induce RNNs to learn a variety of qualitatively different dynamical solutions, and that regularization with ∂DSA induces changes in *mechanism* through the RNN weights (Fig. S15, S12). While *Pagan et al. 2025* required hand-designing RNN solutions to show that rats achieving identical decision-making performance did so in diverse ways not spanned by standard RNN training, DSA provides an unsupervised way to explore the space of solutions and to compare these with data. Beyond offering a scalable route to relating model diversity to individual variability observed across animals, our results further suggest that ∂DSA could be applied to the problems of hypothesis generation and quantification of the size and shape of solution spaces in RNNs.

## Discussion

DSA fills a gap in the space of comparisons for complex dynamical systems by providing a direct data-based way to quantify the similarity of these systems based on their flow-fields. It is theoretically motivated, scalable, interpretable, and robust to partial observability and noise. DSA efficiently approximates the intractable but idealized goal of computing distances from conjugacy. It does so by leveraging the theory of Koopman Operators to replace a search for conjugacy maps through infinite-dimensional Hilbert spaces with linear system comparisons. DSA uses the spectra or other summaries of the linear systems to compute a similarity measure. DSA is a generic and modular tool, with variations that involve different data embeddings, autonomous or input-driven dynamical models, and various similarity measures including differentiable metrics. The components are simple and this simplicity enables scalability: Across analyses in this paper, we performed on the order of 2 million DSA comparisons.

We applied varieties of DSA to diverse data from neuroscience for illustration, because it can be contextually rich and varied, high-dimensional, and noisy. However, the method is based on general dynamical systems theory developed outside of neuroscience and machine learning, and we evaluated the method on problems in machine learning as well as classical models of chaos in dynamical systems (the Lorenz attractor, the Rössler attractor, and the Logistic map) to demonstrate its broad applicability. Koopman Operator theory and associated empirical methods like the Dynamic Mode Decomposition already have strong footings in fluid dynamics, control theory, and climate science. Yet even in these fields, principled methods for comparing dynamical systems from sampled data have been lacking. DSA could be used to validate digital twins against real-world control systems, benchmark reduced-order models against numerical simulations, identify regime shifts in climate, power grid, or financial market data, or quantify how dynamics change under perturbation in cardiac, ecological, or gene-regulatory systems.

Within neuroscience, we illustrated how DSA enables novel scientific findings that were not possible without it, because many prior comparison methods require time-alignment or other matching techniques (if based on trajectories), or they ignore dynamics (if based on point clouds). By contrast, DSA can compare dynamics across systems under far fewer constraints on how data are acquired and aligned, which allowed us to extend a number of recently published works: DSA showed for the first time, via direct comparison of data (*Peyrache et al., 2015*) with models (*Zhang, 1996; Xie et al., 2002*), that canonical continuous ring attractor models are dynamically equivalent to the head direction circuit, contrasting with more indirect comparison approaches (*Peyrache et al., 2015; Chaudhuri et al., 2019*). Applying DSA at different timescales revealed interpretable changes in the head direction circuit’s dynamics across different brain states. On the neural control of movement (*Safaie et al., 2023*), DSA enabled quantification of the similarity of cross-animal neural dynamics even when the behavioral trajectories did not precisely match, and DSA further showed the existence of a novel differential, context-dependent representational drift: underlying stable behavior across two tasks, the same circuit reorganizes its neural dynamics for one task over years while its dynamics for the other task remain stable. Third, DSA provided an unsupervised approach to explore the space of solutions to computational tasks beyond the inductive biases of RNNs, which could be used to understand the diversity inherent in animals, as in (*Pagan et al., 2025*).

As we have seen, besides data-data comparisons, DSA permits theoretically motivated, easy, and fast comparisons of data with models, provided that one possesses a set of candidate models. In addition, DSA enables novel model discovery and diverse hypothesis generation by acting as a regularizer that repels newly learned models from previous ones. It could therefore be leveraged to understand the structure and size of the solution space to a problem, across domains and fields. It can also be used to create new models of systems with unsatisfactory existing models: if data-model comparison yields poor agreement for a set of existing models, DSA as a regularizer could generate new models that are a better fit to the data. In sum, we believe that DSA will be a valuable tool in any scientific toolkit for studying time-varying systems.

### Limitations and future directions

In its current implementations, DSA is inherently a comparative metric without absolute scale, because scaling can change across hyperparameters (scaling can depend nonlinearly on the number of eigenvalues). Thus, the distances are meaningful relative to a within-condition baseline (e.g. within-animal or within-task variability). Second, our default summary statistic– an optimal transport distance on the discrete eigenspectrum – captures systems dominated by isolated eigenvalues (fixed points, limit cycles, quasiperiodic and attractor dynamics) but it does not fully resolve the continuous spectral component or pseudospectra of strongly mixing, non-normal, or chaotic regimes. For the case of continuous eigenspectra, we showed that a DSA summary statistic can instead be based on methods like Procrustes Alignment of Vector Fields (PAVF), although the spectral metric in practice does almost as well.

Third, the eigenvalue summary discards the geometry of the Koopman modes, so two systems with identical spectra but different eigenvector structure (and therefore different transient, non-normal dynamics) will be scored as similar; for this reason the default distance is a pseudometric rather than a metric, and distinguishing such cases requires mode- or pseudospectrum-based variants (Grassmannian, PAVF distances, the former of which requires aligned bases). Fourth, results depend on the embedding dimension, operator rank, delay length, and sampling rate; we selected these by AIC and ResDMD (*Colbrook et al., 2023*) and verified the choices with hyperparameter sweeps, but poor choices could reduce the accuracy of the comparisons. Finally, though real recordings are stochastic and partially observed, the nonlinear and delay embeddings used by DSA allow it to generate robust estimates of dynamical similarity so long as the chosen embedding approximately identifies a closed Koopman subspace. We regard these as scope conditions, not fundamental flaws: DSA should be applied with thorough evaluations on model fit quality, and the modular nature of DSA should be exploited to select appropriate embeddings and summary metrics as relevant to the problem. Future work will include the development of novel metrics and variants of DSA that expand to fill domain-specific needs, as well as the application to fields well beyond neuroscience and machine learning.

## Author Contributions

M.O. conceived the method with contributions from all authors. W.T.R. contributed to the conceptualization of the Wasserstein metric and the regularization analysis. M.O. performed all empirical and theoretical analyses and wrote the software, with contributions from L.K. and A.J.E. A.J.E., L.K., and I.R.F. advised on the analyses. I.R.F. supervised the project. M.O. and I.R.F. wrote the original manuscript. All authors edited the manuscript.

## Code availability

Code to implement DSA is available at https://github.com/mitchellostrow/DSA.

## Data availability

All data analysed in this study are publicly available. The macaque neural recordings were collected by Perich, Miller and colleagues and are available from the DANDI Archive, dandiset 000688 (https://dandiarchive.org/dandiset/000688) (*Perich et al., 2025*). The head-direction (anterodorsal thalamus) recordings are those of Peyrache et al.(*Peyrache et al., 2015*) and are available from CRCNS, dataset th-1 (http://crcns.org/data-sets/thalamus/th-1).

## Funding Acknowledgements

M.O. is funded by the NSF GRFP under award number 2024368654. This project was also supported by NIH 1U19NS123716 and NIH U19NS132720 to I.R.F.

## A Methods

Throughout the paper, all main-text analyses are reported using a representative set of DSA hyperparameters (rank and delay-embedding length); cross-hyperparameter results and the tuning curves used to select them are documented in SI (Fig. S7 for the head direction and ring data; Fig. S8 for the macaque reach data). Hyperparameters were selected based on minima of the AIC and ResDMD tuning curves (SI F.3). Robustness of all main conclusions to this choice was verified by aggregating across the full hyperparameter sweep, holding out poorly fitting DMD models on either AIC or ResDMD (see SI Fig. S5, SI Fig. S10, and Tables S1–S5). Degrees of freedom are not defined for the rank-based and permutation tests used throughout; we report the test statistic, exact or floor p-value, and effect size instead (Methods D.4.3).

All computation was done on MIT’s Engaging computing cluster, using a node with 32 CPUs and 128 GB of RAM for efficiency, although significantly less compute is needed in practice. All distance measures can be computed in parallel (each pairwise distance is independent). RNNs were trained on a single A100 GPU.

## B Method comparison on synthetic dynamics

A dataset consists of trajectories 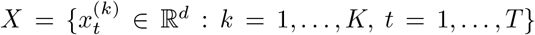, and each pairwise comparison method produces a scalar distance *d*(*X, Y*). Throughout this section, the random-feature map

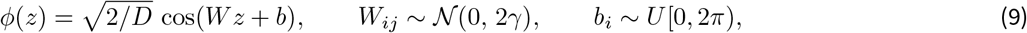

denotes Random Fourier features (RFF) approximating the RBF kernel *k*(*z, z*^*′*^) = exp(−*γ*∥*z* − *z*^*′*^∥^2^), where *D* is the number of features and *γ* the inverse-bandwidth. In order to approximate the Cauchy Kernel *k*(*z, z*^*′*^) = 1*/*(1 + *γ* · ||*z* − *z*^*′*^||^2^), RFFs were used with the weights *W*_*ij*_ ~ Laplace(0,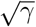) (*Rahimi and Recht, 2007*).

### B.1 Ring model details

#### B.1.1 Neural ring model

We simulated a biologically-constrained ring attractor network following the dual-population architecture described in *Xie et al., 2002* with implementation details derived from *Chaudhuri et al., 2019*. The network consisted of two coupled populations of *N* = 512 neurons each, representing left- and right-preferring subpopulations. Neurons within and across populations were connected via a shift-asymmetric weight profile:

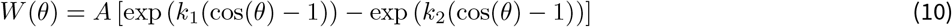

where *θ* is the angular difference in preferred head direction between neurons. As in *Chaudhuri et al., 2019* we used *A* = 1, *k*_1_ = 0.2, *k*_2_ = 0.1, producing a Mexican-hat connectivity profile, with a constant input bias of −2.85 applied to all units. Velocity input asymmetrically modulated the gain of each subpopulation, enabling path integration. Specifically, for velocity *v*(*t*), the left and right population gains were scaled by (1 − *g*_*v*_ · *v*(*t*)) and (1 + *g*_*v*_ · *v*(*t*)), respectively, with velocity gain *g*_*v*_ = 0.1. Network dynamics followed a linear-nonlinear-Poisson dynamics with synaptic time constant *τ* = 0.01. The synaptic variable *s* decayed exponentially:

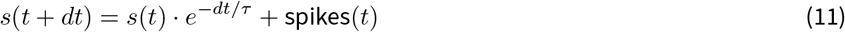

where a spike was generated by sampling a Poisson random variable based on the current firing rate and velocity input:

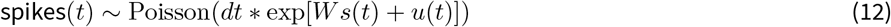

Angular velocity input was generated as an Ornstein-Uhlenbeck process with time constant *τ*_vel_ = 0.2 and standard deviation *σ*_vel_ = 1.0 to mimic naturalistic head movements. Simulations used Euler integration with time step *dt* = 0.001 for a duration of 10000 timesteps per trial, with *n*_ic_ = 20 initial conditions. Spike counts were saved every 10 integration steps (*dt* = 0.01) and binned to *dt* = 0.1, matching the 100ms bins of the experimental data.

#### B.1.2 Neural cut ring attractor generation

To generate a line / cut ring attractor with matched biological complexity to the ring attractor network, we modified the connectivity structure using a topology-breaking parameter *α* ∈ [0, 1]. The weight matrix was modified as follows:

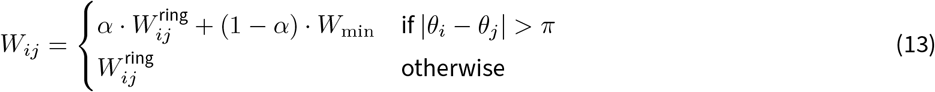

where 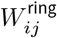 is the standard ring attractor connectivity, *θ*_*i*_ and *θ*_*j*_ are the preferred head directions of neurons *i* and *j*, and 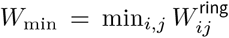 is the most inhibitory weight in the connectivity profile. When *α* = 1, the connectivity is unchanged and the network implements a ring attractor with continuous angular dynamics. When *α* = 0, connections between neurons on opposite sides of the ring (angular separation > *π*) are replaced with maximal inhibition. This creates a break in the ring topology: the network can no longer smoothly transition activity across the inhibitory barrier, effectively confining dynamics to a bounded arc, thereby producing a line attractor. This manipulation preserves all other aspects of the network architecture (number of neurons, local connectivity profile, velocity modulation, noise statistics, and temporal dynamics), isolating the topological difference between ring and line attractors while controlling for model complexity. We simulated these neural line attractors in the exact same manner as the neural rings.

#### B.1.3 Simple polar ring model

To provide controlled comparisons, we implemented a minimal stochastic attractor model in polar coordinates (*r, θ*) using Euler-Maruyama integration with multiplicative noise in the radial direction:

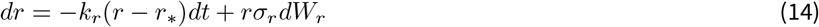

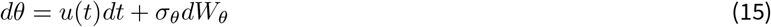

where *dW* is a standard Wiener process, *r*_∗_ = 1 was the stable radius, *k*_*r*_ = 1*/τ*_*r*_ = 10 governed radial relaxation dynamics, and *u*(*t*) was an external angular velocity input. This produced a stable circular manifold with diffusive angular dynamics. To implement the line attractor, we modified the ring attractor to create attractive boundary conditions at two points of the ring:

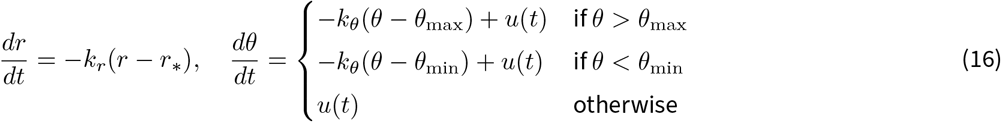

where *θ*_max_ = −*θ*_min_ = *π* · *c* defined angular boundaries determined by the coverage parameter *c* = 0.75 (3-quarters-circle), and *k*_*θ*_ = 10 controlled the stiffness of the boundary restoring force. This produces a curved line segment rather than a closed ring. For both models, we simulated *n* = 60 trajectories from random initial conditions near the attractor, each of duration *T* = 10 s with time step Δ*t* = 0.01 s. States were mapped to Cartesian coordinates via *x* = *r* cos(*θ*), *y* = *r* sin(*θ*) for visualization and analysis. In Fig. S1 we modulated *c* from 0.9 to 0.99.

### B.2 Synthetic system classes

To benchmark DSA against alternative dynamical and geometric metrics (Fig. 2c-f), we constructed a dataset of 6 synthetic model systems, with 5 independent samplings from each based on random seeds for noise, yielding *N* = 30 systems in total. The 6 classes are the Cartesian product of three dynamics types (ring attractor, line attractor, limit cycle) and two model classes (simple polar and neural-network rings). Simple systems were simulated with 60 initial conditions × 1000 timesteps and NN systems were simulated for 20 initial conditions and 1000 timesteps.

For each comparison method (DSA variants, MMD, nonlinear CKA on trajectories, Procrustes on trajectories, Wasserstein Procrustes on point clouds, TDA, MARBLE), we computed the distance matrix between each dataset and quantified its clustering and class-separation ability via cross-validated linear decoding and silhouette score. To estimate variability, we bootstrapped the data itself by subsampling initial conditions at a fraction of 0.7 within each system, then refitting each distance matrix and recomputing all scores; this was repeated for 10 bootstrap resamples. Unless stated otherwise, all metrics operate on the full datasets.

- **Silhouette score**. Given the distance matrix 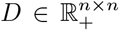 and labels y, we compute the silhouette score via *scikit-learn*: silhouette 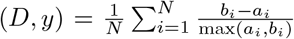 where *a*_*i*_ is the mean intra-label distance and *b*_*i*_ is the mean inter-label distance. The score ranges from −1 to 1, with 1 indicating that all elements of each label cluster perfectly together. A score of 0 typically means that all classes are overlapping.
- **Linear decoding** A linear support vector classifier is fit directly to the rows of *D*. We report the five-fold cross-validation accuracy.

### B.3 Distance methods

- **MMD** Maximum Mean-discrepancy in RFF space (*Gretton et al., 2012*),

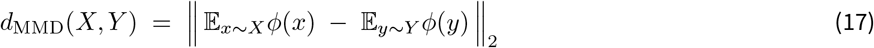

RFF features were used with dimension 5000 and *γ* = 1.0*/*(2*σ*^2^) with *σ*^2^ the empirical variance of the flattened data.
- **Nonlinear CKA** On RFF features Φ_*X*_ ∈ ℝ^*m×D*^ obtained after density-weighted subsampling, with centered features 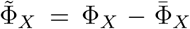, define the Frobenius inner product 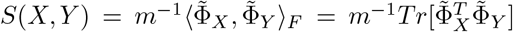. The CKA distance

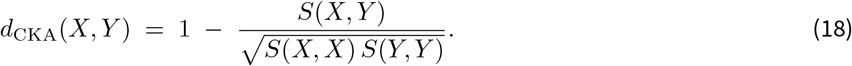

RFF features were defined as in MMD. For trajectory geometry comparisons, *n* initial conditions and *t* timepoints were flattened into the *m* dimension. When datasets had different numbers of timepoints, the larger dataset was truncated.
- **Procrustes** As is standard, we mean-centered and normalized each data array before applying Procrustes:

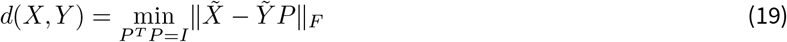

We categorize this method as a trajectory geometry method because it entails a fixed alignment between points to be compared, which can be used in practice to compare trajectories by aligning points in each dataset by time. When datasets had different numbers of timepoints, the larger dataset was truncated.
- **Wasserstein Procrustes (WP, *Grave et al***., ***2019*)** WP alternates between (i) a hard assignment *π* extracted from the discrete optimal transport plan

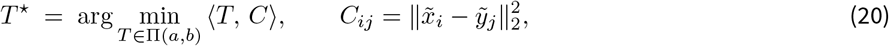

between centered and normalized point clouds of 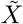 and 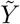, with uniform marginals *a, b*, and (ii) an orthogonal Procrustes alignment

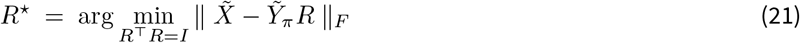

where 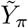 denotes the transported points. After convergence of the Procrustes score we report 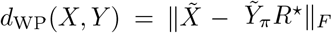. We iterated the above up to 10 maximum times, with tolerance 10^−4^, but the procedure typically stopped well before the maximum. When datasets had different sizes, the transport plan is restricted to a hard permutation for step (ii). Following this, optimal alignment *R* was applied to every point and the transport plan was recomputed on the full dataset each time. WP is a point cloud geometric comparison because it allows for variable, non-fixed alignment between data points, thereby treating it as an unordered set unlike standard Procrustes. Because the alternation is a coordinate descent that converges only to a local optimum and the result depends on initialization, WP is not guaranteed to be symmetric or to satisfy the triangle inequality; we report it as a dissimilarity score rather than a metric.
- **Topological Distance (TDA)** Topological Data Analysis is notoriously slow on large datasets (on our setup a TDA computation on 8000 data points took 40 minutes), so we subsampled our data based on their inverse density with a Kernel Density Estimator (oversampling low density regions in order to capture the full shape of the data), then ran a Vietoris– Rips filtration to obtain persistence diagrams *D*^(0)^, *D*^(1)^, *D*^(2)^. Counting features whose persistence exceeds a threshold *τ*_*i*_, defined as a fixed multiple of the data standard deviation for each Betti number (0.9, 0.4, 0.4, selected by direct visualization of the birth–death distribution), gives the Betti vector. We used the *ripser* and *gudhi* packages in Python. We confirmed that our subsampling did not destroy the expected results for each dataset. On persistence diagrams *D*^(0,1,2)^, which denote the birth and death of homological features (*b, d*) based on filtration length, we compute the Betti number vector (*Ghrist, 2008*):

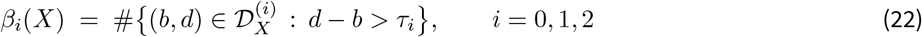

where *β*_0_ denotes the number of connected components, *β*_1_ denotes the number of 1-dimensional holes, and *β*_2_ denotes the number of 2-dimensional holes. The distance is then defined as:

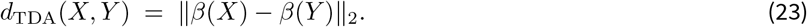
- **kernel DSA** We embedded our data with a Cauchy kernel of *γ* = 0.1, and then applied DSA with rank 6 and 1 delay after sweeping hyperparameters based on ResDMD and AIC. We chose 500 random features as it approximated the full kernel with high accuracy while maintaining computational efficiency.
- **MARBLE** MAnifold Representation Basis LEarning (*Gosztolai et al., 2025*) is an unsupervised neural network that jointly embeds local flow fields and states into a unified embedding space across multiple datasets, which enables comparison across datasets based on the distribution of local dynamical structure. We used MARBLE on our systems with default hyperparameters (following a sweep), which the original work also found to be sufficient across many diverse systems with similarity to ours. To ensure that our results were robust, we followed their application in ensuring convergence and stability by recomputing across multiple random seeds. Latent dimension was set to 6 to match the DSA rank for comparable evaluation. We chose a Multilayer Perceptron architecture with two layers of size 64 to balance between expressiveness and efficiency. MARBLE also subsamples point clouds using farthest point sampling, which we left at the default value of 0.03. Without this parameter, we found that the method was extremely memory and compute intensive, with a single graph construction taking over 45 minutes in our setup. We trained for 200 epochs to ensure convergence.

### B.4 Lorenz analysis

We simulated the Lorenz attractor

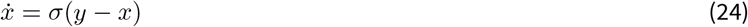

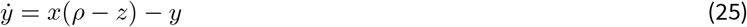

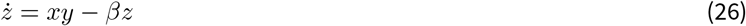

we fixed *σ* = 10, *β* = 8*/*3.

The LLE is computed by evolving a small perturbation to an initial condition *X*_0_ + *δX* over *t* timesteps, yielding new perturbed state 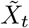 and computing the average log-deviation:

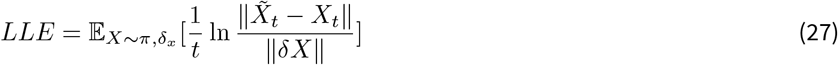

where *π* indicates the invariant distribution of the attractor, and *δX* was sampled from a 3D Gaussian distribution with variance 10^−9^. We found that setting *t* = 10 was sufficient for approximating the LLE (e.g. for default *ρ* = 28, we found the LLE was 0.904, and it is known to be 0.905 (*Alligood et al., 1996*)). We also reported the standard deviation of the LLE across 5 different random samples of the above computation.

We simulated each model with *dt* = 0.001 using a random initial condition generated by running the model for 300 timesteps, then generated 40, 000 timesteps for each initial condition. We then subsampled the data by a factor of 10 and added Gaussian noise with standard deviation 0.1 to each dimension before whitening the data.

For DSA, we utilized 15 delays and rank 6. Procrustes is performed for trajectory geometry comparisons. TDA, MMD, and MARBLE were all computed using the same hyperparameters as in the ring attractor analysis (Methods B.3).

## C Head direction circuit analysis with DSA

### C.1 Experimental data

We analyzed extracellular recordings from head direction cells in the anterodorsal thalamic nucleus (ADn) of freely behaving mice during waking exploration, REM sleep, and Slow Wave Sleep (SWS). Neural data were obtained from a previously published dataset (*Peyrache et al., 2015*). Following *Chaudhuri et al., 2019*, spike trains were smoothed using a Gaussian kernel (*σ* = 100 ms), square-root transformed to reduce variance, and binned at 100 ms resolution. Population activity across all three states was first projected with PCA to the top dimensions that explained 95% of the total variance, then embedded using Isomap to 8 dimensions (using a hyperparameter of 5 nearest neighbors). Joint embedding ensured that all states shared a common coordinate system, enabling direct comparison of manifold structure and dynamics. To share dimensionality across all datasets and models, DSA was applied to the top two dimensions (noting that delay embedding recovers unobserved dimensions). Data were z-scored prior to analysis. We utilized all possible datasets for each animal and day, yielding 31 datasets for 5 animals across all brain states. Preliminary analysis identified 1 dataset as an outlier due to erroneous Isomap embedding (Mouse17-130125) which was held out of all primary analyses.

### C.2 Statistical analysis

DSA was applied to the combined dataset comprising wake ADn neural recordings, ring attractor simulations (both simple stochastic and neural network), and line attractor simulations (both simple stochastic and neural network). This produced a pairwise Wasserstein distance matrix *D* ∈ ℝ^*n*×*n*^ where *n* is the total number of datasets. To visualize the structure of dynamical similarities, we applied PCA to this distance matrix.

#### C.2.1 Paired comparison of ring vs. line attractors

To test whether wake neural data dynamics were more similar to ring attractors than to cut ring attractors, we computed for each wake recording session *i* the mean DSA distance to all ring model instances and to all cut ring model instances:

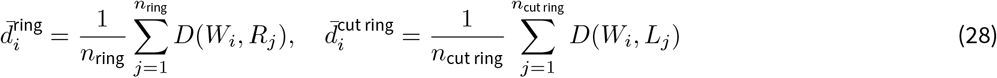

where *D*(*W*_*i*_, *R*_*j*_) denotes the DSA distance between wake session *i* and ring model instance *j*. The dataset for this comparison comprised *N*_*W*_ = 12 wake recording sessions and 20 model instances (5 simple rings, 5 NN rings, 5 simple cut rings, 5 NN cut rings), yielding 12 × 5 = 60 wake→ring and 12 × 5 = 60 wake→cut ring distances per model class. For each wake session, we averaged the 5 wake → ring (or wake→cut ring) distances within a model class to obtain a single paired value, giving *N* = 12 paired observations per test. We then tested the hypothesis that wake dynamics are closer to ring attractors (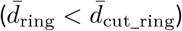) using the one-sided Wilcoxon signed-rank test, which is appropriate for paired comparisons and does not assume normality. The test is one-sided because the ring attractor is the established model of the HD circuit, making *d*_ring_ < *d*_cut_ring_ an a priori prediction rather than a post hoc choice. The test was performed separately for simple stochastic models and for neural network models. As a reference, we computed the inter-quartile interval of cross-wake session DSA distances from the 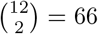 unique wake-wake pairs to contextualize the magnitude of wake-to-model distances.

#### C.2.2 Hyperparameter robustness analysis

To ensure that our conclusions were not artifacts of particular DSA hyperparameter choices, we performed a systematic sweep over the number of time delays *d* ∈ [2, 40] and rank *k* ∈ [6, 30]. For each (*d, k*) combination, we recomputed the DSA distance matrix and performed statistical testing.

To filter out hyperparameter settings where DSA failed to meaningfully distinguish between different dynamical systems, we applied a quality control criterion: we required that across-system distances (ring-to-cut ring) be significantly different from within-system distances (wake-wake, ring-ring, and cut ring-cut ring combined). Specifically, we computed:

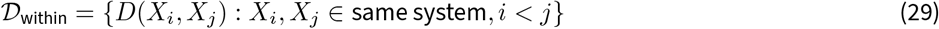

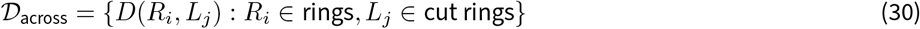

and tested whether these distributions differed using the Mann-Whitney U test, as each dataset was sampled independently. Hyperparameter settings where *p*_across vs. within_ ≥ 0.05*/*3 (Bonferroni-correction) were excluded, as they indicated that DSA could not reliably distinguish ring from cut ring dynamics at that setting.

For the remaining valid hyperparameter configurations, we performed the paired Wilcoxon signed-rank test comparing 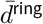 vs. 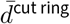 across wake sessions. We applied Bonferroni correction for multiple comparisons, using a significance threshold of *α* = 0.05*/*2 to account for testing both simple and neural network model families. Results were visualized as cumulative distribution functions (CDFs) of the Wilcoxon test statistic across the hyperparameter sweep, with the fraction of significant tests reported separately for simple and neural network models.

### C.3 Cross-state DSA: Wake, REM, SWS

#### C.3.1 Multi-timescale DSA

To probe dynamical similarity across different temporal scales, we varied the DMD prediction horizon parameter *s* ∈ [1, 80]. This parameter controls how far into the future the fitted Koopman Operator is used to predict state evolution:

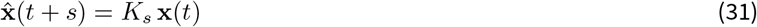

where *K* is the Koopman Operator and *s* is the number of steps ahead. At small *s*, DSA emphasizes fast dynamical modes; at large *s*, fast modes are averaged out and slow, persistent modes dominate the comparison. For each value of *s*, we computed the DSA distance matrix using *d* = 20 time delays and rank *k* = 6, then extracted pairwise distances:

1. wake-wake: DSA distance between different wake recording sessions (inter-session variability)
2. wake-REM: DSA distance from wake sessions to REM sessions
3. wake-SWS: DSA distance from wake sessions to SWS sessions
4. wake-shuffled: DSA distance from wake to temporally-shuffled wake (baseline control)

#### C.3.2 Statistical analysis

For each prediction horizon *s*, we tested whether wake-REM distances were smaller than wake-SWS distances using the one-sided Wilcoxon signed-rank test. The dataset comprised *N*_*W*_ = 12 wake, *N*_*R*_ = 9 REM, and *N*_*S*_ = 9 SWS recording sessions. Specifically, for each wake session *i*, we computed the average distance to each non-wake state:

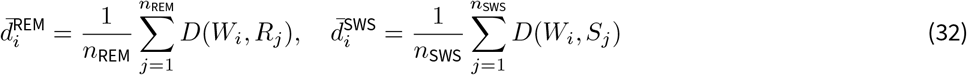

giving 12 × 9 = 108 wake–REM and 12 × 9 = 108 wake–SWS distances total, which were averaged per wake session to yield *N* = 12 paired observations for the test. As an inter-animal reference, we also computed wake–wake distances; averaging each wake session’s 11 distances to other wake sessions gave *N* = 12 inter-animal reference values, against which 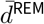 was tested. We averaged across sessions rather than treating each pairwise distance as independent because the experimental sampling unit is the recording session, not the pairwise comparison. The direction (REM being closer to wake than SWS) was specified a priori from prior reports (*Chaudhuri et al., 2019*). Multiple comparisons across prediction horizons were corrected using Bonferroni correction accounting for the primary comparisons: wake vs. wake, REM vs. wake, and SWS vs. wake.

## D Macaque reaching neural data analysis

The data used in this study originates from the publicly available dataset collected by Perich, Miller, and colleagues (*Perich et al., 2025*), accessed from the DANDI archive. The dataset includes multielectrode recordings from three male rhesus macaques (monkeys C, M, and J) performing a Center Out (CO) reaching task as well as a Random Target (RT) task.

### D.1 Subjects and behavioral tasks

We analyzed neural recordings from three male macaques (*Macaca mulatta*: monkeys C, M, J) performing two upper-limb reaching tasks, as previously described (*Safaie et al., 2023; Perich et al., 2025*). In the Center Out (CO) task, monkeys held a cursor at a central target, received a go cue after a variable-length delay, and subsequently reached to one of eight equidistant peripheral targets. In the Random Target (RT) task, monkeys made four consecutive reaches to randomly placed targets within a 20 × 20 cm workspace; each target’s position was selected by drawing a distance (5–15 cm) and an angle (0–360^*о*^) relative to the preceding target. Multi-unit activity was recorded from chronically implanted electrode arrays in primary motor cortex (M1) and dorsal premotor cortex (PMd). Recordings comprised 101 sessions across animals (monkey C: 70 sessions, October 2013– October 2016; monkey M: 28 sessions, January 2014–June 2015; monkey J: 3 sessions, April 2016). After excluding sessions with fewer than 10 simultaneously recorded M1 neurons with a firing rate > 0.1 Hz on average, 16 RT, 70 CO, and 59 control sessions were retained for the main analysis.

### D.2 Neural data preprocessing

Spike times were binned at 10ms resolution. Neurons with mean firing rates below 0.1 Hz (averaged across all trials and time steps) were excluded. Spike counts were square-root transformed for variance stabilization and smoothed with a non-causal Gaussian kernel (*σ* = 100 ms). Firing rates were soft-normalized by dividing each neuron’s activity by its standard deviation plus a constant (*c* = 5.0) to prevent low-firing neurons from dominating the population signal. Dimensionality was reduced to 10 principal components via condition-averaged PCA: PCA was fit on trial-averaged trajectories (averaged across trials within each reach direction), and all individual trials were then projected onto the resulting basis. PCA projections were subsequently *z*-scored across trials.

For movement epochs, neural trajectories were extracted from 50ms before to 450ms after movement onset (50 time steps). Movement onset was detected as the first time point at which hand velocity magnitude exceeded a threshold defined as 10% of the peak velocity within a trial. For the preparatory (control) epoch, trajectories were extracted from the 900ms preceding the go cue and truncated to 500ms (50 time steps), capturing the delay-period preparatory activity.

### D.3 Dynamical Similarity Analysis

We used Dynamical Similarity Analysis (DSA) to quantify pairwise dynamical similarity between recording sessions. Because M1 is input-driven, we applied SubspaceDMDc (*Huang et al., 2025*) which estimates the DMD in a latent space in the presence of input. Trial-related inputs were provided to the SubspaceDMDc model to account for trial-varying movement goals. For all trials, the target direction was encoded as the 2D hand position at the end of the reach, plus a constant term. This input was held constant across time steps, and one SubspaceDMDc model was fit per session. Control epochs used the same encoding as their corresponding CO movement sessions.

To assess robustness to hyperparameter selection, DSA was computed over a grid of hyperparameters: ranks ∈ {8, 10, …, 28} (11 values) for kinematics and ∈ {18, 20, …, 38} for neural data, and delay embeddings ∈ {10, 11, …, 16} (7 values) for each, chosen based on the hyperparameter sweep (Fig. S8). If the rank parameter was greater than the delay size multiplied by the data dimensionality, that hyperparameter set was removed, resulting in 67 hyperparameter settings for the kinematics data and 77 hyperparameter settings for the neural data. In the main results, we chose a rank of 18 for the kinematic data and a rank of 26 for the neural data, and a delay of 14 for both.

### D.4 Statistics

#### D.4.1 Statistical testing

We computed a one-sided label permutation test to compare the DSA distances across groups (within-monkey versus cross-monkey, within-task versus cross-task). The test statistic was defined as the difference of medians between cross-group and within-group pairwise distances; medians were chosen over means to prevent outlier distances from having an outsized effect. Labels were randomly permuted 10,000 times and the cross-group versus within-group null distance was computed, then the *p*-value was computed as the fraction of samples that exceeded the test statistic.

#### D.4.2 Equivalence testing

To assess whether pairwise DSA distances were statistically equivalent across experimental conditions (e.g., within-monkey versus cross-monkey, or within-task versus cross-task), we additionally used a one-sided bootstrap equivalence test on the difference between medians as in the corresponding statistical tests. For each comparison, 10,000 bootstrap resamples were drawn with replacement independently from each group (preserving original sample sizes). This test utilized the null hypothesis that the median difference exceeded an equivalence margin Δ equal to the standard deviation of the DSA distances for the within-group reference distribution, and the *p*-value was computed as the fraction of bootstrap resamples that the test statistic exceeded the equivalence margin. For within-vs-cross-task comparisons, the minimum of the CO and RT margins was used as a conservative bound.

#### D.4.3 Test statistics and effect sizes

For every statistical test we report the test statistic, the *p*-value (or floor, if it is achieved), and effect size. Degrees of freedom are not defined for our nonparametric rank- or permutation-tests. For comparisons between two independent groups, on which we apply permutation or Mann–Whitney *U*-tests, we report Cliff’s 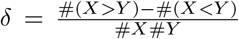, where *X* denotes the cross-group distances, and *Y* the corresponding within-group distances. *δ* ranges from −1, 1, with *δ* = 1 indicating that every cross-group distance is greater than within-group distances. This statistic is a linear transformation of the Mann–Whitney statistic, in that *δ* = 2*U/*#*X*#*Y* − 1, making these statistics directly comparable across our different tests.

For the paired comparisons in the head direction analyses, we report the Wilcoxon signed-rank statistic *W* with the Cliff’s *δ*. At *n* = 12 in the Wake regime the smallest attainable one-sided *p*-value is 2^−12^ = 2.4414 × 10^−4^.

For the equivalence tests we report the median difference Δ_med_ as the test statistic, alongside the equivalence margin *σ*_*w*_ and their ratio, which is a standardized effect size. Values below 1 indicate an observed effect smaller than the margin of equivalence (and therefore equivalence is attained at this threshold).

For the longitudinal comparisons we report the OLS slope as the test statistic with its bootstrapped 95% CI, together with the *R*^2^.

### D.5 Eigenvalue Kernel Density Estimate difference

We computed the eigenvalues across all sessions for each task, then applied a Kernel Density Estimate (KDE) with bandwidth equal to the data variance to each model separately. On a 300 × 300 grid within the complex plane spanning [−1, 1] in the real and imaginary directions, we took the pointwise difference of the RT and the CO KDEs. We then thresholded this difference by visualizing only points that were outside the 95% quantile of a null KDE difference distribution that we constructed by shuffling task labels and recomputing the KDE difference. We finally rescaled by the standard deviation of the null distribution.

### D.6 Cross-day stability of dynamics

To assess the temporal stability of neural dynamics across recording sessions, we computed the DSA distance for all pairs of sessions within a single animal and plotted it against the number of days elapsed between recordings. Session pairs were grouped by type: within-task (both sessions from the same task, either RT vs RT or CO vs CO) and cross-task (one RT session paired with one CO session). For each group, a line of best fit was computed via least-squares linear regression. Statistical significance of the slope being of magnitude greater than zero (one-sided test) was assessed with a session-date grouped permutation test for robustness. Results were aggregated across monkeys C and M, as they had more than 10 valid sessions (see above).

### D.7 Robustness across hyperparameters

To assess whether our conclusions were robust to DSA hyperparameters, we ran aggregate robustness analyses on the full sweep of DSA hyperparameters. For all permutation and equivalence tests, we ran 10,000 simulations.

We tested the one-sided hypothesis that the median DSA distance was significantly greater in the cross-task (cross-animal) comparisons than within-task (within-animal):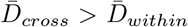. We used a label permutation test on the distribution of median 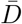 over all individual DSA hyperparameters. We also performed aggregate equivalence tests on the distribution of median distances using bootstrap resampling, taking the equivalence threshold as the standard deviation of the within-group distribution as in the individual hyperparameter comparisons. We chose to shuffle labels across medians rather than individual distances within hyperparameters because the same distance (e.g. between monkey M day 1 and monkey C day 2) was correlated across hyperparameters.

To visually compare the aggregate cross-hyperparameter kinematic and neural DSA distances directly, we centered and standardized the distances based on the median within-task (as a noise floor) and cross-epoch scores (as a noise ceiling) for each hyperparameter:

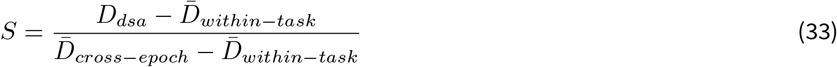

Lastly, we also reported the statistics of each individual statistical and equivalence test for all hyperparameters, including the effect size, the fraction of tests that were significant, and the distribution of p-values.

#### D.7.1 Longitudinal robustness testing

For each dataset type (neural, kinematic) and task comparison type (within-CO, within-RT, cross-task), we tested the one-sided hypothesis that Δ*d*_*DSA*_*/day* > 0 using grouped session-date permutation tests as in Sec. D.6. We defined our test statistic as the median slope across all DSA hyperparameters. Each sample of the null distribution was created by shuffling dates of each session within each hyperparameter, computing the Δ*DSA/day* slope, and taking the median over all hyperparameters. Our p-value was then computed as the fraction of shuffles that yielded slopes greater than or equal to the test statistic. We chose to shuffle dates of each session within each hyperparameter setting as these were the fundamental units of analysis, rather than the hyperparameter settings.

### D.8 Coupling analysis

#### D.8.1 Synthetic validation of coupling detection

We validated the sliding-window DSA coupling framework on two synthetic systems with known coupling structure. An autonomous 3D Rössler system (parameters *a* = 0.2, *b* = 0.2, *c* = 5.7) was coupled to a Lorenz system (3D, parameters *σ* = 10, *ρ* = 28, *β* = 8*/*3) via a quadratic coupling term (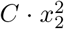, *C* = 3, added into the Lorenz 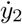 equation), with coupling offset at *t*^∗^ = 10000 (5 trials, 20,000 simulation steps in total with Δ*t* = 0.01 and Euler integration). Sliding-window DSA was applied with window size 2000, stride 2000, rank 12, and 10 delays.

#### D.8.2 Cross-brain region comparison

To quantify time-varying dynamical coupling between PMd and M1, we applied DSA in a sliding-window framework. For each of 14 valid paired PMd–M1 recording sessions, neural trajectories were segmented into overlapping temporal windows (300ms width, 50ms step) spanning 900ms before to 900ms after movement onset (31 windows per session). Within each window, DSA with SubspaceDMDc was utilized to compare the PMd and M1 population trajectories. Control inputs encoded target direction in the primary analyses, held constant across time. To ensure robustness, results were averaged across 8 hyperparameter configurations (ranks ∈ {8, 10}, delays ∈ {4, 5, 6, 7}). Larger delays were invalid due to the small window size of 30 timepoints, and adding larger ranks resulted in excess noise for the same reason.

As a baseline, within-region DSA was computed separately for M1 and PMd by comparing across sessions within each window, and averaging across both brain regions. We evaluated the significance of cross-region versus within-region DSA distance using one-sided Mann-Whitney *U* test at each time window. Bonferroni-correcting for 31 simultaneous comparisons changes the threshold of 0.05 to 0.00161.

To characterize the spectral properties of the fitted dynamics, dynamics matrices (**A**) were extracted from the SubspaceDMDc model at each window. The maximum absolute imaginary eigenvalue component (*ω*_max_, reflecting oscillation frequency of the dynamics) of each dynamics matrix for each region was reported as a function of time relative to movement onset.

## E DSA regularization methods

### E.1 RNN architecture and task definition

We simulated a continuous-time vanilla recurrent neural network (RNN) via Euler integration:

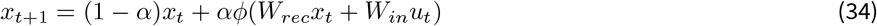

where *x*_*t*_ ∈ ℝ ^D^ is the hidden state with *D* = 64, *u*_*t*_ ∈ ℝ is the external control input, *α* = 0.2 is the discrete integration step, and *ϕ*(·) is the nonlinear activation function (we used ReLU max (0, *x*)). The network output *y*_*t*_ = *W*_*out*_*x*_*t*_ ∈ ℝ is computed via a linear readout.

The networks were trained on a continuous sequence integration task over sequences of length *T* = 50. Inputs *u*_*t*_ comprise piecewise-constant step functions with amplitude uniformly sampled between [−1, 1] and randomized transition times as well as additive Gaussian noise with *σ* = 0.25. The target output at each time *y*^t∗^ was the cumulative integral of the prior input sequence:

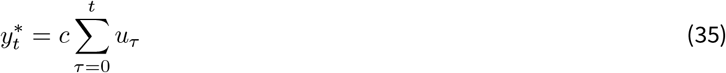

where *c* = 0.1 is a scaling constant. The primary task objective was the mean squared error (MSE),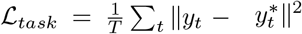.

All models were trained for 10,000 epochs with batch size 32 using the AdamW optimizer (learning rate 5 × 10^−4^, weight decay 10^−4^, *β*_1_ = 0.9, *β*_2_ = 0.999), with gradient clipping at norm 2 and recurrent weight matrix initialized from a standard Gaussian. Networks were initialized from the same uniform random seed (seed 42) across iterations to isolate the effect of the regularizer. The unregularized baseline ensemble (used in the MDS comparison) was generated by re-running iteration 0 with different random seeds, as well as sweeping across multiple hyperparameters: learning rate 5*e*−4, 5*e*−3, 5*e*−2, recurrent dimensionality 16, 32, 64, 128, batch size 16, 32, 64, 128, weight decay 1*e* − 4, 1*e* − 3, 1*e* − 2, and standard deviation of initial recurrent weight matrix 0.75, 1, 1.5.

### E.2 Differentiable DMD, DMDc, and DSA regularization

Standard backpropagation through the Singular Value Decomposition (SVD) fails numerically for near-degenerate singular values due to gradient terms proportional to 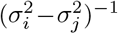. To enable end-to-end optimization of the manifold dynamics, we utilized gradient-detached SVD bases and applied them as differentiable projections.

#### E.2.1 Differentiable DMDc

We incorporated the control input into the operator estimation via Dynamic Mode Decomposition with control (DMDc). We constructed delay-embedded Hankel matrices for both the states *H*_*x*_ and inputs *H*_*u*_ using 5 delays. Detached SVDs were computed as 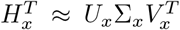 and 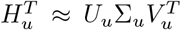. To maintain the autograd graph, the right singular bases were computed differentiably as:

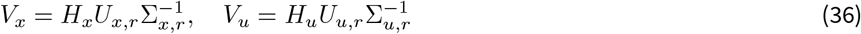

where the rank *r* was fixed at 12, and delays were chosen to be 5. Letting *V*_*x*−_ and *V*_*x*+_ denote the time-shifted state bases, and *V*_*u*−_ the input basis, we define the concatenated basis 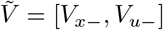. The state transition operator *A*_*v*_ and input operator *B*_*v*_ were jointly estimated via differentiable ridge regression:

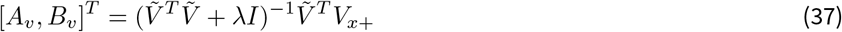

where *λ* is a small ridge constant (10^−5^) for numerical stability. DSA distances were computed on the hidden-state operators (not the readout) and re-fit every epoch. To improve noise robustness and stability in the comparison against baselines, we reduced to rank 6 in our subsequent analyses.

#### E.2.2 Differentiable DSA distance

The dynamical similarity between two operators was computed via the entropy-regularized Wasserstein distance between their spectra. Given the complex eigenvalues Λ_*A*_ and Λ_*B*_, mapped to 2D coordinates *x*_*i*_, *y*_*j*_ ∈ ℝ ^2^, we constructed the pairwise squared distance matrix *C*_*ij*_ = ∥*x*_*i*_ − *y*_*j*_∥^2^. *d*_∂*DSA*_(*A, B*) is given by the Sinkhorn distance (*Cuturi, 2013*):

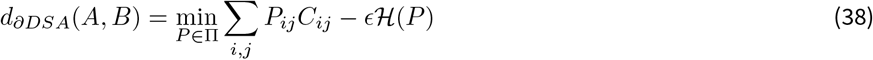

where ℋ (*P*) is the Shannon entropy, *ϵ* = 0.05, and Π is the set of transport maps with uniform marginals.

### E.3 Iterative regularization pipeline

Let *θ*^(*k*)^ define the network parameters at iteration *k*. The base iteration *k* = 0 was trained strictly on ℒ_*task*_ to establish an unregularized manifold of solutions. For iterations *k* ≥ 1, the network was initialized from the same uniform seed and trained with a repelling DSA penalty evaluated against the last-epoch DMDc operators *A*^(*i*)^ of all previously-trained models *i* < *k*. Each *A*^(*i*)^ was fit to the hidden-state trajectories of network *i* over *n*_ref_ = 64 task-matched reference trials, and recomputed once per epoch on the same input batch used by the learner so that operator estimates are not biased by input-distribution mismatch. The combined loss for *k* ≥ 1 is:

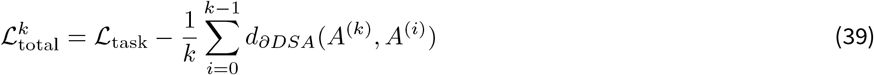

so that maximizing *d*_∂*DSA*_ minimizes the loss.

### E.4 Analyses

#### E.4.1 Relaxation analysis

We isolated the autonomous dynamics by driving networks with task-matched inputs for *T*_*drive*_ = 25, followed by *T*_*coast*_ = 400 steps of zero input across *N* = 1000 trials. The hidden states *H* ∈ ℋ^*N*×*T*×*D*^ were projected via PCA (after collapsing the first two dimensions) to geometrically evaluate the transient and asymptotic dynamics.

#### E.4.2 Analytic baselines and references

Regularized solutions were compared against baseline RNNs trained purely on ℒ_*task*_ with variable random seeds. We constructed two ground-truth linear dynamical solutions for comparison:

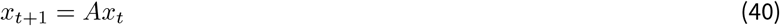

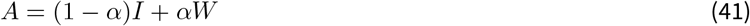

- *Line Attractor (LA):* A strictly normal system characterized by a single integrating mode (largest eigenvalue *λ*_1_ of *A, λ*_1_ = 1 − 10^−3^) and uniformly decaying orthogonal dimensions (*λ*_*i*>1_ = 0.2), with *b*_0_ = 1.0, *b*_*i*>0_ = 0.
- *Feedforward Chain (FF):* A highly non-normal shift operator matrix (*W*_*i,i*+1_ = 1) representing a pure delay-line, with eigenvalues clustered at 0 and input isolated to the terminal node (*b*_*N*−1_ = 1). Eigenvalues of *A* are therefore at 1 − *α* = 0.8.

#### E.4.3 Jacobian analysis

To dissect the globally nonlinear RNN into its constituent linear regimes, we exploit the piecewise-linear structure of the ReLU activation. During integration (on 128 random trials for 50 timesteps each), the state space is partitioned by the active pre-activation signs *m*_*t*_ = I(*W*_*rec*_*x*_*t*_ + *W*_*in*_*u*_*t*_ > 0).

For each uniquely visited partition *k*, we define the local Jacobian:

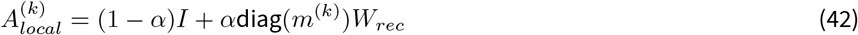

The dynamics of each partition were characterized by three quantities:

- *Maximum real component of all eigenvalues:* max Re(*λ*_*i*_*), quant*ifying the dominant timescale.
- Second largest real component eigenvalue, which is used to determine the dimensionality of the integrating space. If significantly less than 1, this suggests that the integrator is a line.
- *Non-normality* Computed as 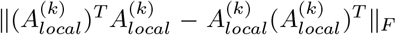. This metric measures the degree of transient non-normal amplification within the local regime (*Trefethen and Embree, 2005*). Note that the normalization constant is not included here as all matrices are equivalently sized.

We also computed and plotted the distribution of all Jacobian eigenvalues across every partition.

## F Supplementary Information (SI)

### F.1 Related work

#### F.1.1 Neural networks (embedding models)

Neural network-based embedding models learn representations of system dynamics, or flow fields, from data (*Gosztolai et al., 2025*). While these models have been applied to neural data and can handle unaligned time series, they come with significant drawbacks: high technical, computational, and data complexity, requiring large datasets and substantial computing resources. Their theoretical grounding for similarity is only partial, and the resulting embedding spaces have low interpretability as they are not proper metric spaces.

#### F.1.2 Neural networks (generative models)

Generative neural network models learn a latent space in order to predict the dynamics of multiple systems (*Cotler et al., 2023; Brenner et al., 2025*). Like other deep learning methods, they require significant technical expertise, computational resources, and large amounts of data. Their primary weaknesses are a lack of theoretical grounding for how they measure similarity and low interpretability (they likewise do not learn proper metric spaces). Crucially, these specific models have not yet been successfully applied to neural data.

#### F.1.3 Neural networks (alignment models)

Another class of neural network models seeks to directly learn a conjugacy map that aligns different systems or prototypes (*Chen et al., 2024; Friedman et al., 2025*). These methods benefit from strong theoretical motivation. However, they are much more computationally complex than other methods as they require training a new neural network for each *comparison*, which scales quadratically in the number of comparisons. Likewise, they are unable to handle unaligned data, with prior work in this family primarily resorting to comparing or aligning to a computational model on which the dynamics are computable.

#### F.1.4 Shape metrics

Shape-based metrics directly compare the geometry of system trajectories (*Nejatbakhsh et al., 2025; Williams et al., 2021; Bar-bosa et al., 2025*) via timewise alignment of two ℝ^*T*×*d*^ data matrices (T timesteps, d dimensions). If there are multiple trials, this dimension is typically flattened or averaged over. The main advantages of this approach are its low technical complexity, low data requirements, and high interpretability. The primary limitations are a lack of theoretical grounding for dynamical similarity and an inability to handle unaligned data.

#### F.1.5 Koopman-based comparisons

Koopman operator or Dynamic Mode Decomposition (DMD) methods approximate a linear operator from trajectory data, allowing for direct comparison (*Ostrow et al., 2023; Redman et al., 2024; Huang et al., 2025; Cohen et al., 2023; Germain et al., 2026*). These approaches are advantageous due to their strong theoretical grounding, low computational complexity, and high interpretability. Furthermore, they are capable of handling unaligned data. A range of DMD methods have been designed for different conditions, which can be used to leverage structure in the data for improved fitting. For example, *Cohen et al., 2023* utilized Physics-Informed DMD (which identifies the optimal linear operator constrained to some manifold) to study and compare the dynamics of undulatory (snake-like) bodies as they moved. Notably, DMD is a model agnostic method, meaning that no knowledge of the underlying system is necessary. This means that it can be flexibly applied to a wide range of datasets.

DSA generalizes Koopman-based comparisons into a standard set of practices:

1. Identify an optimal embedding of data (based on how well the dynamics can be represented with a linear operator).
2. Fit a linear model to the embedding to predict the dynamics.
3. Compare the learned models to compute a similarity score.

Each of these steps has a large degree of flexibility, and parts can easily be swapped in and out due to one’s goal or data constraints and assumptions. Step 1 can be modified by applying different nonlinear embeddings. Step 2 can be modified by performing different types of regression. For step 3, there exist multiple notions of similarity on linear dynamical systems, each of which can give complementary pictures. In the following section, we describe in further technical detail different variants and offer principled selection methods for each step.

### F.2 Identifying optimal DMDs

The first DSA (*Ostrow et al., 2023*) utilized the Hankel Alternative View of Koopman (*Brunton et al., 2017*), a variant of the Dynamic Mode Decomposition that fits an optimal linear dynamics operator on a delay (Hankel) embedding, where each row has structure: *H*_*t*_ = [*x*(*t*), *x*(*t* − *τ*), *x*(*t* − 2*τ*), …, *x*(*t* − *nτ*)]. However, DSA need not use delay embeddings if other nonlinear embeddings provide a better fit to the data. In general, we seek a nonlinear basis for the dynamics that maximizes the signal captured by the DMD, and to find parameters in that basis that minimize noise pollution. This implies that dimensionality reduction methods should be applied to the data to improve linear estimation and ease the search for an appropriate basis. For example, if the dynamics of a system are described by a nonlinear attractor manifold, flattening the manifold using a method such as Isomap or Locally Linear Embeddings will make DMD fitting more effective while maintaining the predominant structure of the dynamics.

In this section we review three general methods for learning DMD operators that can result in better comparisons, in order of complexity: delay embeddings, kernel methods and neural networks. In practice, we recommend beginning with delay embeddings, as we have found them to be empirically highly successful. If they fail to give interpretable results, kernel methods should be used next, and finally neural networks if neither is viable.

#### F.2.1 Delay embeddings

Time-delay embeddings construct a high dimensional nonlinear state space by stacking time-delayed copies of the state, as above. Delay embedding improves estimation capabilities over partially-observed systems (*Kamb et al., 2020*). As in the Hankel Alternative View of Koopman (HAVOK) method (*Brunton et al., 2017*), we apply the SVD and extract the top *r* ranks.

#### F.2.2 Kernel methods

Kernel methods seek to implicitly project data to a nonlinear feature space in infinite dimensions, upon which machine learning tasks such as linear classification or regression are easier to solve. Naturally, these have been applied to the setting of DMD, originally introduced as the Extended DMD (*Williams et al., 2016*). Kernel linear regression can be straightforwardly generalized from standard least squares and hence computing the DMD operator is equally as efficient, once the feature space is identified. However, kernel methods scale quadratically in the number of data points (as opposed to the number of dimensions in standard linear regression) which can make computation of the kernels highly inefficient.

One alternative is therefore to approximate kernel regression by using random features: *Rahimi and Recht, 2007* demonstrated a correspondence between different random feature spaces and well-known kernels, such as Fourier features and Radial Basis Function kernels. In fact, the limit of infinite features is not necessary to get a close approximation of the full kernel method (*Rahimi and Recht, 2007*). For the purposes of DSA, random features are effective because they can introduce useful nonlinearities as in a kernel method, but are computationally efficient. Note however that for the purposes of DSA, many random features may be highly unpredictable and hence strong noise pollution is often introduced.

#### F.2.3 Deep learning methods

When it comes to learning useful nonlinear features for the DMD, many different deep learning architectures have been explored (e.g., *Lusch et al., 2018*). Deep learning is powerful and can learn useful features for comparison. However, these methods have more hyperparameters and computational complexity than previously described methods. These challenges can make them inefficient to use in the setting of DSA.

#### F.2.4 Regression methods for the DMD operator

The DMD literature has introduced a wide variety of regression methods to improve operator fitting under different data conditions. DSA can be flexibly adapted to use any of these methods, as they all return the same object: a finite dimensional approximation of the Koopman Operator. For a full review, please see *Schmid, 2022*. The basic DMD methods apply regression as it is standardly formulated with linear regression, usually based on least-squares. Kernel DMD uses a kernelized version of the regression, utilizing the Gram matrix across samples instead of the covariance matrix across dimensions. Total-least squares (*Dawson et al., 2016*), forward-backward DMD (*Hemati et al., 2017*), BOP-DMD (*Sashidhar and Kutz, 2022*) and subspace DMD (*Takeishi et al., 2017*) all seek to improve estimation under noisy data (either observational or process noise).

When the system itself is forced by a known external input, DMD with control (DMDc) jointly estimates the autonomous and input matrices (*Proctor et al., 2016*), and its subspace variant combines noise robustness with input handling (*Huang et al., 2025*). When the system is known to have particular structure, such as invariance across reference frames (*Cohen et al., 2023*), a physics-informed DMD constrains the operator to a manifold that determines that structure (*Baddoo et al., 2023*).

### F.3 Metrics for tuning and robustness

Poor hyperparameter choice can lead to noise entering the DMD estimate, which can bias a comparison. However, there may be unavoidable noise that we would like to guard against as well. Here, we introduce two methods.

#### F.3.1 Standard prediction error metrics

Empirically, we have found that test predictivity scores of the DMD matrix are decent proxies for fitting hyperparameters. We use the normalized Akaike Information Criterion (AIC) and the Mean Absolute Standardized Error (MASE). Given a time series *y* = [*y*_0_, …, *y*_*n*_] a model with *k* parameters (here the squared rank of the DMD, *r*^2^ + 1), and a prediction series 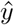,

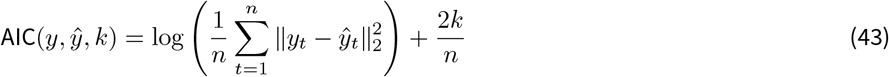

AIC is a predictivity metric that is penalized by the number of parameters *k*, which is useful for assessing overfitting.

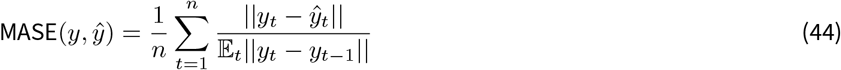

MASE is a useful predictivity metric because it is normalized by the simplest possible prediction, the persistence baseline. It is more numerically stable for MASE to be computed by normalizing each time point with the average persistence baseline prediction error, written in Eqn 44 as E_*t*_||*y*_*t*_ − *y*_*t*−1_||, rather than the persistence error at every time point.

#### F.3.2 Residual DMD

As the Dynamic Mode Decomposition (DMD) is a linear dynamics model, one can split it into eigenmodes: pairs of directions in state space with decay rates and frequencies that govern how those directions evolve under the model. However, most systems to which the DMD is fit are not linear, hence these modes are only an approximation. How good of a fit is any given mode? To answer this question, *Colbrook et al., 2023* introduced the Residual DMD (ResDMD), which computes errors (residuals) that measure how close each mode is to *actually* evolving linearly.

Given a DMD model with eigenvalues *λ*_*i*_ and eigenvectors *v*_*i*_, data matrices *X, Y* where *Y* represents the next time steps after *X*, the residuals can be defined as:

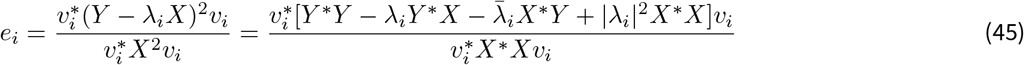

When there is a clear residual gap, signal and noise can be easily distinguished. However, when the residuals are roughly on a continuum, an optimal cutoff can be more challenging to discern. We compute the average residual, with the intuition that once noise begins polluting the DMD matrix, the average residual will start rapidly increasing. Jointly trying to minimize the average residual and prediction error is a powerful combination to maximize signal (AIC) while minimizing noise (resDMD).

### F.4 Comparison metrics

#### F.4.1 Metrics for unaligned spaces

The primary metric we utilize is the 2-Wasserstein distance over the set of (usually complex) eigenvalues *λ*^*x*^, *λ*^*y*^ for each matrix *K*_*x*_, *K*_*y*_, introduced in *Redman et al., 2024*:

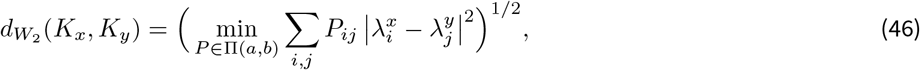

Where 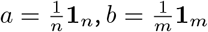, and 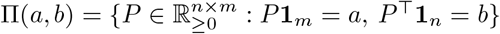 These couplings, also called transport plans, have marginals according to the original empirical distributions Λ_*x*_, Λ_*y*_. This is implemented with standard eigendecomposition and Optimal Transport algorithms via *Python Optimal Transport* (*Flamary et al., 2021*).

The metric compares the discrete part of the Koopman eigenspectrum and is most informative when the dynamics are dominated by discrete timescales and frequencies (limit cycles, quasiperiodic motion, attractors with isolated eigenvalues); strongly mixing or chaotic regimes carry a continuous spectral component that no finite eigenvalue set can fully resolve (*Brunton et al., 2022*). Second, the standard Wasserstein distance on the eigenvalues treats the spectrum as a sufficient summary of *K*, which is exact for normal operators but only approximate in general: two non-normal operators can share eigenvalues while differing in transient behavior, a discrepancy captured by the pseudospectrum (*Trefethen and Embree, 2005*). In both of these settings, we may supplement the eigenvalue comparison with Procrustes Alignment over vector fields (PAVF), which is implicitly a distance on pseudospectra as we now describe.

The Procrustes Analysis over Vector Fields (PAVF, *Ostrow et al., 2023*) is defined as a Procrustes-style optimization problem, where the optimization is over a similarity transform that describes how vector fields transform under linear coordinate transforms.

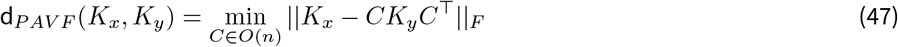

See (*Ostrow et al., 2023; Behrad et al., 2025*) for implementation details. The alignment matrix is restricted to be orthogonal because general invertible matrices make d_*P AV F*_ not a proper metric (SI F.4.2). However, this metric bears a quantitative relationship with the Wasserstein distance on eigenvalues. In particular, when *K*_*x*_ and *K*_*y*_ are normal matrices (*K*^⊤^*K* = *KK*^⊤^), d_*P AV F*_ reduces to the 2-Wasserstein distance over the eigenvalues of *K*_*x*_ and *K*_*y*_ (*Ostrow et al., 2023*).

#### F.4.2 Why minimize over orthogonal matrices?

We motivate why the original DSA metric (PAVF, *Ostrow et al., 2023*) minimizes over orthogonal matrices, rather than invertible matrices. Distance measures that are proper metrics can be studied using the standard mathematics of inner product spaces, which means that methods such as clustering and classification can be done. Minimization over invertible matrices is not a proper metric, as we prove below. Studying the eigenvalues of the matrices, however, is still a proper (pseudo)metric and also a reasonable one to utilize.

##### **Lemma 1** (Schur’s Inequality).

*For any matrix A* ∈ ℂ^*n*×*n*^ *with eigenvalues λ*_1_, …, *λ*_*n*_, *the following inequality holds:*

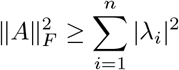

*Equality holds if and only if A is a normal matrix*.

##### Proof

The proof relies on the Schur decomposition of *A*. Any square matrix *A* can be written as *A* = *QU Q*^*H*^, where *Q* is a unitary matrix (*Q*^*H*^ *Q* = *I*) and *U* is an upper triangular matrix. The diagonal entries of *U* are the eigenvalues of *A*, so *u*_*ii*_ = *λ*_*i*_.

The Frobenius norm is invariant under unitary transformations. Therefore:

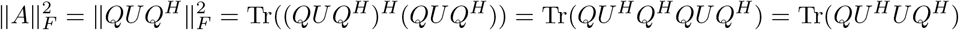

Using the cyclic property of the trace, this simplifies to:

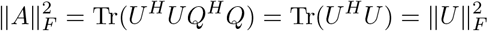

The squared Frobenius norm of the upper triangular matrix *U* is the sum of the squared magnitudes of all its entries. We can separate this sum into the diagonal and off-diagonal entries:

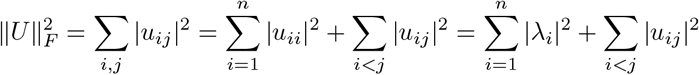

Since ∑ _*i*<*j*_ |*u*_*ij*_|^2^ ≥ 0, we have:

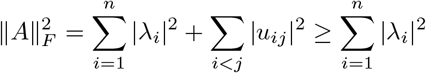

Equality holds if and only if ∑ _*i*<*j*_ |*u*_*ij*_|^2^ = 0, which means all off-diagonal entries of *U* are zero. A matrix *A* is unitarily diagonalizable (i.e., its Schur form *U* is diagonal) if and only if it is normal (*A*^H^ *A* = *AA*^H^).

##### Proposition F.1.

*The function D* : ℝ^*n*×*n*^ × ℝ^*n*×*n*^ → ℝ_*≥*0_ *defined by*

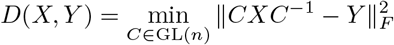

*is not a metric*.

##### Proof

We test the symmetry axiom of a metric, *D*(*X, Y*) = *D*(*Y, X*), by finding a counterexample. We begin by expanding the objective function using the trace definition of the Frobenius norm, 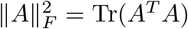.

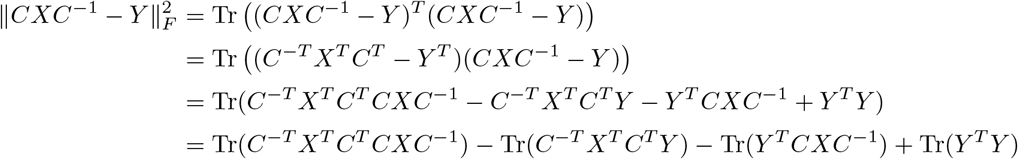

Using the cyclic property of the trace, Tr(*ABC*) = Tr(*CAB*), we note that Tr(*C*^−*T*^ *X*^*T*^ *C*^*T*^ *Y*) = Tr(*Y*^*T*^ *CXC*^−1^). The expression simplifies to:

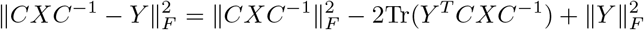

Thus, 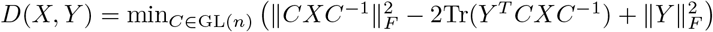.

Now, assume for contradiction that *D*(*X, Y*) = *D*(*Y, X*) for all *X, Y*. Let us choose a specific counterexample. Let *X* = *I* (the identity matrix) and let *Y* be any matrix that is diagonalizable but **not** normal.

First, we compute *D*(*I, Y*):

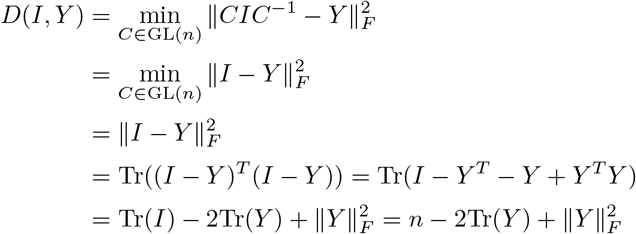

Next, we compute *D*(*Y, I*):

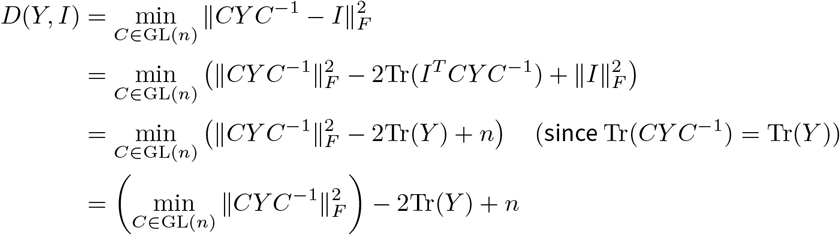

The last step follows because the minimization only affects the first term.

By our symmetry assumption, *D*(*I, Y*) = *D*(*Y, I*), so:

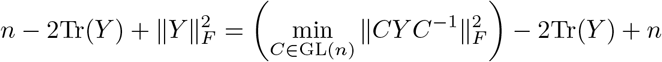

Canceling the constant terms yields:

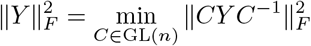

However, by Lemma 1 (Schur’s inequality), we know 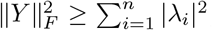, with equality if and only if *Y* is normal. For a diagonalizable matrix, it is known that 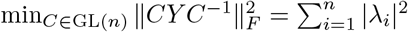. Since we chose *Y* to be not normal, the inequality is strict:

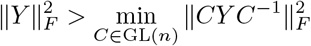

This contradicts the equality derived from the symmetry assumption. Therefore, the symmetry axiom does not hold, and *D*(*X, Y*) is not a metric.

#### F.4.3 Metrics for aligned spaces

The metrics defined above are notably designed to handle misaligned spaces. This is important when the observed dimensions of a system are randomly chosen, as in the experimental recording of neurons, or the random selection of participants in an economic survey. However, there are cases in which dynamical comparison is done in *aligned spaces*. There, the complex alignment process of the above metrics could be dropped. However, note that these alignment procedures are also designed to remove geometric deformations, such as stretching and rotations in the Koopman lifted space. Thus, when dynamical comparison is done in aligned spaces, the combination of aligned and unaligned metrics is additionally informative.

In the case of aligned spaces, we also introduce here a notion of *subspace angle*, which was used as a similarity metric in *Cohen et al., 2023*. Given two dynamics matrices *K*_*x*_, *K*_y_, an orthonormal basis for each (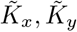) is computed (for example, via SVD or QR decomposition). Then, the subspace angles are computed as:

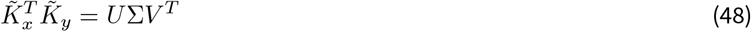

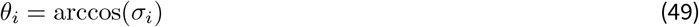

Where *σ*_*i*_ is the *i*-th singular value defined by Σ. *θ*_*i*_ is defined as the *i*-th principal angle, and the Grassmannian distance (sub-space distance) is defined as:

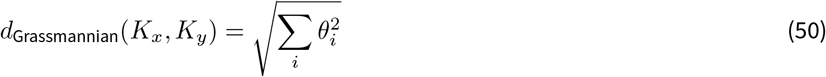

This metric captures the *orientation* of the operators relative to one another. This is informative when the original spaces are equivalent and the operators are not full rank. This metric captures the similarity of eigenmodes that describe the evolution of each system, but not their temporal properties (which are captured by the eigenvalues) or the relationships between eigen-modes *within the same system*. In a related fashion, (*Germain et al., 2026*) combined the Grassmannian distance on eigenspaces with the Wasserstein distance on eigenvalues to jointly compare eigenvalues and eigenmodes of an operator.

#### F.4.4 Comparison of input-driven dynamics

When dynamics are forced by external input, it is a natural question to ask how similar two dynamical systems are in their response to input. InputDSA (*Huang et al., 2025*) introduced a metric to approach this question based on the dynamical systems notion of *controllability*, which is tractably defined for input-driven linear systems. Given two linear dynamical systems:

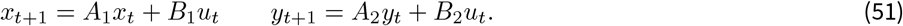

Controllability is defined via the T-step controllability matrix (with T typically taken as the dimension of the system):

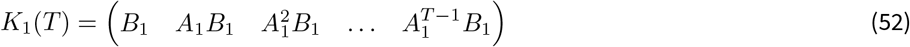

and its corresponding Gramian:

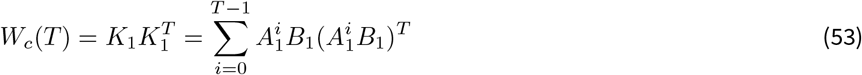

Controllability, as measured by the eigenvalues of the Gramian, is only preserved under orthogonal transformations between state spaces:

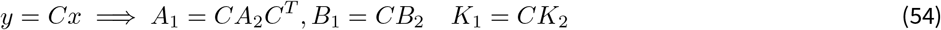

This motivates the InputDSA metric:

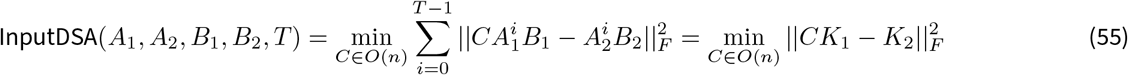

which is efficiently solved with Procrustes Analysis. See *Huang et al., 2025* for further details.

#### F.4.5 Picking PAVF or Wasserstein

The observation that PAVF and Wasserstein are equivalent under normality suggests that PAVF is more restrictive and is sensitive to differences in *non-normality* as well. We make the connection formal here, by connecting to the pseudospectrum of an operator (*Trefethen and Embree, 2005*): for some *ϵ* > 0 and matrix *A*, the *ϵ*-pseudospectra is defined by the set *σ*_ϵ_(*A*) = {*z* ∈ ℂ : ∥(*zI* − *A*)^−1^∥_2_ > *ϵ*^−1^}. Intuitively, this describes how the eigenvalues of a matrix change under perturbation, and it can be used to describe the transient behavior of a linear dynamical system. The pseudospectra are naturally connected to classical eigenspectra by taking the limit *ϵ* → 0. Two linear dynamical systems can have equivalent eigenvalues but different transient behavior. The following theorem makes the connection to PAVF explicit:

##### **Theorem F.2** (Orthogonal Transformations Preserve Transient Behavior).

*If S* ∈ ℝ ^*N*×*N*^ *is orthogonal*,

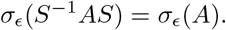

##### Proof

Let ∥*A*∥_2_ = *σ*_max_(*A*). Choose *z* ∈ *σ*_ϵ_(*A*). Then

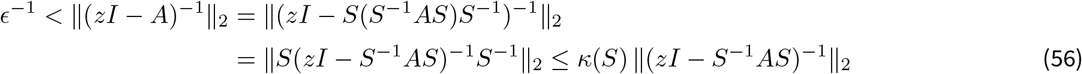

where *κ*(*S*) = ∥*S*∥_2_∥*S*^−1^∥_2_ = *σ*_max_(*S*)*/σ*_min_(*S*). This implies that *z* ∈ *σ*_*ϵκ*(*S*)_(*S*^−1^*AS*), or *σ*_ϵ_(*A*) ⊆ *σ*_*ϵκ*(*S*)_(*S*^−1^*AS*).

Next, suppose *z* ∈ *σ*_*ϵ*/*κ*(*S*)_(*S*^−1^*AS*). Then

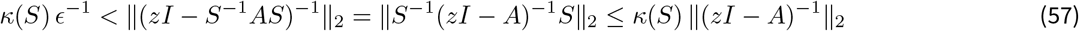

which implies *σ*_*ϵ*/*κ*(*S*)_(*S*^−1^*AS*) ⊆ *σ*_ϵ_(*A*). Putting the two inclusions together,

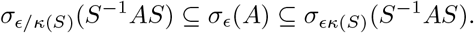

Since *S* is orthogonal, *κ*(*S*) = 1, and the two pseudospectra coincide.

Therefore, for the PAVF metric to be zero, not only does the distance between each operator’s eigenvalues have to be zero, but also their *pseudospectra*. This therefore motivates a method for choosing when to use PAVF. Two standard measures of non-normality are (*Trefethen and Embree, 2005*):

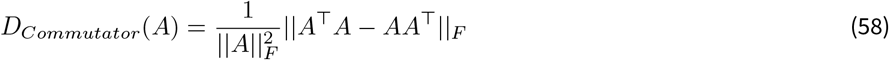

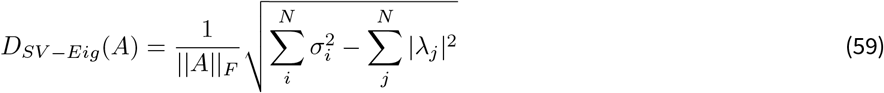

Where *σ*_*i*_, *λ*_*j*_ are the i-th singular values and j-th eigenvalues, respectively, and the prefactor divisor is a scaling term to enable comparison across dimensionalities. When these measures are large, transient behavior (non-normal) differs substantially from the asymptotic behavior (normal), which suggests that comparison of pseudospectra is additionally informative.

Next, we demonstrate these ideas on linear systems.

#### f.4.6 Toy example: normal and non-normal linear systems

We borrow a classical example from (*Trefethen and Embree, 2005*) to demonstrate the distinction. We define three linear operators:

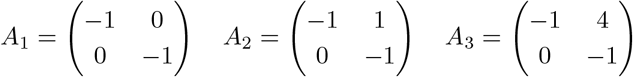

and visualize their dynamics under 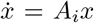 in Fig. S3a. Under *A*_2_, the dynamics decay, yet more slowly than *A*_1_. Under *A*_3_, the state transiently grows before decaying to zero. Its pseudospectrum can be observed to be very wide relative to the other matrices (Fig. S3b right). This is reflected in each non-normality metric.

We compute the PAVF and 2-Wasserstein distance for these matrices, noting that the PAVF metric identifies the dissimilar transient behavior, whereas the 2-Wasserstein distance identifies the similar asymptotic behavior. Likewise, there is a strict increase in the PAVF score as a function of increasing non-normality, when comparing to the baseline *A*_1_ (Fig. S3c). When studying the behavior of the 2 × 2 matrix *A*_*i*_ for a range of values on the upper triangular entry *a*_01_, we note that transient growth only occurs when *ω*(*A*_*i*_) ≥ 0 (Fig. S3d). The changes in non-normality can effectively be captured by the two non-normality scores, although note that these say nothing about matrix similarity (Fig. S3e). Lastly, we increased the value of the upper triangular parameter *a*_10_ from 0 to 10 and compared with PAVF to the baseline matrix *A*_1_ (Fig. S3f). Here we note a strict monotonic relationship between the non-normality score and the PAVF distance.

**Figure S1.**
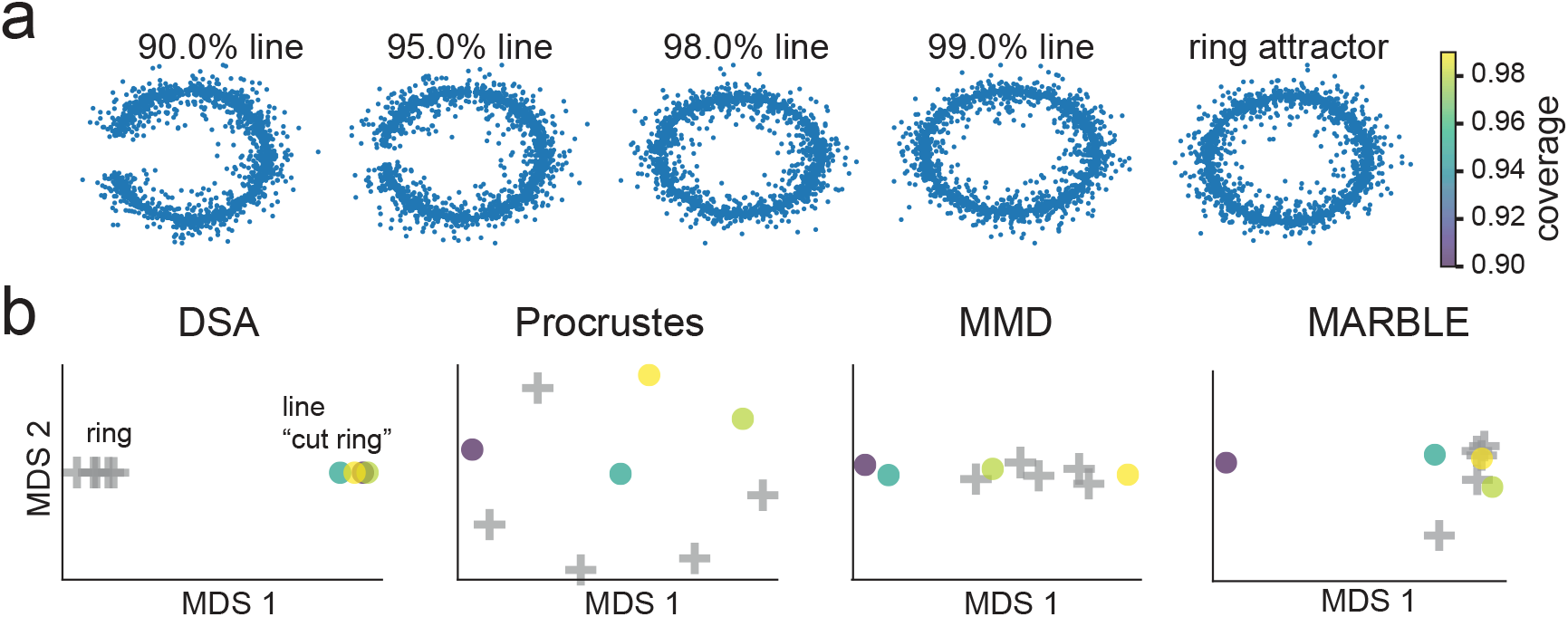
DSA distinguishes ring attractors from geometrically near-ring line attractors, while baseline methods do not. To test whether geometric similarity confounds dynamical comparison, we constructed a family of cut ring (line) attractor models whose attractor manifolds progressively approach a full ring (extending the arc until it closes), and compared each to a true ring attractor. **(a)** State-space scatterplots of trajectories from each model. Percentages denote the fraction of the full ring covered by the attractor; the rightmost panel shows a standard ring attractor for reference. As the percentage approaches 100%, the cut ring attractor becomes geometrically indistinguishable from the ring, despite its dynamics remaining qualitatively different from the ring’s continuous attractor. **(b)** Multidimensional scaling (MDS) embedding of the pairwise DSA distance matrix between all models in (a). DSA cleanly separates the ring from every line attractor regardless of geometric overlap, indicating that DSA captures the underlying dynamical (rather than geometric) distinction. Procrustes on trajectory geometry, MMD, and MARBLE do not.

**Table S1.**
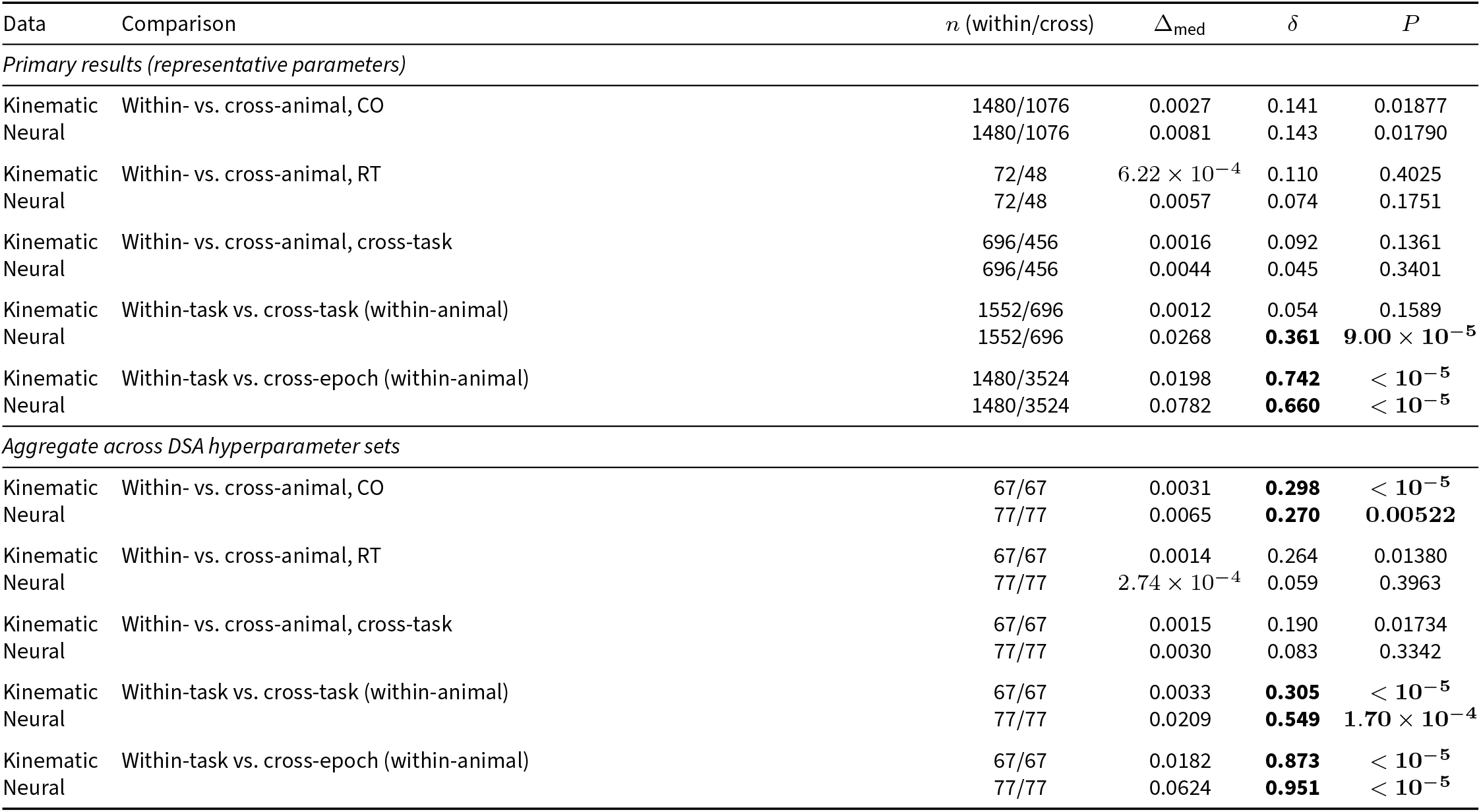
Session-label permutation tests for DSA distance comparisons. P-values are from one-sided session-label permutation tests (*n*_perm_ = 100,000). *P* values are uncorrected; significance is assessed against a Bonferroni-adjusted threshold α = 0.05/8 = 6.25 *×* 10^−3^ for *m* = 8 simultaneous comparisons. Bold indicates *P* < *α*. Δ_med_ is the median difference between the two distributions being compared. δ is Cliff’s delta (Methods D.4.3.

| Data | Comparison | $n$ (within/cross) | $\Delta_{\text{med}}$ | $\delta$ | $P$ |
| --- | --- | --- | --- | --- | --- |
| *Primary results (representative parameters)* |  |  |  |  |  |
| Kinematic | Within- vs. cross-animal, CO | 1480/1076 | 0.0027 | 0.141 | 0.01877 |
| Neural |  | 1480/1076 | 0.0081 | 0.143 | 0.01790 |
| Kinematic | Within- vs. cross-animal, RT | 72/48 | $6.22 \times 10^{-4}$ | 0.110 | 0.4025 |
| Neural |  | 72/48 | 0.0057 | 0.074 | 0.1751 |
| Kinematic | Within- vs. cross-animal, cross-task | 696/456 | 0.0016 | 0.092 | 0.1361 |
| Neural |  | 696/456 | 0.0044 | 0.045 | 0.3401 |
| Kinematic | Within-task vs. cross-task (within-animal) | 1552/696 | 0.0012 | 0.054 | 0.1589 |
| Neural | | 1552/696 | 0.0268 | **0.361** | $9.00 \times 10^{-5}$ |
| Kinematic | Within-task vs. cross-epoch (within-animal) | 1480/3524 | 0.0198 | **0.742** | $< 10^{-5}$ |
| Neural | | 1480/3524 | 0.0782 | **0.660** | $< 10^{-5}$ |
| *Aggregate across DSA hyperparameter sets* |  |  |  |  |  |
| Kinematic | Within- vs. cross-animal, CO | 67/67 | 0.0031 | **0.298** | $< 10^{-5}$ |
| Neural |  | 77/77 | 0.0065 | **0.270** | **0.00522** |
| Kinematic | Within- vs. cross-animal, RT | 67/67 | 0.0014 | 0.264 | 0.01380 |
| Neural | | 77/77 | $2.74 \times 10^{-4}$ | 0.059 | 0.3963 |
| Kinematic | Within- vs. cross-animal, cross-task | 67/67 | 0.0015 | 0.190 | 0.01734 |
| Neural |  | 77/77 | 0.0030 | 0.083 | 0.3342 |
| Kinematic | Within-task vs. cross-task (within-animal) | 67/67 | 0.0033 | **0.305** | $< 10^{-5}$ |
| Neural | | 77/77 | 0.0209 | **0.549** | $1.70 \times 10^{-4}$ |
| Kinematic | Within-task vs. cross-epoch (within-animal) | 67/67 | 0.0182 | **0.873** | $< 10^{-5}$ |
| Neural | | 77/77 | 0.0624 | **0.951** | $< 10^{-5}$ |

**Figure S2.**
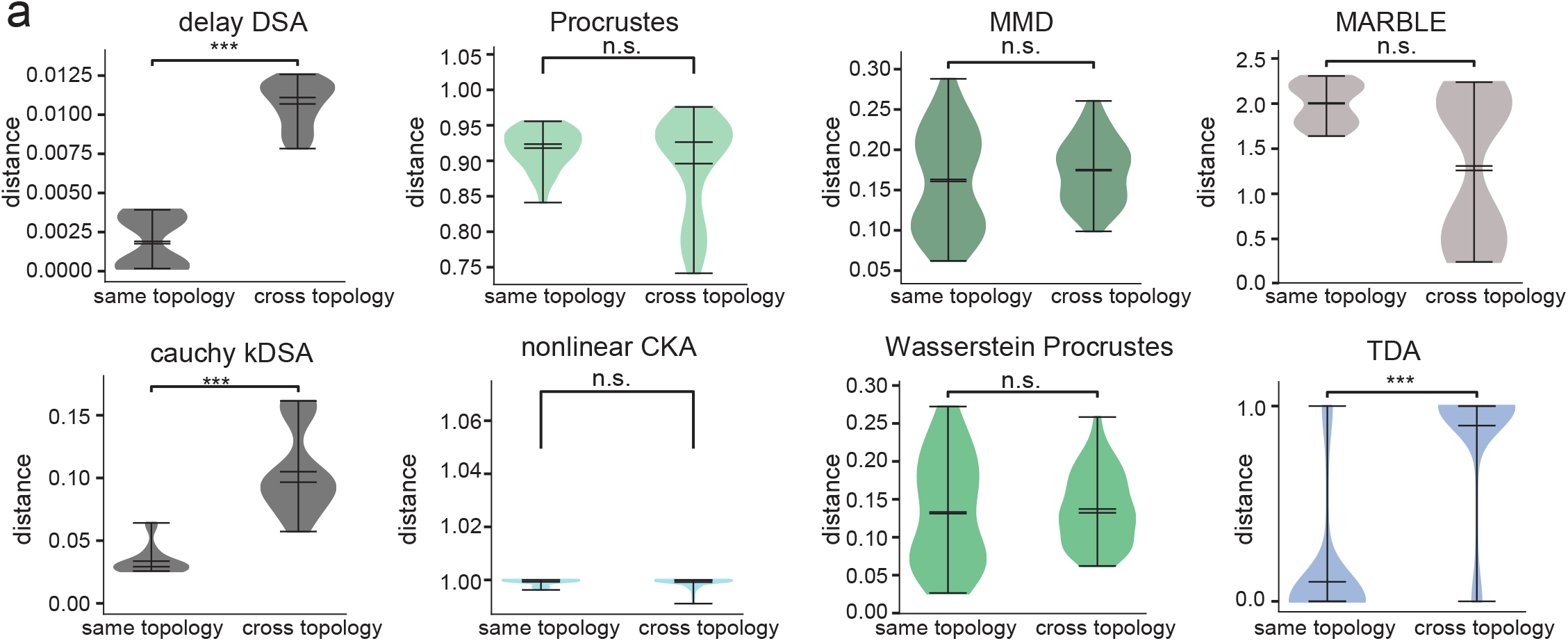
Extended cross-method analysis. **(a)** DSA distances between rings and cut rings (cross-topology) are significantly greater than distances between models of the same type, regardless of their label class. Other methods besides TDA cannot do so (but TDA cannot distinguish rings and limit cycles).

**Figure S3.**
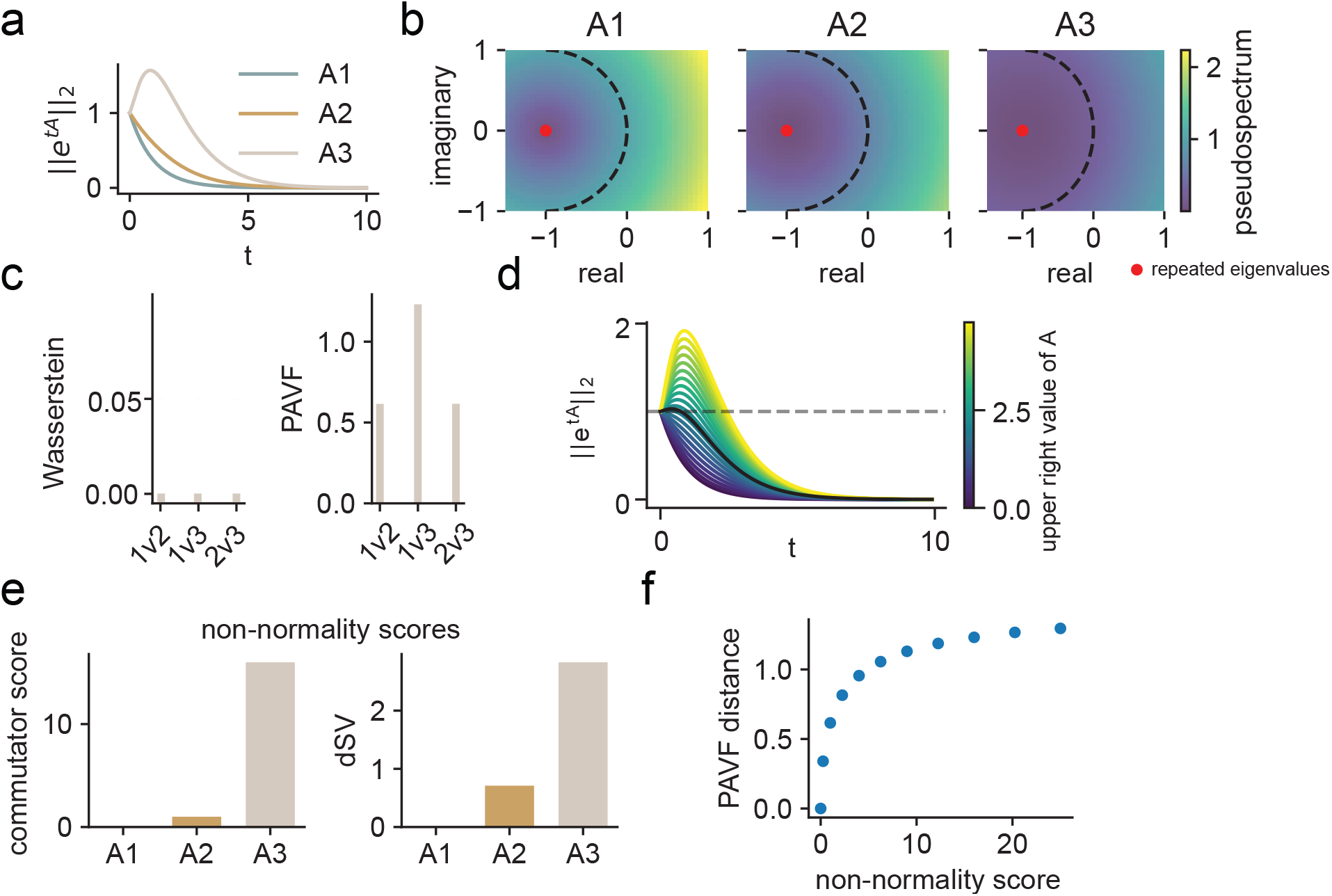
Non-normality scores can be used to determine which DSA metric to use. **(a)** Visualization of the norm of the activity vector under three linear systems, each progressively less normal than the last. **(b)** Visualization of the pseudospectrum and eigenvalues of each matrix, with a circle of radius one superimposed. The more non-normal matrix is much more easily made unstable, as demonstrated by the smaller value of the pseudospectrum (color) required to breach the instability threshold (unit circle). **(c)** The Wasserstein metric reports that the three systems are equivalent whereas the Procrustes Analysis over Vector Fields (PAVF) metrics identifies a difference. **(d)** The numerical abscissa captures when transient growth emerges across a range of linear dynamics. a_01_ represents the value of the upper triangular parameter in the 2 *×* 2 matrix with − 1 on the diagonal. **(e)** Two non-normality scores can be used to determine that A2 and A3 are much more non-normal than A1 **(f)**. The relationship between the commutator score and the PAVF distance between each of the matrices in (d) and the purely diagonal matrix A_1_.

**Figure S4.**
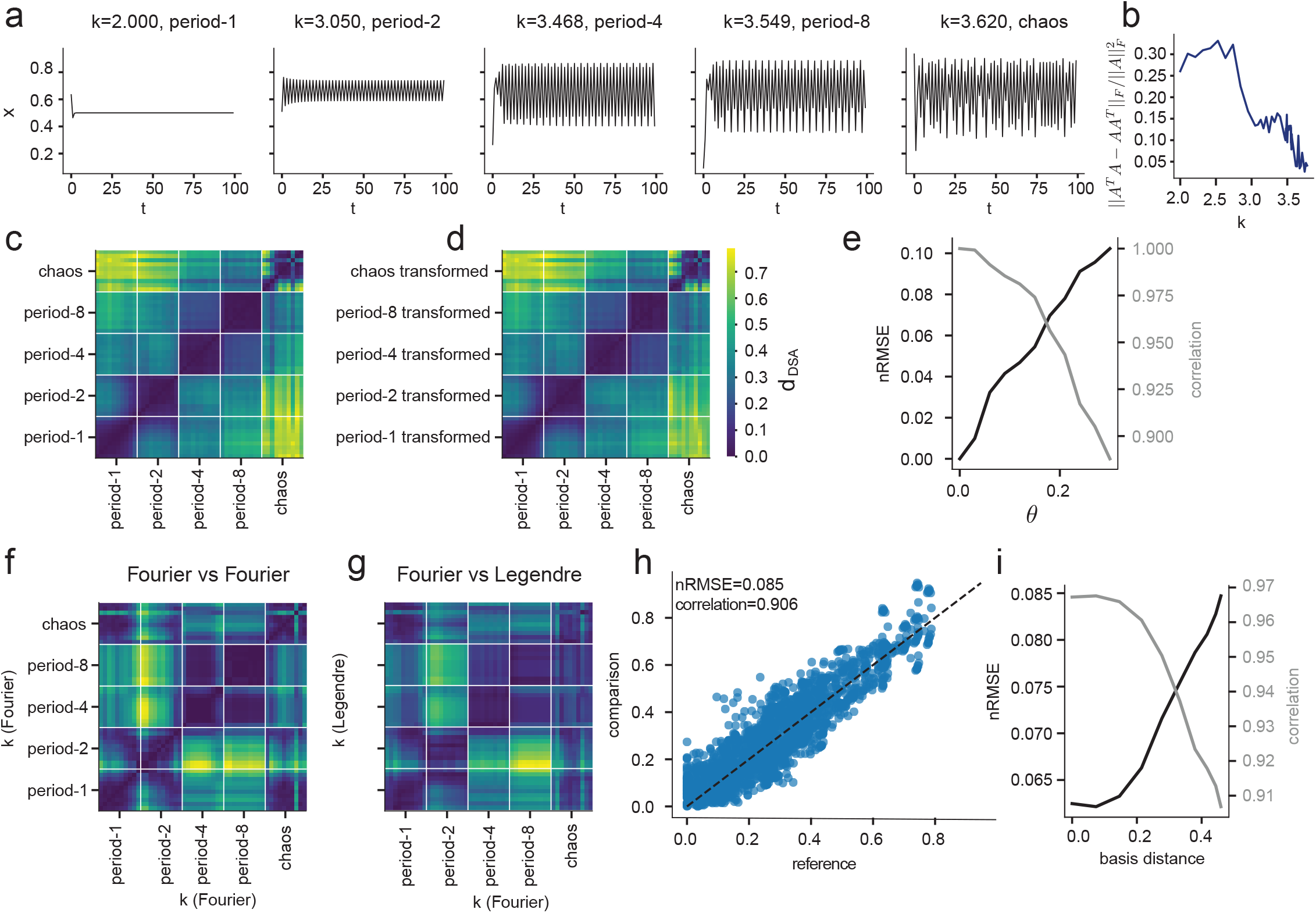
DSA captures transition to chaos in the logistic map and is robust to different observables. We apply DSA to a setting described in (*Godara et al., 2026*). For further technical details, see their paper. Data is sampled from the logistic map *x*_*t*+1_ = *kx*_*t*_(1 − *x*_*t*_) *x ∈* [0, 1], *k* ∈ [2, 4] for 50 randomly-initialized trials of 100 timesteps each. **(a)** Sample trajectories of one system in each bifurcation regime. 50 different selections of *k* are chosen, with 10 in each regime. **(b)** Non-normality metric for DMDs applied to each value of *k*. Matrices are relatively normal, as trajectories are primarily on the attractor. This motivates the use of the 2-Wasserstein distance on DMD eigenvalues. **(c)** DSA distance between data sampled from logistic map with varying *k* parameter. Bifurcation regime boundaries are denoted in white. **(d)** Data is transformed with a smooth conjugacy map *h*_*θ*_(*x*) = *x* + *θ* sin(π*x*) with θ = 0.1, then compared back to the original DSA matrix in (c). **(e)** nRMSE and Pearson correlation between transformed distance matrices as in (d) and original matrix (c) across different parameters of the conjugacy map. The normalized root mean squared error (nRMSE) is computed by measuring the elementwise linear regression error from each distance in (d) to its corresponding distance in (c), and the Pearson correlation relates the transformed matrix to the original matrix. DSA is highly robust to these transformations. **(f)** Approximate kernel DSA distance using a Fourier basis of 25 features. **(g)** DSA distance between DMD models fit to a Fourier feature basis on the logistic map data or a Legendre feature basis of the same size. Each of these kernels is universal, meaning that they span the ℒ^2^ function space, so it is expected that a DMD model fit well should have similar features to each basis. **(h)** Elementwise scatterplot between each distance value in (f) and (g). Matrices are highly related in both nRMSE and correlation despite using different features. **(i)** nRMSE and correlation between matrices as in (g) and as the comparison feature space (Legendre in g) is interpolated between the Fourier and Legendre feature spaces. See (*Godara et al., 2026*) for further details. Here once again, DSA is robust to changes in kernel features.

**Figure S5.**
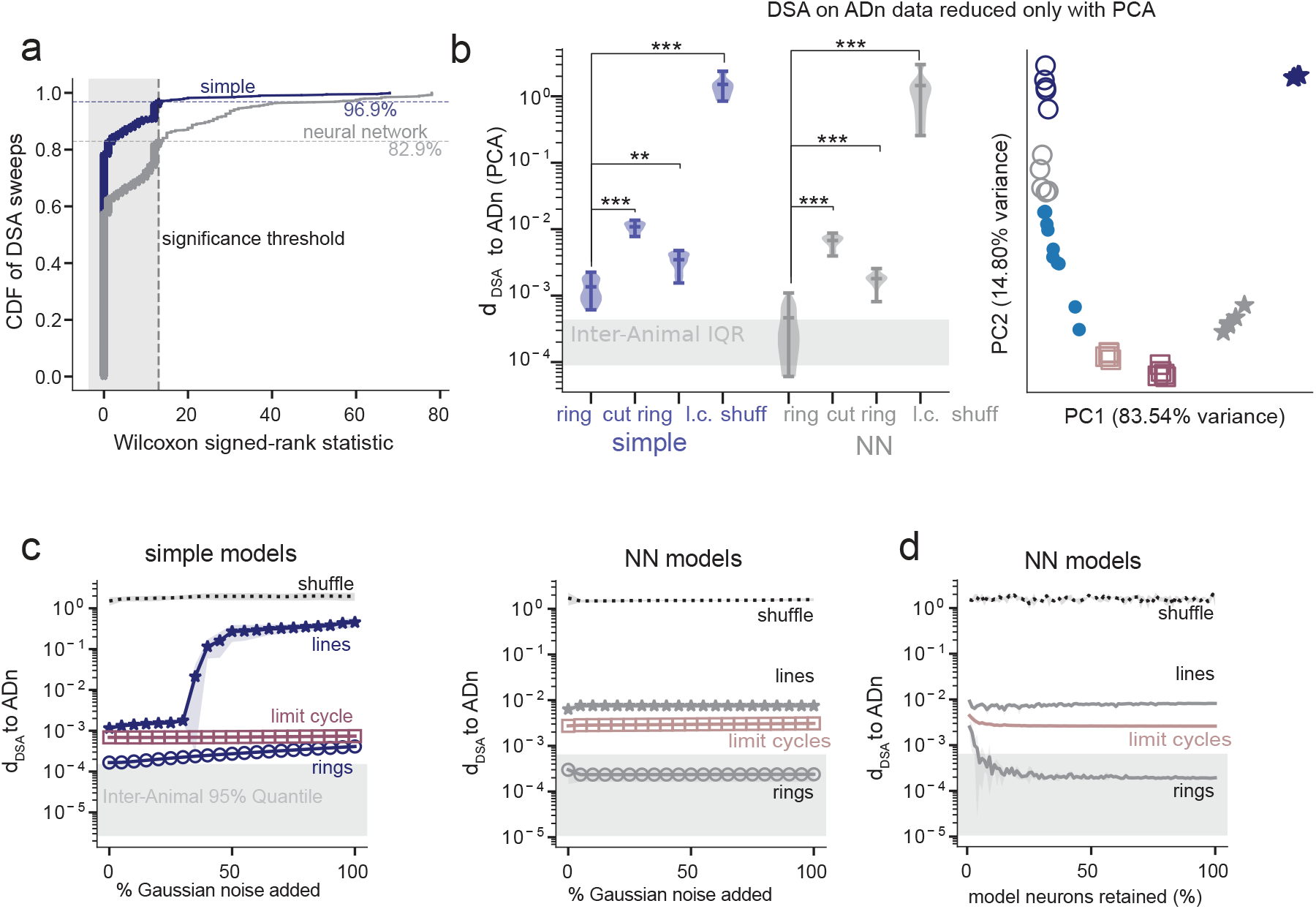
DSA robustly clusters the wake ADn data with the ring models across hyperparameters. **(a)** Cumulative distribution (CDF) of the Wilcoxon Signed-Rank Statistic testing the hypothesis DSA(ring, ADn) < DSA(cut ring,ADn). CDF is computed over a wide range of DSA hyperparameters. A statistic of 0 indicates that all ring distances are strictly less than the line distances. Over 912 hyperparameter combinations, we found that the simple ring test statistic was significant 96.9% of the time, and the NN ring was significant 82.9% of the time. Almost all of those p-values were associated with a Wilcoxon statistic of 0, which indicates no overlap between the distributions (all ring distances are strictly less than the line distances). **(b)** Replication of DSA analysis on ADn data preprocessed only with PCA. **(c)** Each model class is compared to data as Gaussian noise is added to each data dimension. Variance is added from 0 to 100% of the data variance. Left displays the simple models, right displays the NN models. NN models are more robust, and orderings are preserved across all amounts of noise added. For the NN models, 10% of neurons are observed across all noise conditions. Curves are averaged over 3 independent runs of noise. **(d)** The same analysis in (c) is performed for the NN rings (with fixed 5% Gaussian noise added) as the fraction of neurons retained before the analysis is swept from 1 to 100%. Here too, the models are robust for even 1% of neurons retained in the analysis.

**Figure S6.**
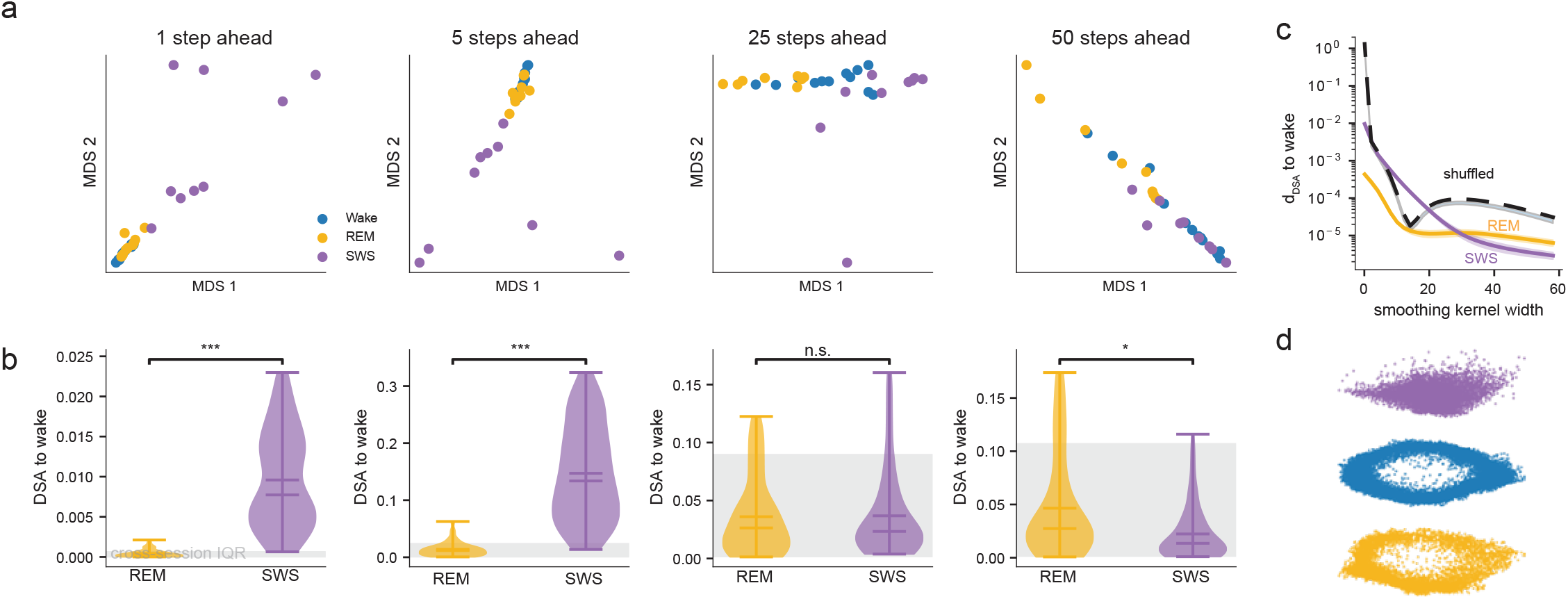
DSA differentially clusters wake, REM and SWS data across different timescales. **(a)** MDS clustering analysis for DMD models fit with various steps ahead: *X*(*t* + *s*) = *K*_*s*_*X*(*t*) for *s ∈* {1, 5, 25, 50}. **(b)** Distribution of distances to waking data for REM and SWS data for each of the steps ahead values in (a). **(c)** DSA distance to waking data as a function of increasing the Gaussian smoothing kernel width, as in Fig. 3h using kernel width instead of steps ahead. **(d)** Visualization of a single neural dataset (mouse 12 day 6 dataset), top 2 Isomap dimensions in SWS (purple), wake (blue) and REM (yellow).

**Figure S7.**
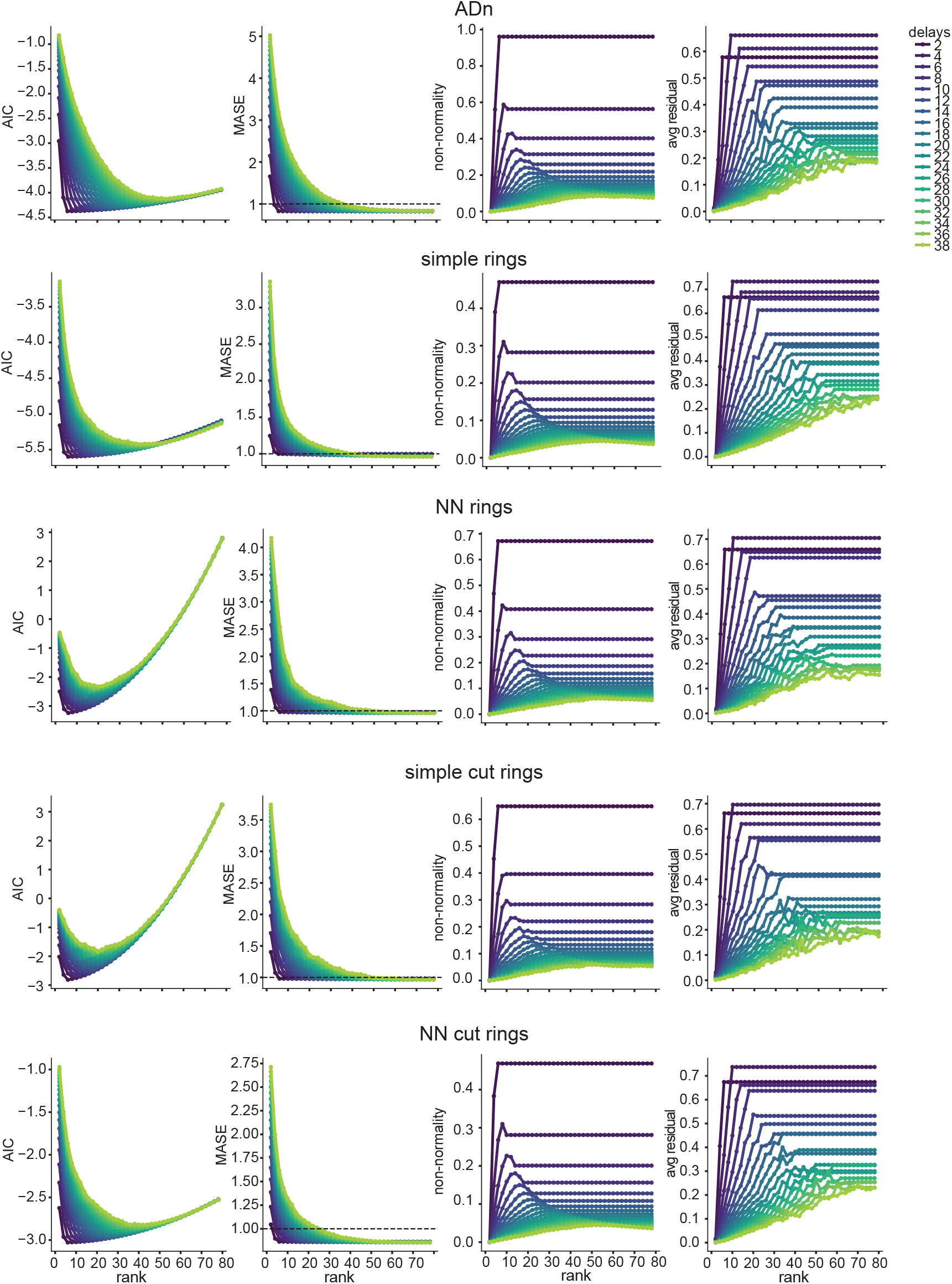
DSA hyperparameter tuning curves for ADn data and each type of ring dataset. Far left, Akaike Information Criterion (AIC). Middle left, Mean Absolute Standardized Error (MASE). Middle right, non-normality score. Far right, average ResDMD residual for all eigenvalues. x-axis indicates the rank of the DMD model, y-axis indicates the number of delays in the delay embedding.

**Figure S8.**
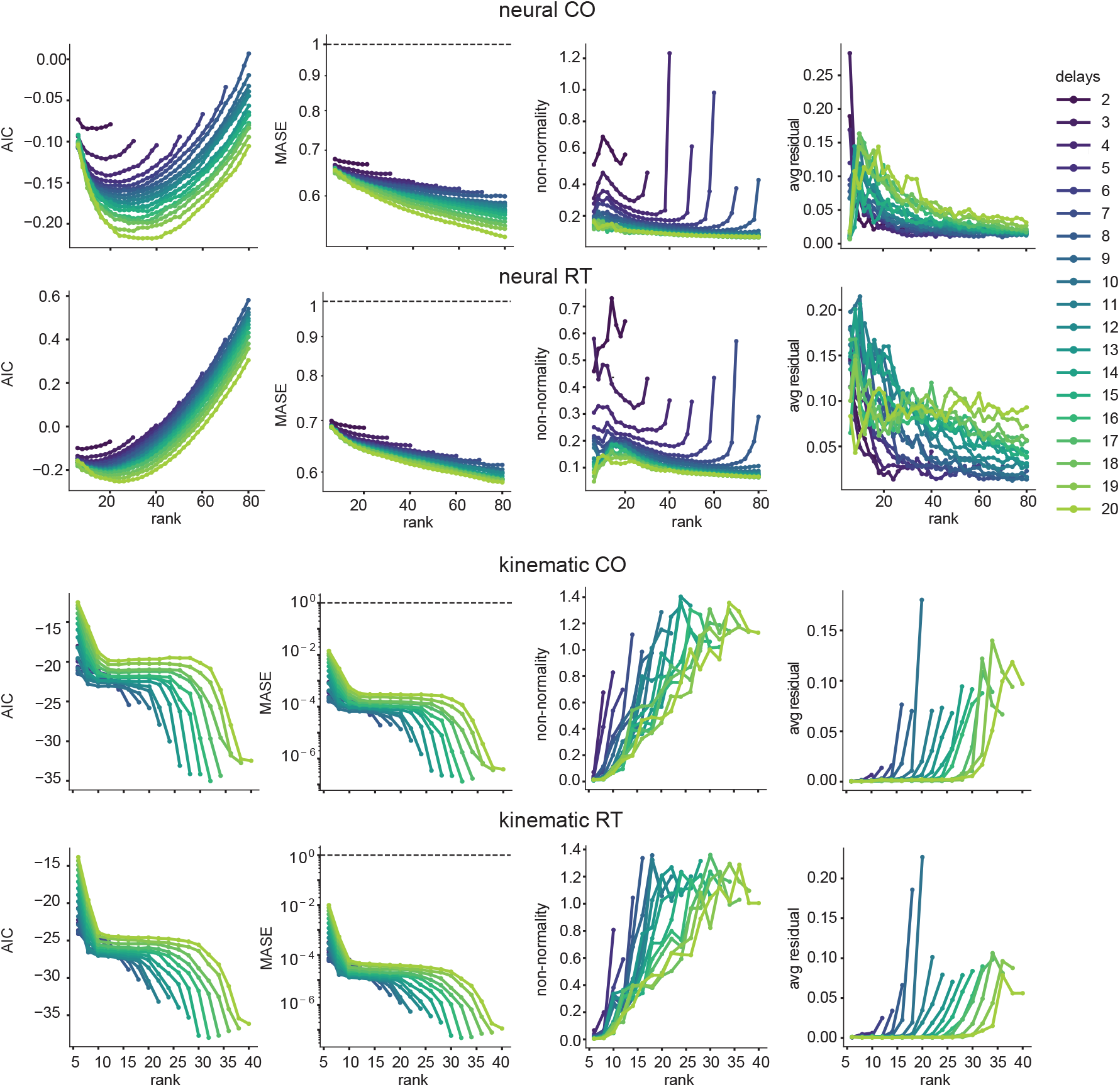
Hyperparameter tuning curves for each type of dataset and task type, as in Fig. S7.

**Figure S9.**
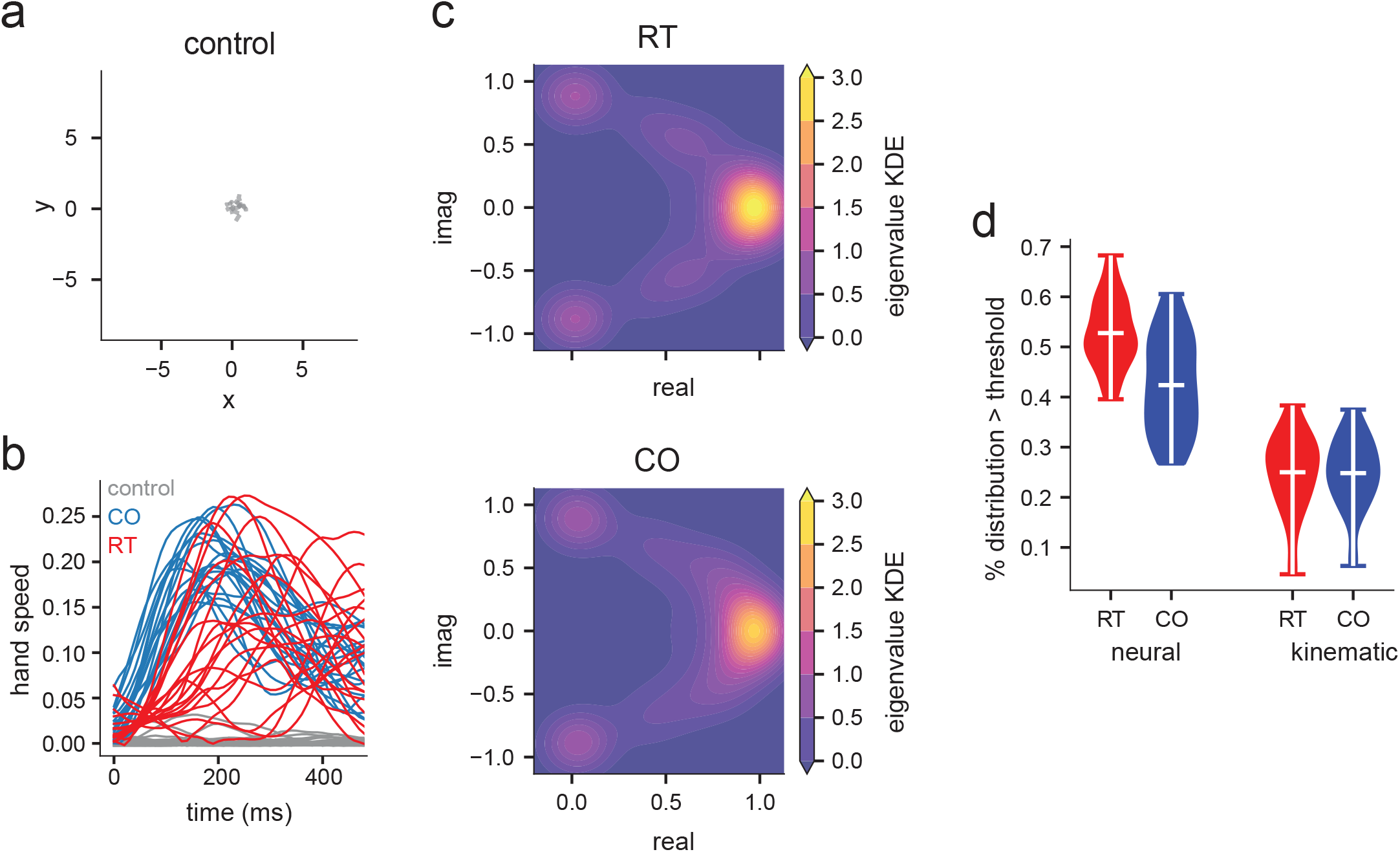
Reach Kinematic Extended Analysis. **(a)** Control (Preparatory epoch) trajectories for individual trial conditions as in Fig. 4b. **(b)** Sample trajectories of hand speed over time for individual trials. **(c)** Kernel Density Estimate of kinematic eigenvalues across all sessions and animals for the RT task (top) and CO task (bottom). **(d)** Cumulative density of the eigenvalue distribution differences that are above the null threshold for neural and kinematic datasets, comparing CO and RT dynamics. Eigenvalues for all sessions are aggregated and a Gaussian KDE is computed with bandwidth equal to the variance of the data, then the difference between CO and RT KDEs are computed on a 300 *×* 300 grid in the complex square on [− 1, 1]. A null distribution is computed by randomly shuffling CO and RT labels, then recomputing the KDE and taking the difference between the CO and RT shuffled sessions. The 95% CI is computed and the percent significant is defined as the number of grid points where the test KDE value is outside the CI. Left, neural dataset. Right, kinematic dataset.

**Figure S10.**
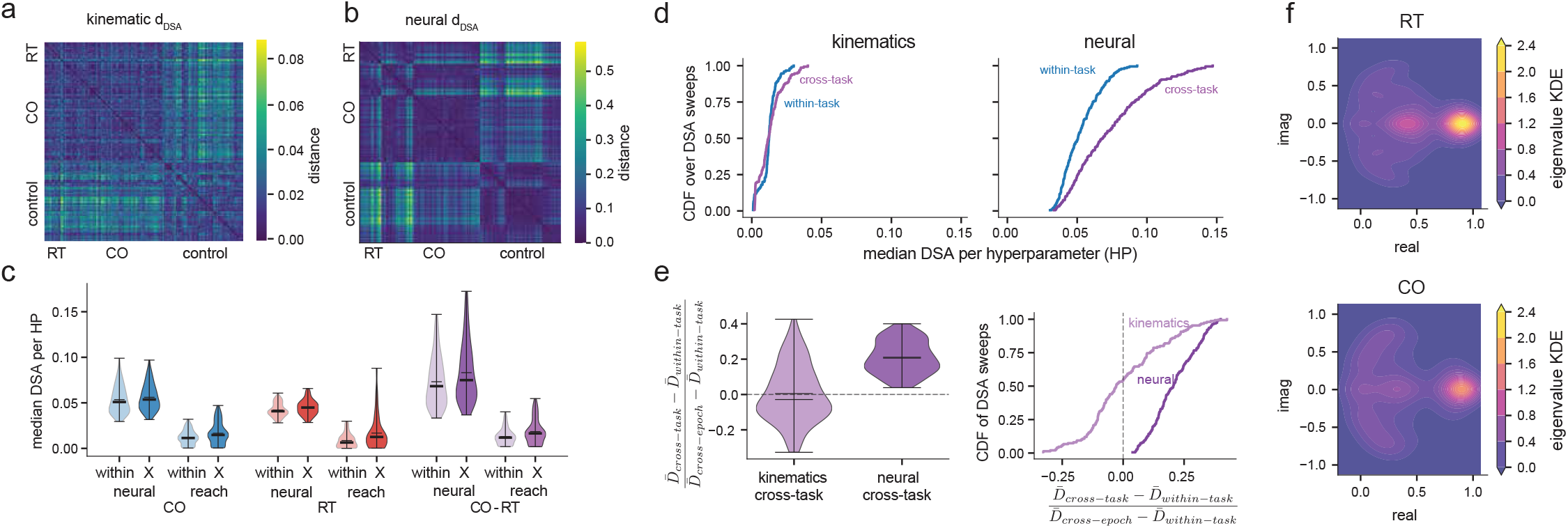
Reach results are robust across DSA hyperparameters. **(a)** DSA matrix on kinematic data evaluated in Fig. 4. There is no visual distinction in the distances between RT and CO datasets. **(b)** DSA matrix on neural data evaluated in Fig. 4. There is a clearer distinction between RT and CO than in the kinematic data. **(c)** Distribution of median within and cross (X) animal distances for each type of comparison, over all DSA hyperparameters (HP) **(d)** CDF of DSA distances across DSA hyperparameters for cross-task, within-animal versus within-task, within-animal comparisons for kinematic data (left) and neural data (right). **(e)** Left, distribution of normalized distances over DSA hyperparameters. Denoting 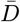 as a median over the distances for a single DSA matrix, and using the within-task, within-animal distribution as the noise floor, and the cross-epoch, within-animal distribution as the noise ceiling, the normalized distance is defined as 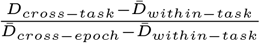 for cross-task, within-animal comparisons. Right, cumulative distribution function (CDF). **(f)** KDE density of neural eDMD eigenvalues across all sessions and animals for the RT task (top) and CO task (bottom).

**Figure S11.**
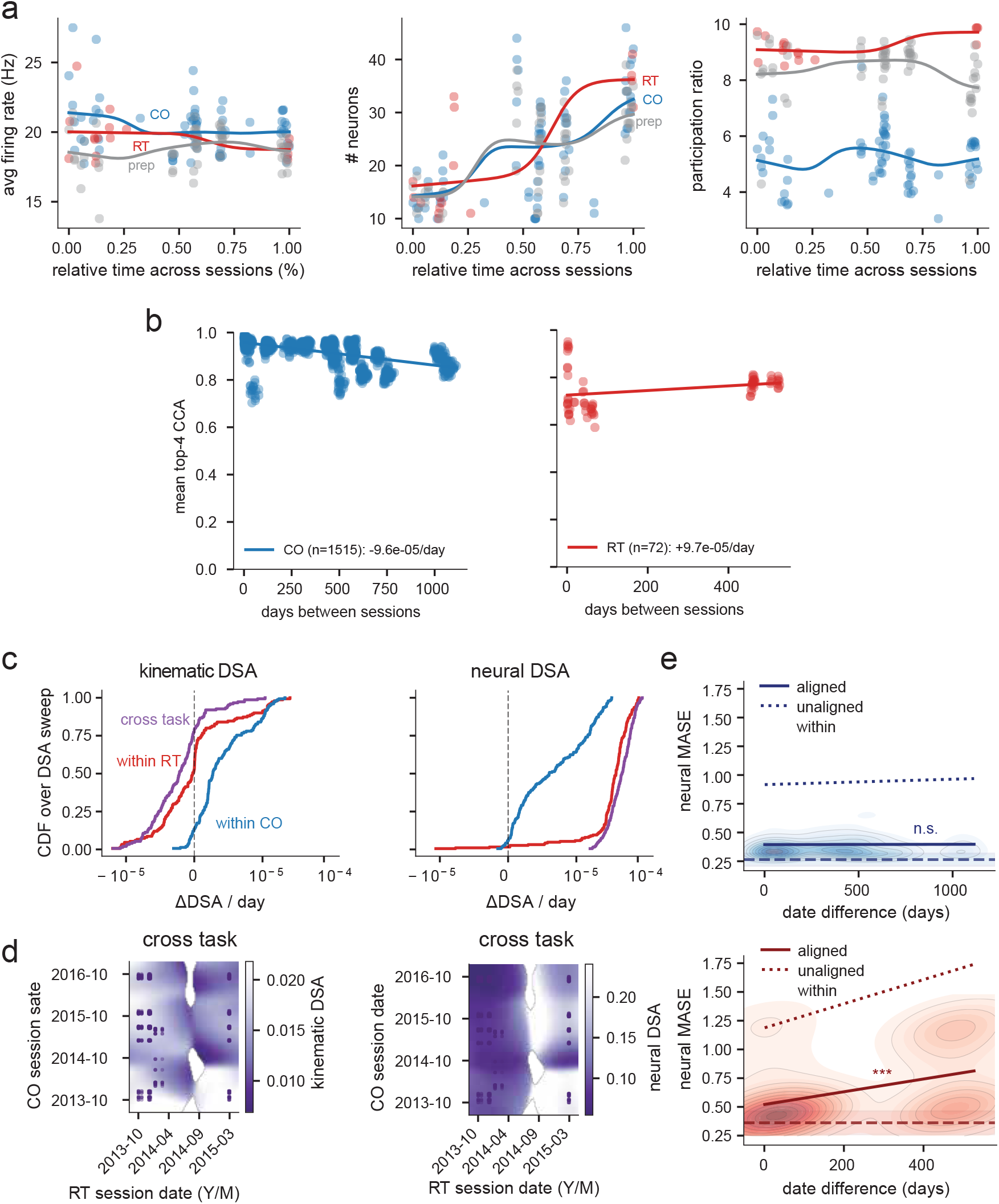
Longitudinal DSA analyses are robust across hyperparameters and provide a clear picture about the divergence between CO and RT across time. **(a)** Average firing rate, number of neurons recorded, and participation ratio (PCA dimensionality) across sessions for CO, RT and preparatory control epoch. Solid line denotes a KDE interpolation. **(b)** Mean of the top 4 Canonical Correlations applied to trajectory data (exactly the same data on which DSA is applied) compared as a function of time between sessions for CO (left) and RT (right). Solid line is the line of best fit. **(c)** CDFs of the longitudinal DSA slopes across DSA hyperparameters for kinematic distances (left) and neural distances (right), separated by comparison type. **(d)** DSA between tasks labelled by session times, aggregated across animals, using the hyperparameters in Fig. 4j. For the neural cross task distance (right), early RT sessions have small distances to all CO sessions, while later RT sessions diverge. **(e)** We evaluated the cross-session prediction error before and after alignment with the PAVF metric (Table S5). The learned PAVF alignment matrix is numerically well-conditioned, motivating its use over the typically ill-conditioned alignment matrix learned by the Wasserstein metric. In CO, prediction error did not change significantly with session date difference, with or without PAVF alignment, and the aligned prediction was nearly as good as the within-session test error. In RT, both unaligned and aligned prediction errors grew significantly with session date difference, reaching the level of a null persistence baseline after ~ 200 days. DMD models therefore successfully transfer across CO sessions over the entire recording window, but not RT.

**Table S2.** Robustness of permutation tests across DSA hyperparameters. Fraction of HP settings where the per-cell permutation test is significant at *α* = 0.05/8 = 6.25 *×* 10^−3^ (Bonferroni-corrected), with uncorrected *P* distributions across all HPs (mean / median / SD). Bold indicates *P* < *α*. *δ* is Cliff’s delta (Method D.4.3.

| Data | Comparison | $n_{\text{HP}}$ | $\Delta_{\text{med}}$ (mean / med / SD) | $\delta$ (mean / med / SD) | $P < \alpha$ | $P$ (mean / med / SD) |
| --- | --- | --- | --- | --- | --- | --- |
| Kinematic | Within- vs. cross-animal, CO | 67 | 0.0051 / 0.0031 / 0.0058 | 0.289 / 0.260 / 0.202 | 45/67 | 0.0741 / 0.0008 / 0.1732 |
| Neural |  | 77 | 0.0066 / 0.0065 / 0.0021 | 0.157 / 0.173 / 0.056 | 44/77 | 0.0448 / 0.0041 / 0.0829 |
| Kinematic | Within- vs. cross-animal, RT | 67 | 0.0091 / 0.0014 / 0.0178 | 0.310 / 0.255 / 0.272 | 16/67 | 0.1885 / 0.1054 / 0.2026 |
| Neural |  | 77 | 0.0024 / 0.0003 / 0.0058 | 0.074 / 0.051 / 0.080 | 0/77 | 0.4067 / 0.4204 / 0.2149 |
| Kinematic | Within- vs. cross-animal, cross-task | 67 | 0.0045 / 0.0015 / 0.0080 | 0.114 / 0.119 / 0.198 | 17/67 | 0.2374 / 0.0673 / 0.3148 |
| Neural |  | 77 | 0.0048 / 0.0030 / 0.0045 | 0.075 / 0.072 / 0.033 | 0/77 | 0.3089 / 0.3239 / 0.1185 |
| Kinematic | Within-task vs. cross-task (within-animal) | 67 | 0.0054 / 0.0033 / 0.0059 | 0.279 / 0.279 / 0.177 | 37/67 | 0.0619 / 0.0013 / 0.1240 |
| Neural | | 77 | 0.0228 / 0.0209 / 0.0114 | 0.323 / 0.323 / 0.028 | 77/77 | $4 \times 10^{-4}$ / $3 \times 10^{-4}$ / $3 \times 10^{-4}$ |
| Kinematic | Within-task vs. cross-epoch (within-animal) | 67 | 0.0327 / 0.0198 / 0.0291 | 0.721 / 0.719 / 0.134 | 67/67 | $< 10^{-5}$ / $< 10^{-5}$ / $< 10^{-5}$ |
| Neural | | 77 | 0.0688 / 0.0651 / 0.0225 | 0.668 / 0.670 / 0.045 | 77/77 | $< 10^{-5}$ / $< 10^{-5}$ / $< 10^{-5}$ |

**Table S3.**
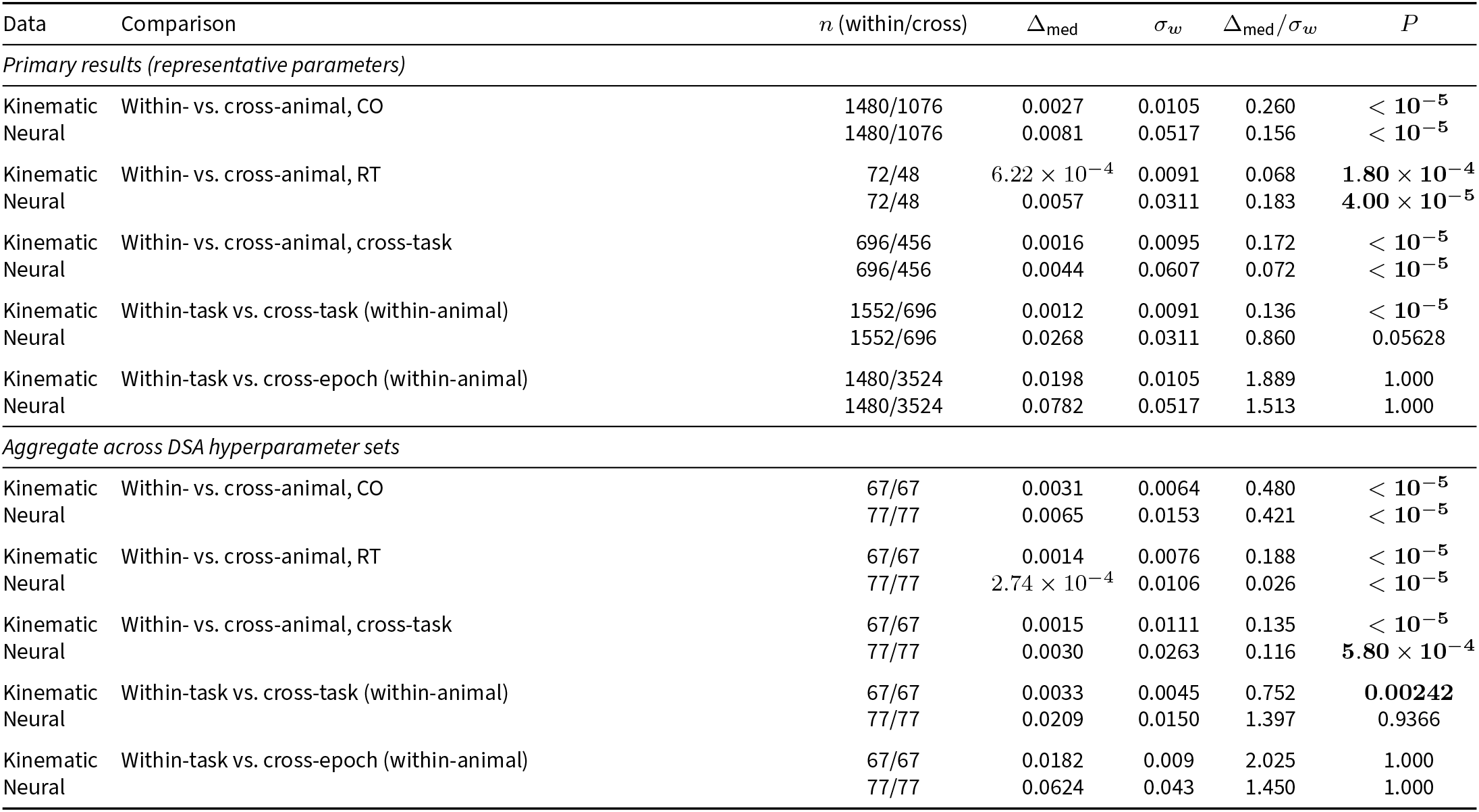
Equivalence tests for DSA distance comparisons. All tests are one-sided bootstrap equivalence tests (*n*_perm_ = 100,000), *P* values are uncorrected; significance is assessed against a Bonferroni-adjusted threshold *α* = 0.05/8 = 6.25 *×* 10^−3^ for *m = 8 sim*ultaneous comparisons. Bold indicates *P* < *α*. The equivalence margin *σ*_*w*_ is the standard deviation of the within-condition distribution. *Δ*_med_/*σ*_*w*_ is the standardized effect size, with values below 1 indicating an effect smaller than the margin.

| Data | Comparison | $n$ (within/cross) | $\Delta_{\text{med}}$ | $\sigma_w$ | $\Delta_{\text{med}}/\sigma_w$ | $P$ |
| --- | --- | --- | --- | --- | --- | --- |
| *Primary results (representative parameters)* |  |  |  |  |  |  |
| Kinematic | Within- vs. cross-animal, CO | 1480/1076 | 0.0027 | 0.0105 | 0.260 | $< 10^{-5}$ |
| Neural | | 1480/1076 | 0.0081 | 0.0517 | 0.156 | $< 10^{-5}$ |
| Kinematic | Within- vs. cross-animal, RT | 72/48 | $6.22 \times 10^{-4}$ | 0.0091 | 0.068 | $1.80 \times 10^{-4}$ |
| Neural | | 72/48 | 0.0057 | 0.0311 | 0.183 | $4.00 \times 10^{-5}$ |
| Kinematic | Within- vs. cross-animal, cross-task | 696/456 | 0.0016 | 0.0095 | 0.172 | $< 10^{-5}$ |
| Neural | | 696/456 | 0.0044 | 0.0607 | 0.072 | $< 10^{-5}$ |
| Kinematic | Within-task vs. cross-task (within-animal) | 1552/696 | 0.0012 | 0.0091 | 0.136 | $< 10^{-5}$ |
| Neural |  | 1552/696 | 0.0268 | 0.0311 | 0.860 | 0.05628 |
| Kinematic | Within-task vs. cross-epoch (within-animal) | 1480/3524 | 0.0198 | 0.0105 | 1.889 | 1.000 |
| Neural |  | 1480/3524 | 0.0782 | 0.0517 | 1.513 | 1.000 |
| *Aggregate across DSA hyperparameter sets* |  |  |  |  |  |  |
| Kinematic | Within- vs. cross-animal, CO | 67/67 | 0.0031 | 0.0064 | 0.480 | $< 10^{-5}$ |
| Neural | | 77/77 | 0.0065 | 0.0153 | 0.421 | $< 10^{-5}$ |
| Kinematic | Within- vs. cross-animal, RT | 67/67 | 0.0014 | 0.0076 | 0.188 | $< 10^{-5}$ |
| Neural | | 77/77 | $2.74 \times 10^{-4}$ | 0.0106 | 0.026 | $< 10^{-5}$ |
| Kinematic | Within- vs. cross-animal, cross-task | 67/67 | 0.0015 | 0.0111 | 0.135 | $< 10^{-5}$ |
| Neural | | 77/77 | 0.0030 | 0.0263 | 0.116 | $5.80 \times 10^{-4}$ |
| Kinematic | Within-task vs. cross-task (within-animal) | 67/67 | 0.0033 | 0.0045 | 0.752 | **0.00242** |
| Neural |  | 77/77 | 0.0209 | 0.0150 | 1.397 | 0.9366 |
| Kinematic | Within-task vs. cross-epoch (within-animal) | 67/67 | 0.0182 | 0.009 | 2.025 | 1.000 |
| Neural |  | 77/77 | 0.0624 | 0.043 | 1.450 | 1.000 |

**Table S4.** Robustness of equivalence tests across DSA hyperparameters. Fraction of HP settings where the per-cell equivalence test is significant at *α* = 0.05/8 (Bonferroni-corrected), with uncorrected *P* value distributions summarized across all HPs (mean / median / SD). The ratio of *Δ*_med_ and *σ*_*w*_ is the standardized effect size.

| Data | Comparison | $n_{\text{HP}}$ | $\Delta_{\text{med}}$ (mean / med / SD) | $\Delta_{\text{med}}/\sigma_w$ (mean / med / SD) | $P < \alpha$ | $P$ (mean / med / SD) |
| --- | --- | --- | --- | --- | --- | --- |
| Kinematic | Within- vs. cross-animal, CO | 67 | 0.0051 / 0.0031 / 0.0058 | 0.496 / 0.381 / 0.529 | 61/67 | 0.0492 / $< 10^{-5}$ / 0.2080 |
| Neural | | 77 | 0.0066 / 0.0065 / 0.0021 | 0.170 / 0.181 / 0.073 | 77/77 | $< 10^{-5}$ / $< 10^{-5}$ / $< 10^{-5}$ |
| Kinematic | Within- vs. cross-animal, RT | 67 | 0.0091 / 0.0014 / 0.0178 | 0.696 / 0.466 / 0.867 | 26/67 | 0.2442 / 0.0166 / 0.3876 |
| Neural | | 77 | 0.0024 / 0.0003 / 0.0058 | 0.048 / 0.013 / 0.190 | 76/77 | $5 \times 10^{-4}$ / $4 \times 10^{-5}$ / 0.0011 |
| Kinematic | Within- vs. cross-animal, cross-task | 67 | 0.0045 / 0.0015 / 0.0080 | 0.306 / 0.243 / 0.403 | 55/67 | 0.0634 / $< 10^{-5}$ / 0.1934 |
| Neural | | 77 | 0.0048 / 0.0030 / 0.0045 | 0.098 / 0.085 / 0.065 | 77/77 | $< 10^{-5}$ / $< 10^{-5}$ / $< 10^{-5}$ |
| Kinematic | Within-task vs. cross-task (within-animal) | 67 | 0.0054 / 0.0033 / 0.0059 | 0.731 / 0.511 / 0.984 | 48/67 | 0.1822 / $< 10^{-5}$ / 0.3649 |
| Neural |  | 77 | 0.0228 / 0.0209 / 0.0114 | 0.898 / 0.861 / 0.176 | 16/77 | 0.2682 / 0.0699 / 0.3351 |
| Kinematic | Within-task vs. cross-epoch (within-animal) | 67 | 0.0327 / 0.0198 / 0.0291 | 3.663 / 2.171 / 4.567 | 2/67 | 0.9655 / 1.0000 / 0.1735 |
| Neural | | 77 | 0.0688 / 0.0651 / 0.0225 | 1.571 / 1.546 / 0.253 | 0/77 | 1.0000 / 1.0000 / $4 \times 10^{-6}$ |

**Table S5.** Session-date permutation tests for DSA-distance drift. Statistical tests for whether the slope of DSA distance against date difference differs from zero (one-sided, *n*_perm_ = 2,000). *P* values are uncorrected. Significance is assessed against a Bonferroni-adjusted threshold *α* = 0.05/8 = 6.25 *×* 10^−3^ for *m* = 8 simultaneous comparisons. Bold indicates *P* < *α*. Slope is in units of Δd_DSA_ per day. For the primary results we additionally report the 95% bootstrap confidence interval on the ordinary-least-squares slope (resampling session pairs, *n*_boot_ = 100,000), and the *R*^2^ coefficient of determination between DSA distance and inter-session date difference.

| Data | Metric | Task | $n$ | Slope | 95% CI | $R^2$ | $P$ |
| --- | --- | --- | --- | --- | --- | --- | --- |
| *Primary results* |  |  |  |  |  |  |  |
| Kinematic | DSA | CO | 1480 | $2.26 \times 10^{-6}$ | $[1.04, 3.43] \times 10^{-6}$ | 0.0041 | 0.2629 |
| Neural | | | 1480 | $1.28 \times 10^{-5}$ | $[0.77, 1.79] \times 10^{-5}$ | 0.0055 | 0.2994 |
| Kinematic | DSA | RT | 72 | $3.41 \times 10^{-6}$ | $[-3.14, 10.06] \times 10^{-6}$ | 0.0072 | 0.4543 |
| Neural | | | 72 | $8.46 \times 10^{-5}$ | $[6.64, 10.28] \times 10^{-5}$ | **0.384** | **$9.995 \times 10^{-4}$** |
| Kinematic | DSA | Cross-task | 696 | $-1.17 \times 10^{-6}$ | $[-2.86, 0.53] \times 10^{-6}$ | 0.0024 | 0.6162 |
| Neural | | | 696 | $7.13 \times 10^{-5}$ | $[6.02, 8.24] \times 10^{-5}$ | **0.215** | **$9.995 \times 10^{-4}$** |
| Neural | PAVF-aligned DMD prediction error | CO | 1480 | $2 \times 10^{-6}$ | $[-1.0, 1.5] \times 10^{-5}$ | $< 10^{-3}$ | 0.6830 |
| Neural | PAVF-aligned DMD prediction error | RT | 72 | $5.54 \times 10^{-4}$ | $[3.34, 7.74] \times 10^{-4}$ | **0.158** | **$< 10^{-4}$** |
| *Aggregate across DSA hyperparameter sets* |  |  |  |  |  |  |  |
| Kinematic | DSA | CO | 67 | $1.29 \times 10^{-6}$ | $[5.21 \times 10^{-7}, 2.04 \times 10^{-6}]$ | 0.0032 | 0.02806 |
| Neural | | | 77 | $9.27 \times 10^{-6}$ | $[7.81 \times 10^{-6}, 1.18 \times 10^{-5}]$ | 0.0054 | 0.1890 |
| Kinematic | DSA | RT | 67 | $1.81 \times 10^{-7}$ | $[-8.60 \times 10^{-7}, 8.40 \times 10^{-7}]$ | 0.0107 | 0.2433 |
| Neural | | | 77 | $5.77 \times 10^{-5}$ | $[4.94 \times 10^{-5}, 6.53 \times 10^{-5}]$ | **0.397** | **$5.20 \times 10^{-4}$** |
| Kinematic | DSA | Cross-task | 67 | $-3.40 \times 10^{-7}$ | $[-1.16 \times 10^{-6}, 1.20 \times 10^{-6}]$ | 0.0112 | 0.6920 |
| Neural | | | 77 | $5.34 \times 10^{-5}$ | $[4.44 \times 10^{-5}, 6.06 \times 10^{-5}]$ | **0.234** | **$< 10^{-5}$** |

**Table S6.** Robustness of slope tests across DSA hyperparameters. Fraction of HP settings where the hyperparameter permutation test on slopes is significant at *α* = 0.05/8 (Bonferroni-corrected), with hyperparameter distribution of uncorrected *P* values summarized as (mean / median / SD).

| Data | Task | $n_{\text{HP}}$ | Slope (mean / med / SD) | $R^2$ (mean / med / SD) | $P < \alpha$ | $P$ (mean / med / SD) |
| --- | --- | --- | --- | --- | --- | --- |
| Kinematic | CO | 67 | $1.80 \times 10^{-6} / 1.29 \times 10^{-6} / 3.25 \times 10^{-6}$ | 0.0120 / 0.0032 / 0.0215 | 7/67 | 0.4155 / 0.3593 / 0.3383 |
| Neural | | 77 | $1.35 \times 10^{-5} / 9.27 \times 10^{-6} / 1.05 \times 10^{-5}$ | 0.0069 / 0.0054 / 0.0047 | 0/77 | 0.3226 / 0.2994 / 0.2020 |
| Kinematic | RT | 67 | $1.00 \times 10^{-6} / 1.81 \times 10^{-7} / 9.64 \times 10^{-6}$ | 0.0304 / 0.0107 / 0.0467 | 2/67 | 0.4568 / 0.4543 / 0.3301 |
| Neural | | 77 | $6.55 \times 10^{-5} / 5.77 \times 10^{-5} / 3.55 \times 10^{-5}$ | 0.3846 / 0.3967 / 0.1521 | 75/77 | 0.0104 / 0.0010 / 0.0674 |
| Kinematic | Cross-task | 67 | $-2.78 \times 10^{-8} / -3.40 \times 10^{-7} / 5.51 \times 10^{-6}$ | 0.0325 / 0.0112 / 0.0518 | 7/67 | 0.4306 / 0.3883 / 0.3181 |
| Neural | | 77 | $6.13 \times 10^{-5} / 5.34 \times 10^{-5} / 3.38 \times 10^{-5}$ | 0.2316 / 0.2344 / 0.0205 | 77/77 | 0.0008 / 0.0005 / 0.0005 |

**Table S7.**
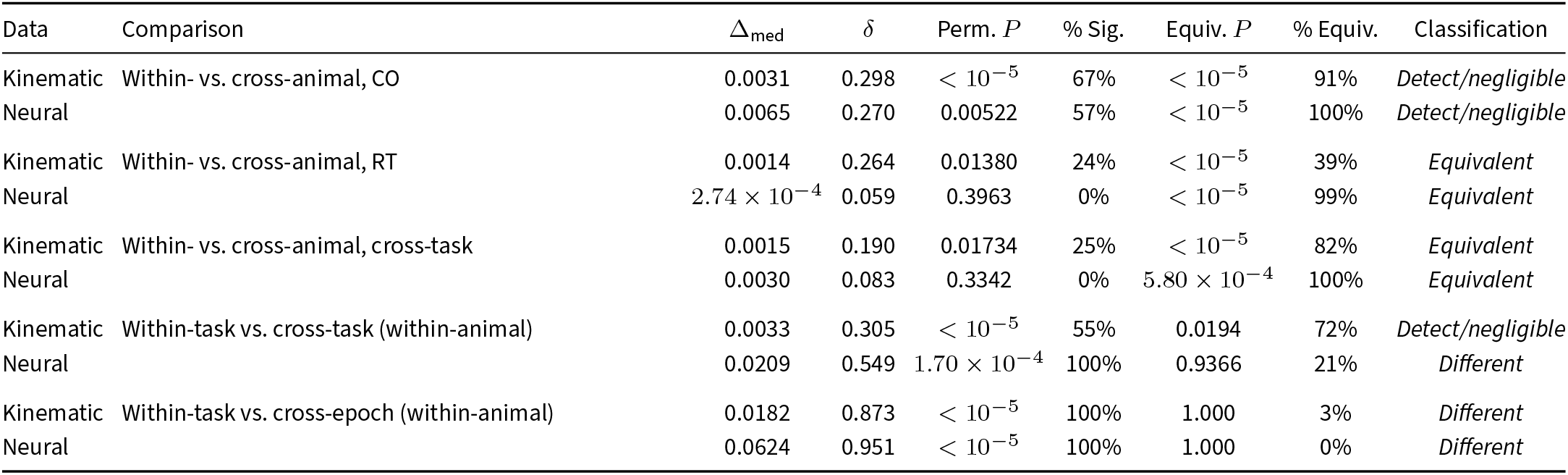
Summary of DSA distance comparisons across hyperparameters. Each comparison is classified using two tests (threshold, not p-values, are Bonferroni-corrected for 8 simultaneous comparisons, *α* = 0.05/8): a one-sided session-label permutation test for significance and a bootstrap equivalence test with margin *σ*_*w*_ (within-condition SD). Aggregate columns pool across all HP settings; % Sig. and % Equiv. report the fraction of individual HP settings where the single-HP permutation / equivalence test is significant at *α* = 0.05/8 (Bonferroni-corrected for 8 simultaneous comparisons). Equivalent: not significant and equivalence confirmed. Different: significant and equivalence rejected. Detect/negligible: significant but equivalence confirmed, indicating a real but practically negligible effect.

## G Derivation of the stochastic Koopman Operator for the diffusive ring attractor and bounded line attractor / cut ring

### G.1 System dynamics

We consider a ring attractor system in polar coordinates for simplicity. The state is defined by the radial amplitude *r* ∈ ℝ^+^ and the phase *θ* ∈ [0, *L*], subject to periodic boundary conditions such that *θ* ~ *θ* + *L*.

The stochastic differential equations (SDEs) describing the evolution over an infinitesimal time step *dt* are:

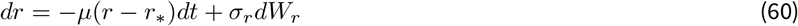

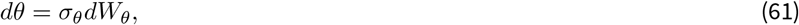

where *dW*_*r*_ and *dW*_*θ*_ are independent Wiener process increments satisfying E[*dW*] = 0 and E[*dW* ^2^] = *dt*.

**Table S8.**
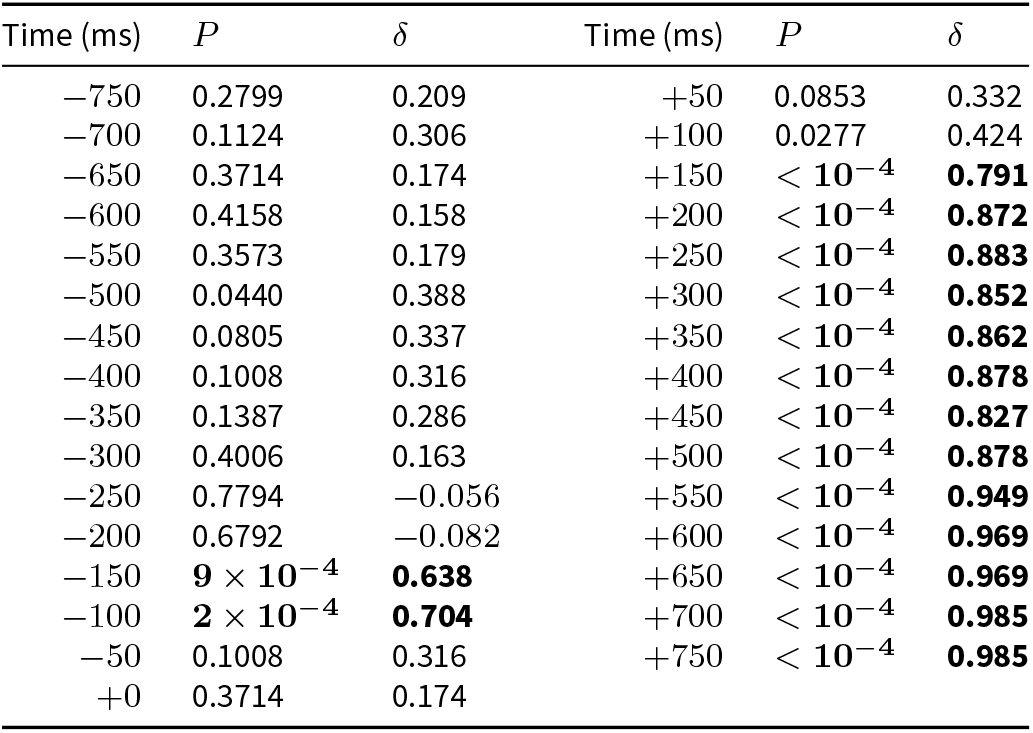
Cross-region versus within-region DSA distance across time windows (M1 and PMd). One-sided Mann–Whitney U tests comparing cross-region DSA distance to the within-region baseline, computed independently in each 50 ms window relative to movement onset. Each entry is the median across 8 DSA parameter sets. P values are uncorrected; significance is assessed against a Bonferroni-adjusted threshold *α* = 0.05/31 = 1.61 *×* 10^−3^ for *m = 31 si*multaneous window comparisons. Bold indicates *P* < *α*. *δ* is Cliff’s delta (Methods Sec. D.4.3); positive values indicate larger cross-region than within-region distances. These are the tests summarised by the significance bars in Fig. 4n.

### G.2 Deriving the Koopman generator

The Koopman Operator describes the evolution of observable functions Ψ(*r, θ*) along the trajectories of the system. In the stochastic setting, the generator ℒ is defined by the expected rate of change of the observable:

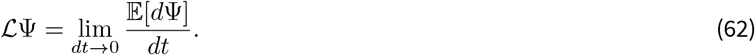

We perform a Taylor expansion of the increment *d*Ψ = Ψ(*r* + *dr, θ* + *dθ*) − Ψ(*r, θ*) up to second order, retaining terms of order *dt* (since *dW* ~ 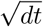 implies *dW* ^2^ ~ *dt*):

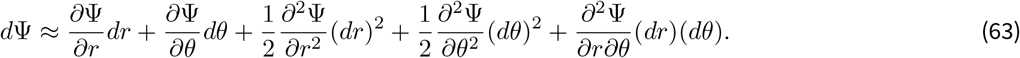

Substituting the SDEs and applying Itô rules (*dt* · *dW* → 0, *dW* ^2^ → *dt*):

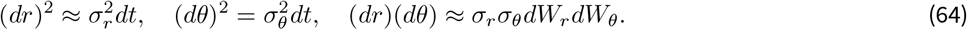

Taking the expectation E[·], the linear noise terms and the cross-term vanish (independence implies E[*dW*_*r*_*dW*_*θ*_] = 0). We are left with the drift and diffusion components:

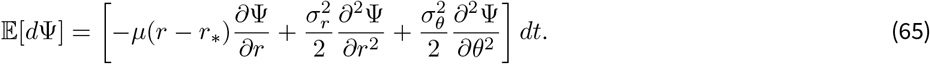

Dividing by *dt* yields the infinitesimal generator:

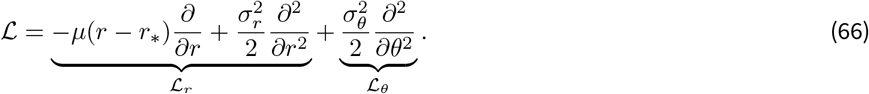

**Figure S12.**
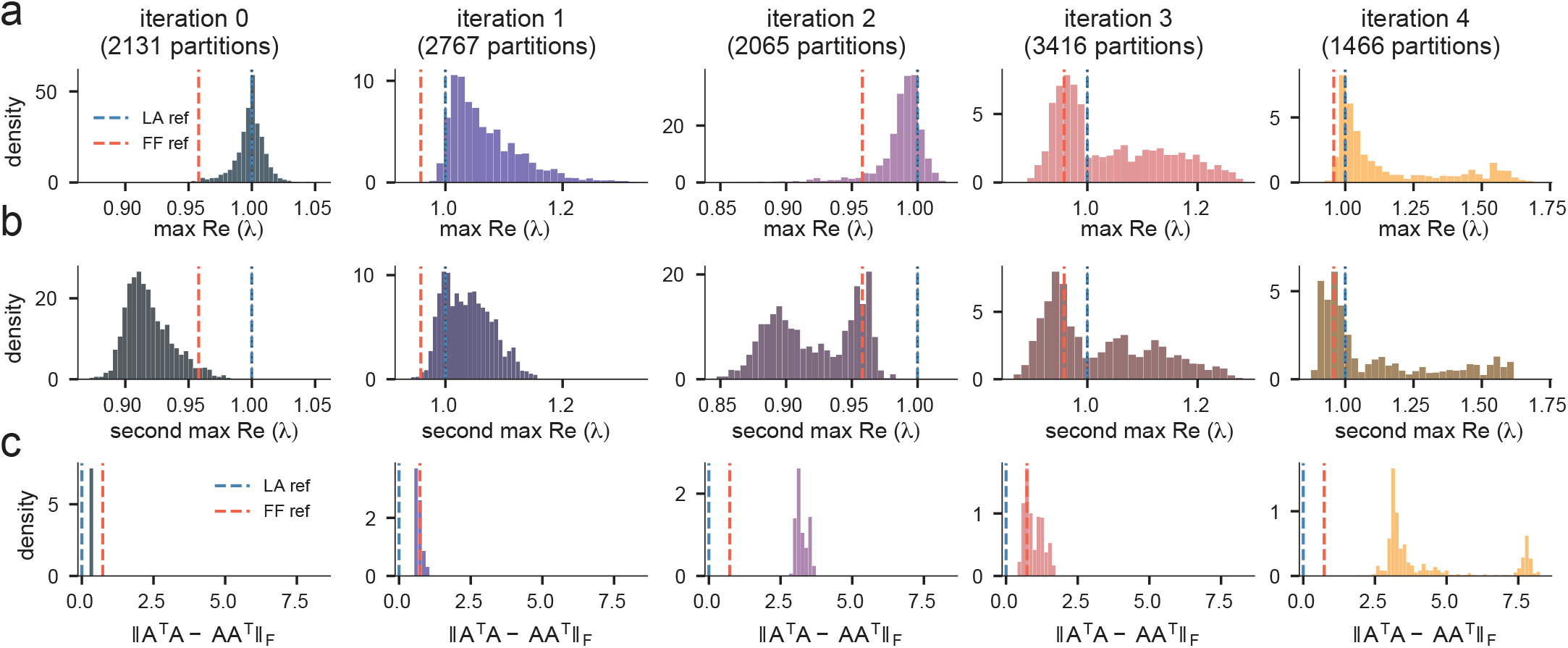
Jacobian dynamics analysis of recurrent matrices reveals diverse mechanisms of integration. Each ReLU RNN can be described as locally linear, with local recurrent dynamics being defined by taking only the rows and columns in the recurrent weight matrix of neurons that are active (*x* > 0). We drove the networks with within-training-distribution inputs, extracted each Jacobian matrix along the trajectory, and computed the eigenvalues and non-normality scores of these matrices. **(a)** Distribution of max real component of all Jacobian eigenvalues. Blue vertical line denotes the max eigenvalue for a sample line attractor model. Red denotes the max eigenvalue for a sample feedforward chain. **(b)** Distribution of second largest real component of all Jacobian eigenvalues. Iteration 0 only has 1 eigenvalue near 1, while Iteration 1 has 2, indicating the presence of a plane attractor. **(c)** Non-normality score distribution for each model. Feedforward chain models are traditionally highly non-normal, as seen in Iterations 2-4.

### G.3 Spectral decomposition via separation of variables

We seek eigenfunctions Ψ and eigenvalues *λ* satisfying ℒΨ = *λ*Ψ. Since ℒ = ℒ_*r*_ + ℒ_*θ*_, we use separation of variables Ψ(*r, θ*) = *ϕ*(*r*)*g*(*θ*). The eigenvalue equation separates into:

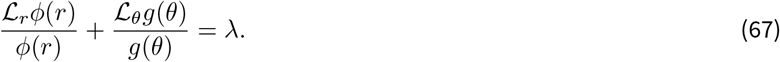

This implies *λ* = *λ*_*r*_ + *λ*_*θ*_.

#### G.3.1 Angular eigenfunctions

The angular dynamics follow diffusion on the interval [0, *L*] with periodic boundary conditions. The eigenvalue problem is:

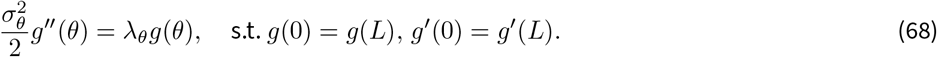

The general solution for spatial diffusion is of the form *g*(*θ*) = *e*^*ik*θ^. Applying the periodicity constraint:

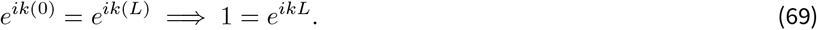

This condition quantizes the wavenumber *k*. For the equality to hold, the argument of the exponential must be a multiple of 2*πi*:

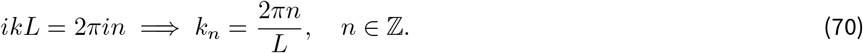

**Figure S13.**
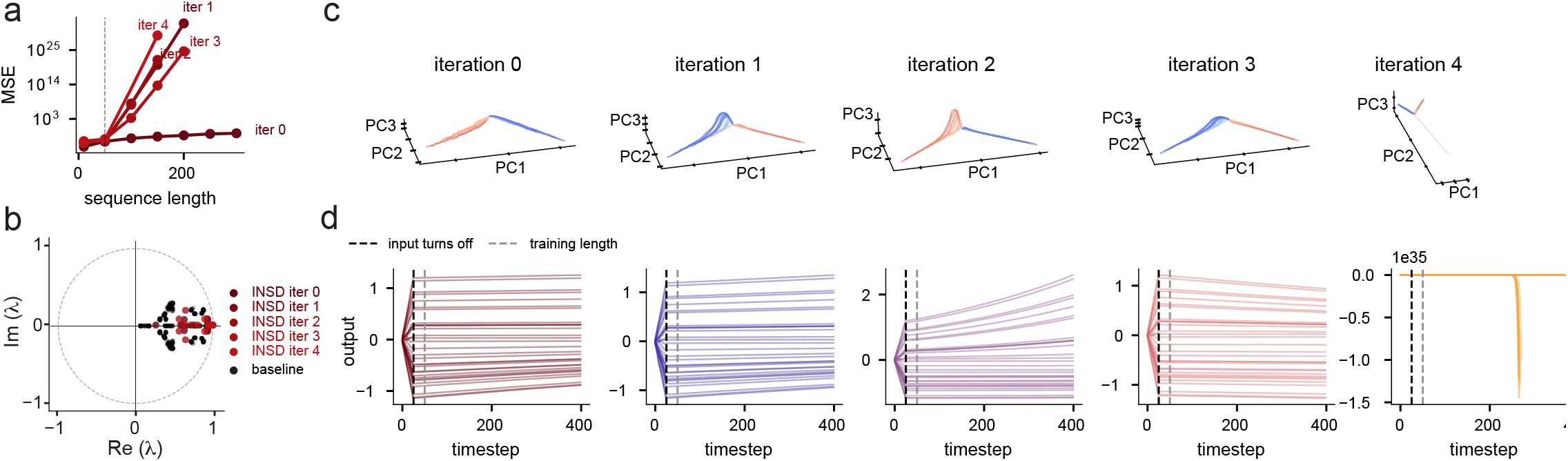
Iterative Neural Similarity Deflation (INSD) learns only unstable line attractors on the integration task. **(a)** MSE of each trained model iteration for increasing task lengths. Dotted line indicates the training interval of 50 timesteps (same as ∂DSA regularization). **(b)** DMD eigenvalues of the final trained solution for each iteration. **(c)** 3D trajectories of each iteration in the top 3 PCs using the same input as Fig. 5k **(d)** Relaxation dynamics of each iteration trained with INSD as in Fig. 5i.

**Figure S14.**
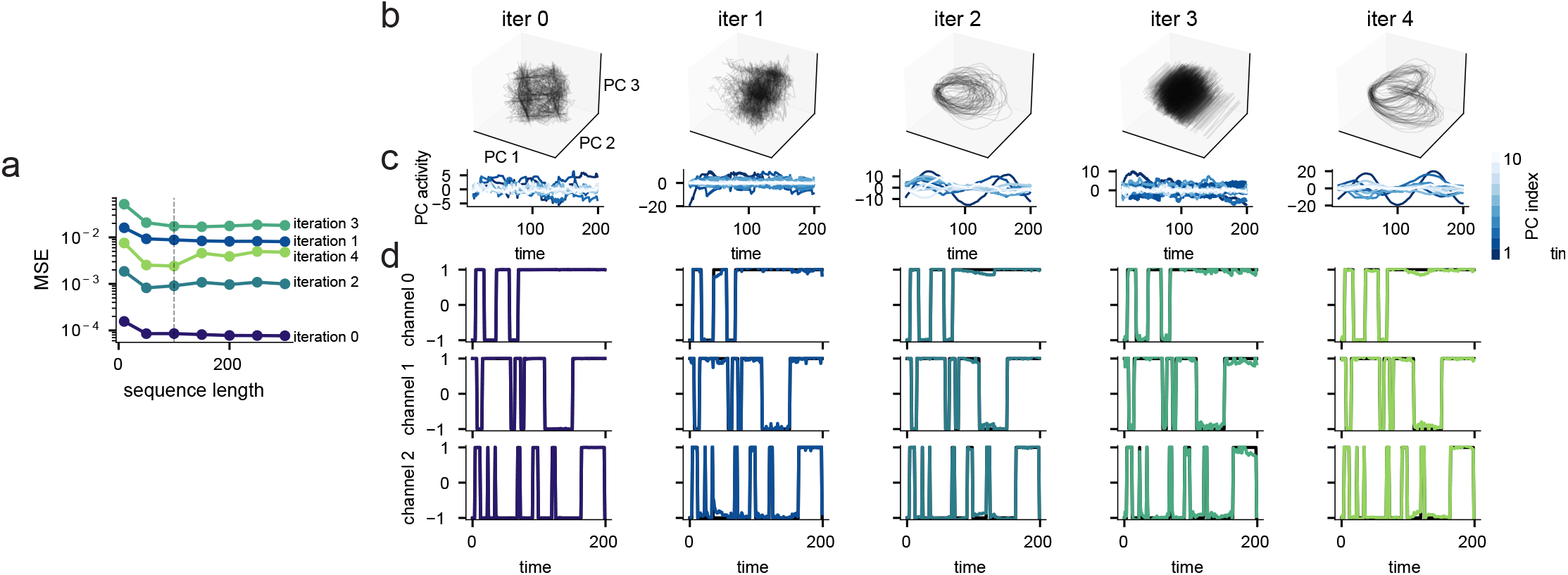
∂DSA applied to the 3-bit flipflop task finds novel solutions. **(a)** MSE of each trained model iteration for increasing task lengths. Dotted line indicates training interval of 100. **(b)** Trajectories of each iteration visualized in the top 3 PCs. The first unregularized iteration learns the cube of fixed points, as is canonical. The second and fourth models learn a noncanonical version of the cube. The third learns the oscillating solution as in (*Qian and Pehlevan, 2026*). The fifth learns a more complex oscillator made up of two lobes. **(c)** Trajectories of individual PCs for each iteration as in (b), for a sample trial of length 200. **(d)** Sample output for each iteration for trajectories as in (b,c).

**Figure S15.**
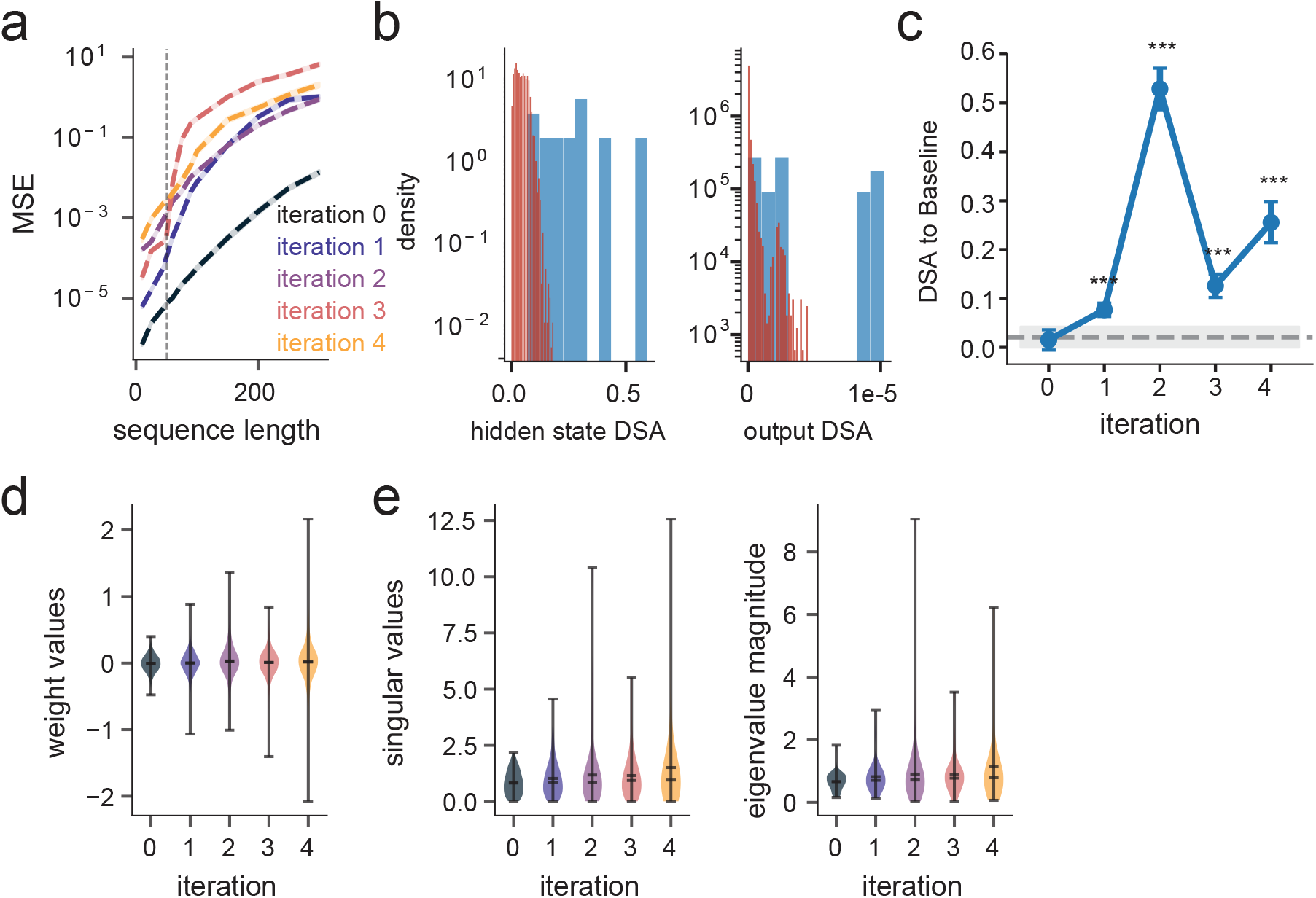
∂DSA regularized models have diverse generalization capacities, baseline DSA distances, and weight distribution properties. **(a)** Length generalization error (mean squared error, MSE). We drove models with varying sequences of increasing length up to 300 time steps, and measured the MSE of the output. Dotted vertical line indicates training sequence length. **(b)** Distribution of DSA distances applied on the hidden state (left) and output (right). Blue indicates the DSA distances between all pairs of iteratively learned solutions, red indicates the DSA distance between all pairs of baseline solutions. **(c)** Average DSA distance to the baseline distribution (Circles in Fig. 5g) across iterations. Bars indicate standard deviation, asterisks indicate significance test against within-basin distancesy *p* < 0.001 (Mann-Whitney U-test). **(d)** Distribution of recurrent weight matrix values across iterations. Center horizontal line indicates mean. **(e)** Distribution of singular values (left) and eigenvalues (right) of recurrent weight matrices.

The eigenfunctions are the Fourier modes on the scaled interval:

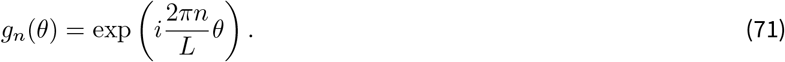

Substituting *g*_*n*_ back into the operator yields the eigenvalues:

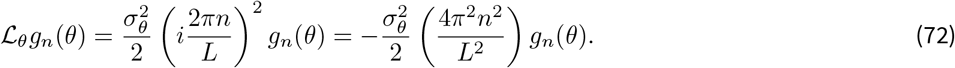

Thus, the angular eigenvalues are:

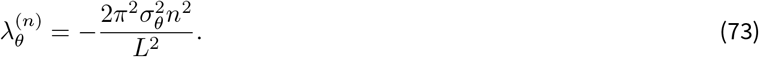

#### G.3.2 Radial eigenfunctions

The radial dynamics describe an Ornstein-Uhlenbeck process centered at *r*_∗_. Defining the centered variable *x* = *r* − *r*_∗_, the operator is:

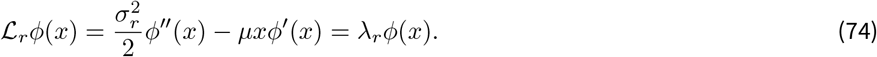

We search for polynomial solutions *ϕ*_*k*_(*x*) of degree *k*. The highest power *x*^*k*^ is mapped by the drift term −*µx*∂_*x*_ to −*kµx*^*k*^, while the diffusion term lowers the order. Matching coefficients implies:

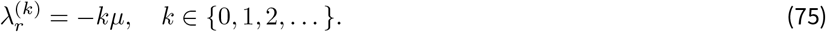

The eigenfunctions are the probabilist’s Hermite polynomials *H*_*k*_, normalized by the standard deviation.

### G.4 Complete solution

The full spectrum of the stochastic Koopman Operator on the ring of length *L* is given by the sum of radial and angular modes. The eigenfunctions Ψ_*k,n*_ and eigenvalues *λ*_*k,n*_ are:

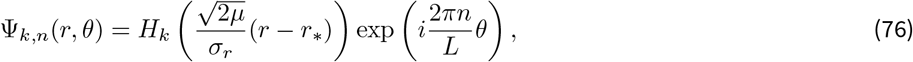

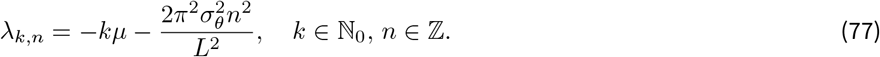

### G.5 Derivation for the bounded line (cut ring) attractor

#### G.5.1 System definition

We consider a dynamical system on a bounded angular interval *θ* ∈ [0, *L*]. The dynamics are governed by radial relaxation and angular diffusion, subject to **reflecting boundaries** at the endpoints.

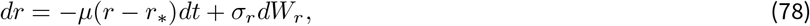

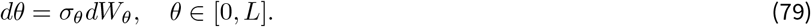

#### G.5.2 Physical interpretation of boundary conditions

The reflecting boundaries imply that any state hitting the end of the line at *θ* = 0 or *θ* = *L* is instantaneously reflected back into the domain. In terms of the Koopman Operator (which evolves observable functions Ψ), this requires Neumann Boundary Conditions:

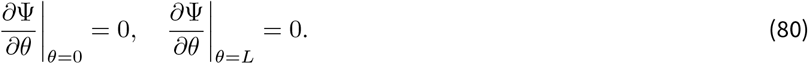

This condition arises from the symmetry of reflection. For the observable Ψ to be consistent with this physical symmetry, it must be an **even function** across the boundary (Ψ(*θ*) = Ψ(−*θ*)).

#### G.5.3 Angular eigenfunctions

The angular operator is 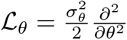. Solving the eigenvalue equation ℒ_*θ*_*g* = *λ*_*θ*_*g* with Neumann conditions selects the cosine modes (standing waves) rather than the complex exponentials (traveling waves) found in the ring attractor.

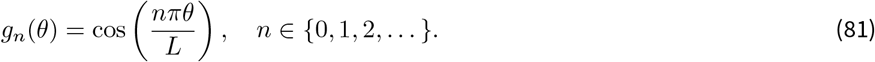

The associated eigenvalues are:

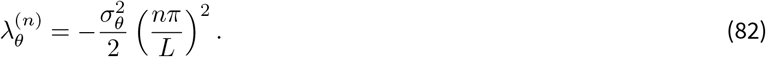

### G.6 Differences between Koopman eigenspectra

#### G.6.1 Ring attractor

For *x* ∈ [0, *L*) with *ϕ*(*x*) = *ϕ*(*x* + *L*), the spectrum is given by:

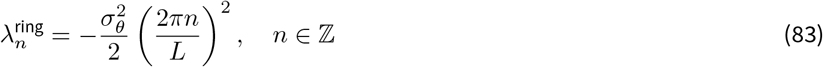

Crucially, for *n* ≥ 1, the spectrum is degenerate with multiplicity 2. The eigenspace is spanned by the orthogonal pair:

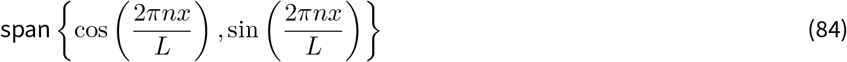

This degeneracy is a reflection of *S*^1^ rotational symmetry.

#### G.6.2 Line attractor

For *x* ∈ [0, *L*] with *ϕ*^*′*^(0) = *ϕ*^*′*^(*L*) = 0, the spectrum is:

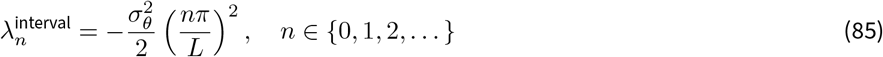

Here, the spectrum is simple (multiplicity 1) for all *n*, as the boundary conditions break the symmetry that permits shifted eigenfunctions. The cut ring spectrum therefore contains the ring spectrum on its even indices, while differing on its odd indices. Additionally, the ring spectrum has two eigenvalues on those shared indices, while the cut ring spectrum is simple.

## H Separation of timescales via delay embedding

We investigate the properties of *s*-step linear models (Koopman Operators) fitted to dynamical systems exhibiting multiple timescales. Specifically, we analyze the effect of the prediction horizon *s* on the spectral composition of the learned operator. We show that increasing *s* enforces the separation of slow (persistent) dynamics from fast (transient) dynamics or noise. This analysis provides a theoretical basis for comparing multiple timescales with DSA.

### H.1 Problem setup

Consider a scalar observable *x*_*t*_ ∈ ℝ composed of an additive superposition of a “slow” signal *m*_*t*_ and a “fast” signal *η*_*t*_:

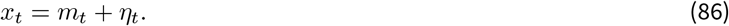

We assume *m*_*t*_ and *η*_*t*_ are zero-mean, stationary processes generated by distinct linear dynamics (or linearized modes of a nonlinear system) characterized by their eigenvalues:

- **Slow Dynamics:** Governed by eigenvalues *λ*_*m*_ *w*ith characteristic timescale *τ*_*m*_, such that 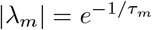.
- **Fast Dynamics:** Governed by eigenvalues *λ*_*η*_ with characteristic timescale *τ*_*η*_, such that 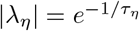.

**Definition 1** (Separation of timescales). *The system exhibits a separation of timescales if τ*_*m*_ > *τ*_η_ > 0, *implying* |*λ*_*m*_| > |*λ*_η_|.

### H.2 Delay embeddings geometrically separate timescales

A direct linear model on the scalar *x*_*t*_ is insufficient to separate the additive components *m*_*t*_ and *η*_*t*_ without bias. To resolve this, we lift the system into a high-dimensional state space using delay coordinates.

Define the delay vector (embedding) **x**_*t*_ ∈ ℝ^d^ for a window length *d*:

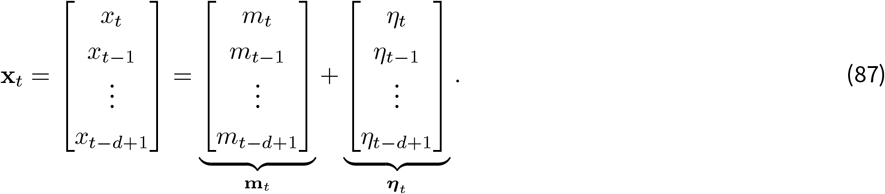

**Lemma 2** (Asymptotic subspace orthogonality). *Let m*_*t*_ *and η*_*t*_ *be jointly stationary, ergodic, zero-mean processes that are uncorrelated at every lag, i*.*e*. E[*m*_*t*_ *η*_*t*−*k*_] = 0 *for all k* ∈ ℤ. *Then, as the embedding dimension d* → ∞, *every entry of the normalized cross-Gram block between the trajectories* **m**_*t*_ *and* ***η***_t_ *vanishes:*

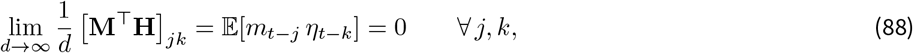

*where* **M** *and* **H** *collect the delay trajectories of m*_*t*_ *and η*_*t*_ *respectively. Consequently the two spanned subspaces become orthogonal*.

#### *Proof*.

By joint stationarity and ergodicity, each normalized inner product converges almost surely to the corresponding cross-covariance E[*m*_*t*−*j*_*η*_*t*−*k*_] = *R*_*mη*_(*j* − *k*), which is zero for all lags by hypothesis. Orthogonality of the spans follows since every pairwise inner product between basis directions vanishes in the limit.

#### Remark 1.

*This orthogonality allows us to treat the evolution of the embedded state* **x**_*t*_ *as the independent evolution of its components. The covariance matrix* 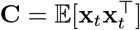 *approximately block-diagonalizes into slow and fast subspaces (King, 1986)*.

### H.3 Spectral analysis of the s-step operator

We fit a linear operator **K**_s_ to predict the embedded state *s* steps into the future:

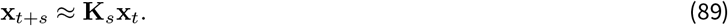

In the exact-separation limit of Lemma 2, the spectrum of **K**_*s*_ is the union of the spectra of the slow and fast dynamics raised to the power *s*:

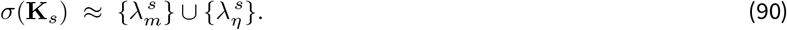

We quantify the filtering effect of the horizon *s* using the Spectral Ratio.

#### **Proposition H.1** (Exponential spectral filtering).

*Let ρ*(*s*) *be the ratio of the magnitude of the slow mode to that of the fast mode in the spectrum of* **K**_s_. *Then ρ*(*s*) *diverges exponentially in s*.

#### *Proof*.

The amplitude of a mode *λ* after *s* steps scales as |*λ*|^s^. We define the ratio *ρ*(*s*):

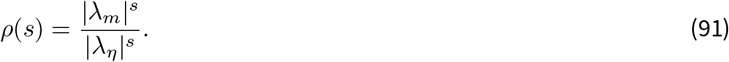

Substituting the timescale definitions |*λ*| = *e*^−1/τ^ :

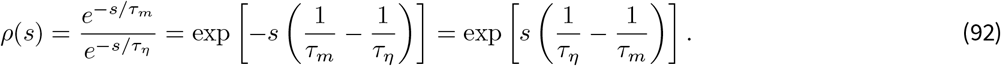

Let 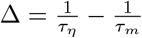. Since *τ*_*m*_ > *τ*_*η*_, we have Δ > 0. Thus:

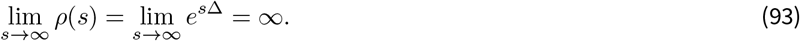

## I Theoretical model for DSA between multiple sleep stages across timescales

### I.1 System dynamics

In order to better align with the head direction circuit data, we extend the ring attractor to include diffusion of *θ* with an integrated Ornstein-Uhlenbeck process. The state is now (*r, θ, ω*) ∈ ℝ^+^ × [0, *L*) × ℝ.

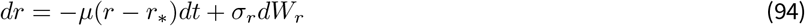

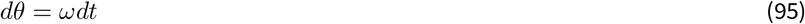

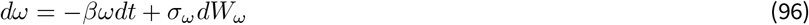

where *dW*_*r*_ and *dW*_ω_ are independent Wiener processes.

### I.2 Infinitesimal generator

We apply Itô’s Lemma to a continuously differentiable observable Ψ(*r, θ, ω*). Keeping terms up to order *dt*, and noting the quadratic variations 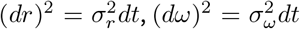, and (*dθ*)^2^ = (*dθ*)(*dr*) = (*dθ*)(*dω*) = 0, the expected rate of change yields the generator ℒ = ℒ_*r*_ + ℒ_*θ,ω*_:

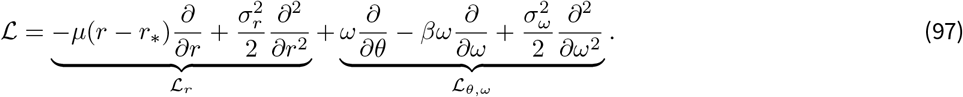

### I.3 Spectral decomposition

Because the radial dynamics do not couple to (*θ, ω*), we employ separation of variables Ψ(*r, θ, ω*) = *ϕ*(*r*)*ψ*(*θ, ω*). The eigen-value equation splits, yielding *λ* = *λ*_*r*_ + *λ*_*θ,ω*_.

#### I.3.1 Radial eigenfunctions

The operator ℒ_*r*_ matches Section G. The solution requires integer constants *m*_1_ ∈ N_0_:

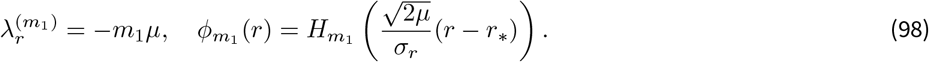

#### I.3.2 Coupled angular-velocity eigenfunctions

Assuming a Fourier basis for the bounded phase coordinate, let 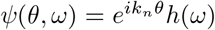 where the periodic boundary conditions enforce wavenumber 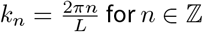:

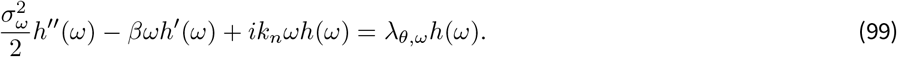

To decouple the linear *ωh* term, we apply the transformation 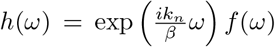. Grouping terms and canceling the dependence on *ωf* (*ω*) yields:

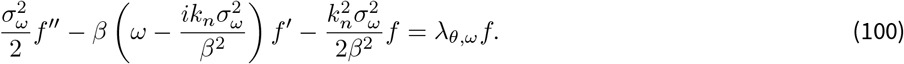

We shift to a new coordinate 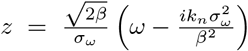, mapping the ODE exactly to the standard Hermite differential equation:

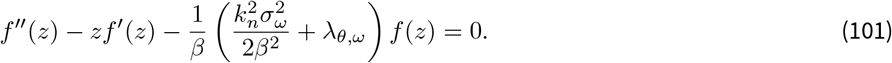

For polynomial solutions bounded under the invariant measure, the constant factor must evaluate to an integer *m*_2_ ∈ N_0_. Solving for *λ*_θ,ω_:

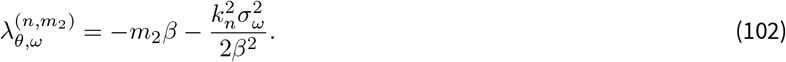

### I.4 Complete spectrum

The fully resolved spectrum is characterized by the indices (*m*_1_, *m*_2_, *n*) ∈ N_0_ × N_0_ × ℤ:

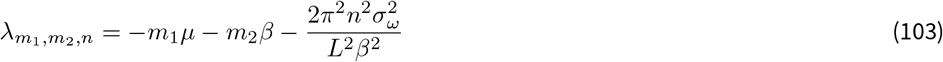

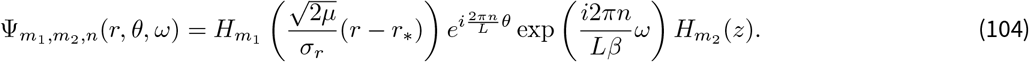

### I.5 Theoretical DSA distance on different sleep stages

We use the above derivation to produce a theoretical prediction of the DSA distance for various sleep stages in the head direction circuit. These are defined by the parameters *µ, β*, and *σ*_*ω*_, which govern the radial decay rate onto the ring, the autocorrelation rate of the angular velocity, and the variance of the angular velocity inputs, respectively. Each sleep stage (Wake, REM, SWS) has different relative values for each parameter:

- Wake: large *µ* (strong radial decay), small *β* (long timescale angular velocity), large *σ*_*ω*_ (large variance angular sweeps).
- REM: large *µ*, large *β*, small *σ*_*ω*_ (together, the latter two drive small diffusive movements along the ring)
- SWS: small *µ* (collapsed ring, modeled by weak radial decay), small *β*, large *σ*_*ω*_. The collapse of the ring in the data is driven by a slow-wave oscillation that modulates ring amplitude in the radial direction (*Chaudhuri et al., 2019*). We averaged over the fast oscillations, just as the data is coarse-grained in time, to recover the slow radial parameter *µ*.

These correspond to different timescales 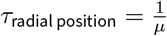 and 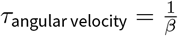 in Fig. 3k.

Above, we derived the eigenvalues for the infinitesimal generator. These map to empirical eigenvalues for discrete timesteps via exponentiation: *λ*_Δ*t*_ = exp(*λ*Δ*t*). The core dynamics are captured by the *principal* eigenvalues, defined by the indices {(1, 0, 0), (0, 1, 0), (0, 0, 1)} for (*m*_1_, *m*_2_, *n*). Taking *L* = 2*π*, we write 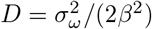 for the effective angular diffusion constant, making the three principal eigenvalues for each state *i*:

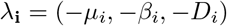

Under this formulation, the *k*-step eigenvalue in discrete time is defined naturally via 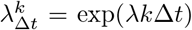. Lastly, because the eigenvalues are entirely real, the DSA distance resolves to the 1D Wasserstein distance, which has a closed-form solution via sorting each eigenvalue list by magnitude. The results on the shape of the distance curves do not depend on the ordering of our eigenvalues.

This formulation allows us to derive a simple analytical description of the DSA curves across multiple prediction horizons *k*Δ*t*. Each matched pair of eigenvalues between systems contributes a term 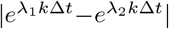, whose shape only depends on whether both rates are bounded away from zero. If they are, the term vanishes at both ends and rises and falls in between. If one eigenvalue is near zero, the term instead rises and saturates, since that mode stays frozen while its partner decays. Only one eigenvalue has this property: the angular velocity diffusion eigenvalue of the REM stage, since *D* is driven to zero only when *β* is large and *σ*_*ω*_ is small (the opposite is true in Wake and SWS). Therefore, the REM-Wake distance will rise and saturate, while the Wake-SWS distance will have the more generic shape as described above. We further quantify this below in the setting where the difference is dominated by one pairing of eigenvalues. However, this generic shape result does not depend on which eigenvalue is compared with which.

Wake and SWS differ in eigenvalues that are all bounded away from zero: their radial rates differ most strongly (*µ*_Wake_ ≫ *µ*_SWS_), although their angular velocity and diffusion rates differ as well. The distance between Wake and SWS is dominated by the difference in their radial decay rates. Up to scaling constants, this distance’s behavior can be described as approximately

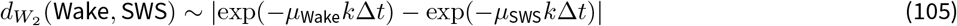

Given an initial mismatch (*µ*_Wake_ > *µ*_SWS_), this distance starts small, peaks early, and decays rapidly toward zero as both transient terms vanish. In practice the asymptote is bounded below by noise in the estimated eigenvalues.

Wake and REM instead have comparable radial rates, so this term is small at every timescale. Their distance is predominantly governed by the effective angular diffusion constant 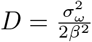.

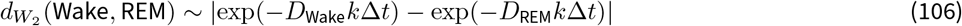

Because *β*_REM_ is large and 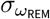 is small, *D*_REM_ ~ 0, yielding the approximation:

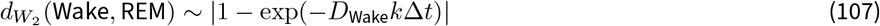

This characteristic curve starts small, grows quickly, and saturates asymptotically away from zero.

## J Koopman eigenvalues between coupled subsystems

Two dynamical systems can be coupled in a time-varying manner, and it can be informative to understand when this occurs. Here we study the relationship between the Koopman eigenvalues of coupled subsystems, to motivate the use of DSA in measuring coupling. The generic expression of coupled systems is:

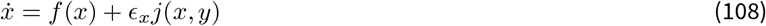

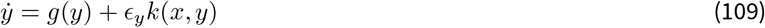

where *ϵ*_*x*_, *ϵ*_*y*_ are scalars that measure the effect of coupling. Note that both subsystems *x, y* may be multidimensional. We consider the setting of comparing observables *x, y* with DSA. Therefore, we may seek to understand how the Koopman Operators on observables of *x* or *y* alone relate. There are typically two settings that are of interest: the bidirectional case, in which *ϵ*_*x*_, *ϵ*_*y*_≠ 0, and the unidirectional case, in which *ϵ*_*x*_ = 0, *ϵ*_*y*_≠ 0 without loss of generality.

The bidirectional case is straightforward, as this can rather be considered a fully autonomous system made up of partial observations. In this setting, delay embedded observables on either *x* or *y* reconstruct the full state up to conjugacy, which preserves Koopman eigenvalues (*Takens, 1981; Budišić et al., 2012*). Therefore in this setting, their eigenvalues are shared.

The unidirectional setting is straightforward as well. In this setting, the subsystem *x* is a dynamical system in its own right, with its own Koopman eigenvalues Λ_*x*_. Delay-embedded observables on *y* reconstruct the full state including *x* (*Stark, 1999*), so the spectrum recovered by observables on *y* is the full spectrum Λ_*x,y*_ ⊇ Λ_*x*_ (*Schlosser and Korda, 2022*). Containment in this fashion is one-directional. Therefore, comparing distributions of eigenvalues is agnostic to the direction, but studying which set of observables’ eigenvalues contains the other’s can give a clue towards directionality.

